# *Drosophila* functional screening of *de novo* variants in autism uncovers deleterious variants and facilitates discovery of rare neurodevelopmental diseases

**DOI:** 10.1101/2020.12.30.424813

**Authors:** Paul C Marcogliese, Samantha L Deal, Jonathan Andrews, J Michael Harnish, V Hemanjani Bhavana, Hillary K Graves, Sharayu Jangam, Xi Luo, Ning Liu, Danqing Bei, Yu-Hsin Chao, Brooke Hull, Pei-Tseng Lee, Hongling Pan, Colleen M Longley, Hsiao-Tuan Chao, Hyunglok Chung, Nele A Haelterman, Oguz Kanca, Sathiya N Manivannan, Linda Z Rossetti, Amanda Gerard, Eva Maria Christina Schwaibold, Renzo Guerrini, Annalisa Vetro, Eleina England, Chaya N Murali, Tahsin Stefan Barakat, Marieke F van Dooren, Martina Wilke, Marjon van Slegtenhorst, Gaetan Lesca, Isabelle Sabatier, Nicolas Chatron, Catherine A Brownstein, Jill A Madden, Pankaj B Agrawal, Roberto Keller, Lisa Pavinato, Alfredo Brusco, Jill A Rosenfeld, Ronit Marom, Michael F Wangler, Shinya Yamamoto

## Abstract

Individuals with autism spectrum disorders (ASD) exhibit an increased burden of *de novo* variants in a broadening range of genes. We functionally tested the effects of ASD missense variants using *Drosophila* through ‘humanization’ rescue and overexpression-based strategies. We studied 79 ASD variants in 74 genes identified in the Simons Simplex Collection and found 38% of them caused functional alterations. Moreover, we identified *GLRA2* as the cause of a spectrum of neurodevelopmental phenotypes beyond ASD in eight previously undiagnosed subjects. Functional characterization of variants in ASD candidate genes point to conserved neurobiological mechanisms and facilitates gene discovery for rare neurodevelopmental diseases.

## Introduction

Autism spectrum disorder (ASD) is a complex neurodevelopmental condition with impairments in social interaction, communication and restricted interests or repetitive behaviors^1^. Individuals affected by ASD, particularly in severe cases, exhibit a higher burden of *de novo* mutations (DNMs) in an expanding list of genes^2–4^. The genetic burden of DNMs in ASD patients has been estimated to account for ~30% of disease causation^4–9^. While these studies have implicated hundreds of genes in ASD pathogenesis, which of these genes and variants causally contribute to this disease remains unknown. Missense DNMs in particular present a unique challenge because most genes lack established functional assays. *Drosophila melanogaster* is a genetically tractable system that is widely used to study human diseases^10–12^. In addition to studying disease mechanisms by establishing preclinical models, flies can be used as a ‘living test tube’ to study functional consequences of variants of unknown significance found in patients. Here, we integrate a number of state-of-the-art technologies in the fly field to establish an *in vivo* pipeline to effectively study the functional impact of DNMs identified in a large cohort of ASD patients.

## Results

### Prioritization of ASD variants to study in *Drosophila*

We prioritized genes with coding DNMs identified in ASD probands from the Simons Simplex Collection (SSC)^4^ that are conserved in *Drosophila*. We primarily focused on missense variants and in-frame indels because functional consequences of these variants are more difficult to predict compared to nonsense and frameshift alleles (Figure 1, Table S1). We also tested a few truncating variants in single-exon genes because these transcripts escape nonsense mediated decay. We then selected variants in genes that correspond to fly genes with intronic MiMIC (*Minos*-mediated integration cassette) elements, a versatile transposon that allows sophisticated genomic manipulations^13,14^. By converting the original MiMICs into T2A-GAL4 (TG4) lines via recombinase-mediated cassette exchange^15–17^, we generated a collection of fly lines that behaves as loss-of-function (LoF) alleles that simultaneously produce a GAL4 transactivator in the same temporal and spatial pattern as the gene of interest^18^. Using this strategy, we successfully generated 109 TG4 lines that correspond to 128 SSC genes (some fly genes correspond to multiple human genes with variants in SSC). In parallel, we also generated 106 UAS human reference transgenic (Ref-Tg) and 88 SSC-DNM transgenic (SSC-Tg) strains (Figures 1E and 1G, Tables S2 and S3). These lines can be used in combination with TG4 lines to ‘humanize’ *Drosophila* genes, or can be crossed to ubiquitous or tissue-specific drivers to ectopically overexpress reference or variant human proteins^10^ (Figure 1H).

**Figure 1:**
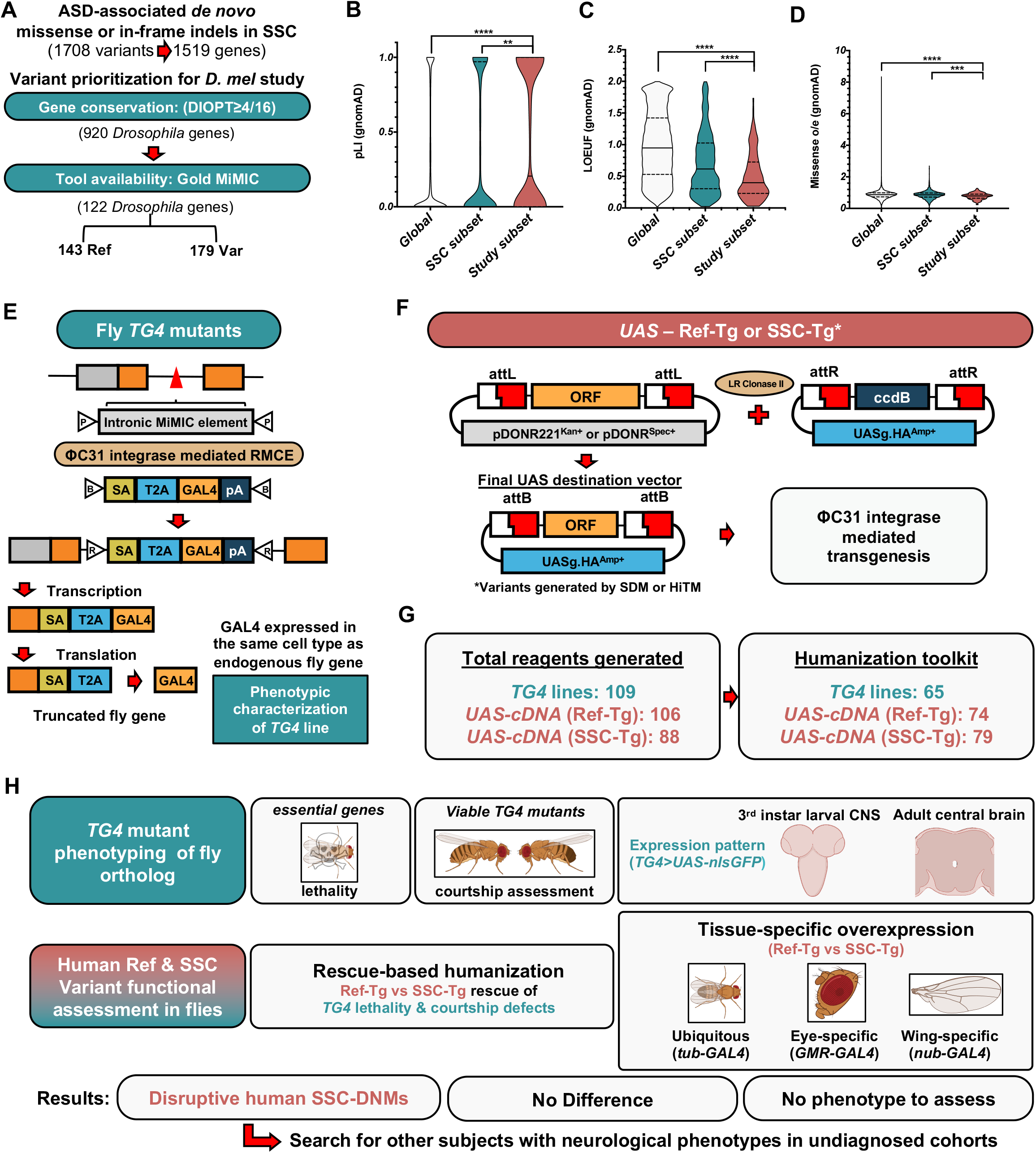
Gene & variant prioritization, resource generation and screening outline. (A) Criteria to prioritize ASD candidate genes and variants for this study. (B-D) Gene level constraints from control individuals (gnomAD). (E) Schematic depicting generation and effect of *TG4* lines on gene function. (F) Schematic illustrating generation of UAS-human cDNA constructs. (G) Total number of *Drosophila* reagents generated for this study. (H) Screening paradigms using both humanization and overexpression strategies to assess SSC-DNM function.

### Humanization of essential *Drosophila* gene reveals loss-of-function ASD variants

We identified 47 lethal TG4 mutants that correspond to 60 human ASD candidate genes for which both reference and variant human cDNA transgenic fly lines were successfully established (Figure S1, Table S4). To assess whether the human homolog can replace the corresponding fly genes, we determined whether UAS-Ref-Tg can rescue the lethality of lethal TG4 mutants. We assessed rescue at four temperatures (18°C, 22°C, 25°C, 29°C) as the GAL4/UAS system is temperature dependent^14^. We found that lethality was suppressed in 17 of 37 genes tested (46%; Figure 2A, Table S5). We next tested whether SSC-DNMs have functional consequences by comparing the rescue efficiency of UAS-Ref-Tg and UAS-SSC-Tg. We observed significant functional differences in the ability for SSC variants to rescue lethality for *ABL2*, *CAT*, *CHST2*, *TRPM6* (2 variants), and *TRIP12* (Figures 2B-2D). For *ABL2* and *CAT*, we further found that humanized flies carrying the SSC-DNM (*ABL2*^*A1099T*^ or *CAT*^*G204E*^) had significantly decreased lifespan compared to reference animals (Figures 2E and 2F). Overall, we found that 32% (6/19) of SSC-DNMs functionally differed from the reference *in vivo*, all behaving as LoF alleles.

**Figure 2:**
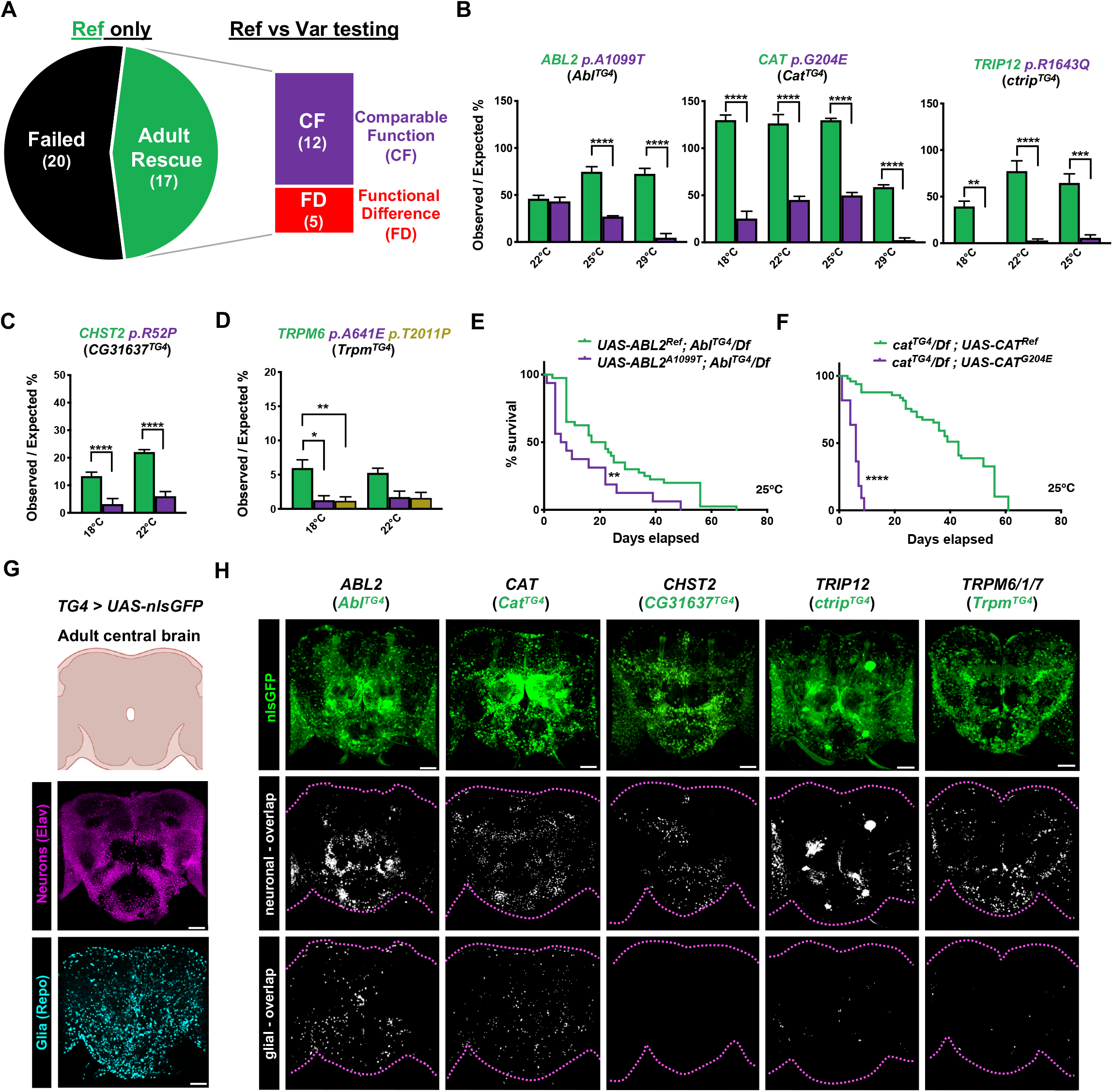
Assessment of SSC-DNM function through humanization of essential fly genes. (A) Rescue of lethality to adult stage by *TG4* driven UAS-reference human cDNA and subsequent comparison of reference and variant cDNA. (B-D) Observed/expected Mendelian ratios for rescue of humanized *TG4* mutants across different temperatures. 3 independent crosses where set per genotype and n>50 flies were quantified for each cross. Statistical analyses were performed by ANOVA followed by Sidak’s multiple comparisons test. (E-F) Lifespan analysis of humanized *TG4* lines at 25°C. Survival comparisons obtained by Log-rank (Mantel-Cox) test. (G-H) Cartoon and confocal images showing expression pattern of UAS-nlsGFP driven by *TG4* (green), and co-staining corresponding to neurons (Elav, magenta) and glia (Repo, cyan). Colocalization of GFP and each cell specific marker (white). Scale bar = 25 μm. Dotted magenta lines outline of the brain. *p<0.05, **p<0.01, ***p<0.001, ****p<0.0001.

To assess whether the fly homologs of human ASD candidate genes from SSC are expressed in the central nervous system (CNS), we crossed each TG4 line to UAS-nlsGFP (green fluorescent protein with a nuclear localization signal) and performed co-staining with neuronal (Elav) and glial (Repo) nuclear markers (Figure 2G). All five genes associated with deleterious LoF DNMs were expressed in the adult brain (Figure 2H) as well as in the developing (third instar larval) CNS (Figure S2). All five genes were found predominantly in neurons in adult brains. In the larval CNS, four genes were also enriched in neurons but *Cat* (corresponding to human *CAT*) was found to be enriched in glia.

### Loss- and Gain-of-Function ASD variants identified through behavior analysis on humanized flies for viable TG4 lines

While we were able to test the function of 37 human genes based on rescue of lethality, 62 TG4 lines corresponding to 70 SSC-ASD candidate genes were viable and did not exhibit any obvious morphological phenotypes. To determine if any of these viable mutants display a behavioral phenotype that can be used for variant function assessment, we performed courtship assays^21,22^ (Figure 3A). Courtship involves a complex set of neurological components involving sensory input, processing and motor output^22^. We measured the amount of time a male fly spent performing wing extensions (courtship) and copulating towards a wild-type (*Canton-S*) female. We also measured the time flies spent moving within the test chamber to assess their locomotion. Finally, we also tracked grooming, a stereotypic behavior in flies that involves a complex neurocircuit^23^. Of 21 viable TG4 lines analyzed, we found that 15 display recognizable behavioral alterations (Figures 3C-3F and S3A-S3D).

**Figure 3:**
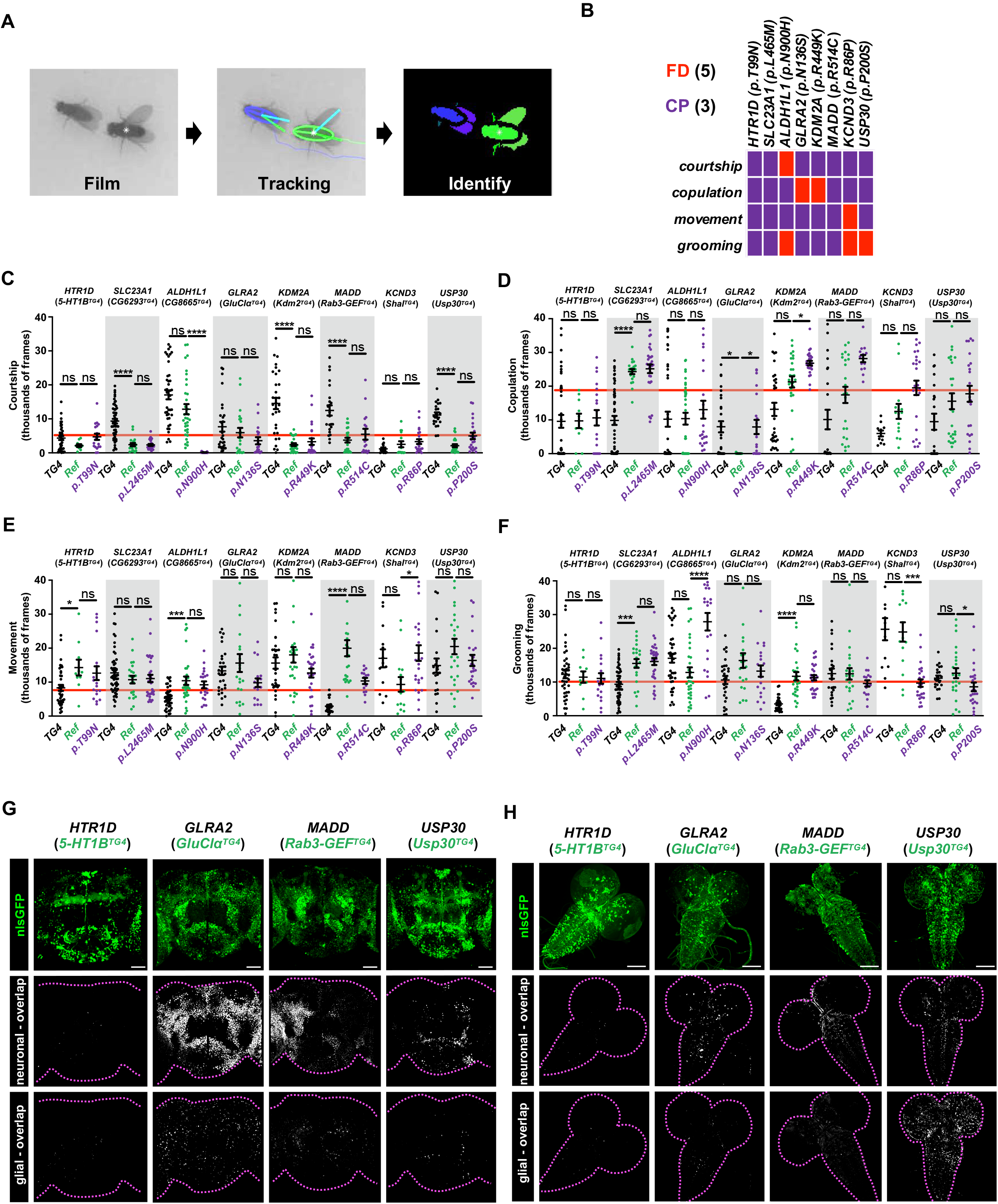
Assessment of SSC-DNM function through humanization of viable TG4 lines and behavioral analysis. (A) Analysis pipeline used to evaluate *Drosophila* behavior. (B) SSC-DNMs in which variants display significant differences in time spent performing a specific behavior (courtship, copulation, movement, or grooming) when compared to reference humanized flies. (C-F) The number of frames male flies spent performing courtship (single-wing extensions), copulating, moving within the chamber, or grooming during a 30-minute test period. The red line represents the average number of frames a *Canton-S* (control) male spends performing the same task. n=10-40 flies were used per genotype. Statistical analysis performed by Kruskal-Wallis one-way analysis of variance and the Dunn’s multiple comparison test. (G-H) Representative images demonstrating the expression pattern of UAS-nlsGFPdriven by *TG4* in the adult or third instar larval brain co-stained with neuronal (Elav) and glial (Repo) markers. Scale bar = 25 μm. White signal in the two bottom rows display overlap between nlsGFP and the neuronal or glial marker. *p<0.05, ***p<0.001, ****p<0.0001, ns (not significant).

Of 15 viable TG4 lines with quantitative behavior defects, we were able to humanize eight of them using Ref-Tgs and SSC-Tg to assess variant function. We found five SSC-DNMs that showed functional alterations from the reference allele in at least one of four behavioral paradigms (Figure 3B). Two variants (*GLRA2*^*N136S*^ and *KCND3*^*R86P*^) behaved as LoF alleles. Humanized reference *GLRA2* flies failed to copulate at all, while the humanized *GLRA2*^*N136S*^ flies were capable of copulating within the trial period similar to the TG4 mutant alone (Figure 3D). Humanized *KCND3*^*R86P*^ flies displayed increased movement and decreased grooming behavior when compared to the humanized reference flies (Figures 3E and 3F). Interestingly, three variants (*KDM2A*^*R449K*^, *ALDH1L1*^*N900H*^ and *USP30*^*P200S*^) behaved as gain-of-function (GoF) alleles. While humanized *KDM2A* reference flies had a trend for increased time spent copulating compared to TG4 mutant alone, *KDM2A*^*R449K*^ flies showed significant increase in time spend copulating (Figure 3D). Humanized *ALDH1L1*^*N900H*^ flies displayed a significant reduction in courtship and an increase in grooming behavior when compared to the humanized reference fly or the TG4 mutant alone (Figures 3C and 3F). Humanized *USP30*^*P200S*^ flies displayed decreased grooming behavior when compared to humanized reference flies (Figure 3F).

Finally, we determined the CNS expression of TG4 lines corresponding to all eight lines we were able to humanize. Surprisingly, we only detected expression of 4/8 genes in the larval and adult CNS (Figures 3G and 3H). In sum, over half (15/21) of the viable TG4 mutants assayed show some behavioral alteration compared to a commonly used control (*Canton-S)* strain (shown in red lines in Figures 3C-3F and S3A-S3D), and 63% (5/8) SSC-DNMs act functionally different from reference alleles using quantitative behavioral measurements in flies: two behaved as LoF alleles whereas three behaved as GoF alleles. This is in contrast to the humanization rescue-based functional studies performed on essential gene homologs that only revealed LoF alleles.

### Overexpression assays revealed ASD variants with diverse functional consequences in genes that correspond to viable TG4 lines

We complemented our rescue-based assays by overexpressing Ref-Tg and SSC-Tg in a wild-type background using ubiquitous (*tub-GAL4*), eye-specific (*GMR-GAL4*), and wing-specific (*nub-GAL4*) drivers (Figure 4). Critically, UAS Ref-Tg and SSC-Tg are inserted into the same genomic landing site in the fly genome to allow for direct functional comparison. Across the three drivers and testing 66 human genes (73 SSC variants), we found 21 SSC-DNMs (in 19 fly genes) display altered function, 17 displayed phenotypes comparable to reference and 35 did not produce a scorable phenotype (Table S5).

**Figure 4:**
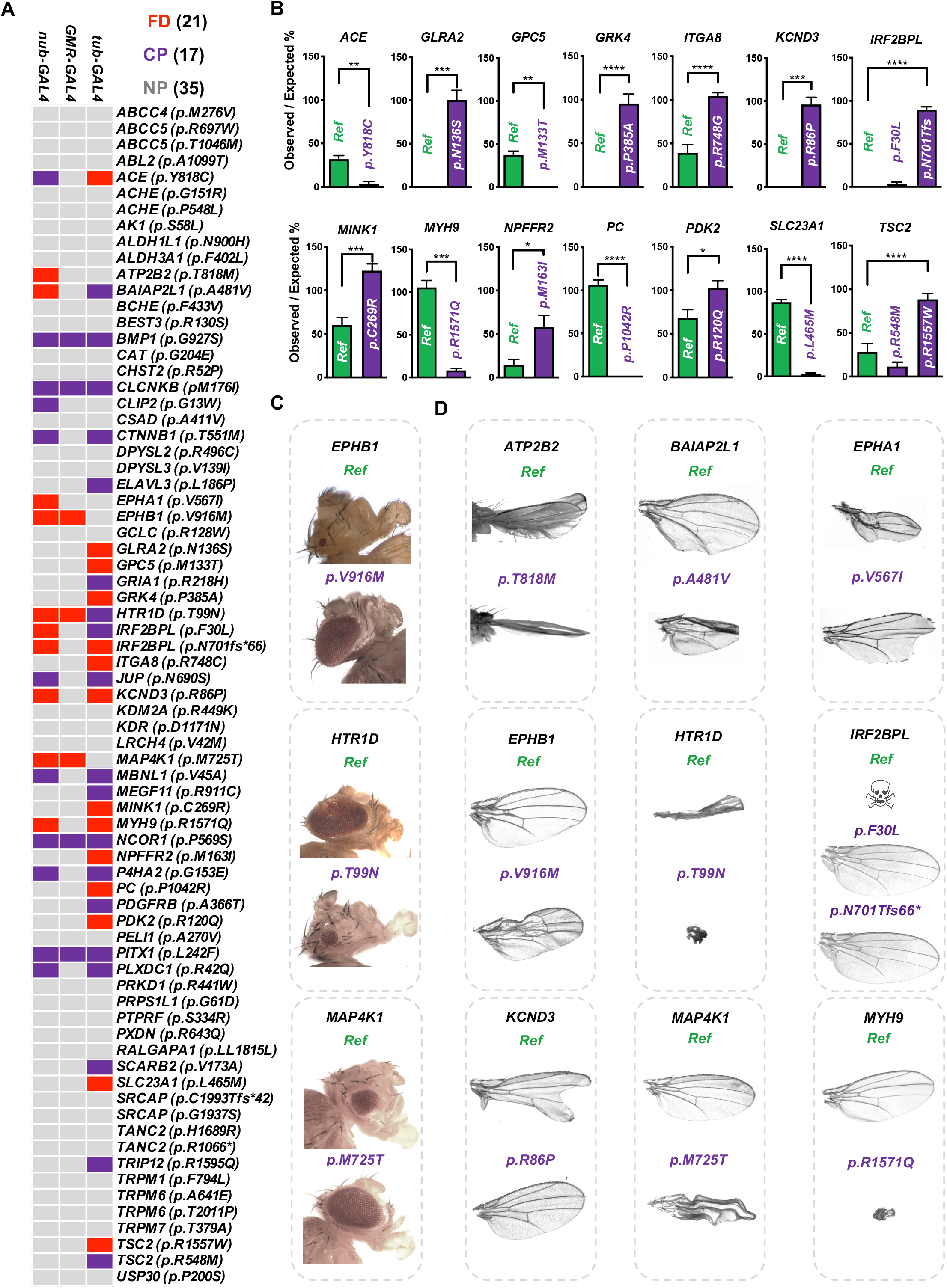
Variant assessment by overexpression of reference and SSC-DNMs. (A) Phenotypes observed upon overexpressing the reference and variant cDNAs using a ubiquitous driver (*tub-GAL4*) at 25°C, an eye-specific driver (*GMR-GAL4*) at 29°C, or a wing-specific driver (*nub-GAL4*) at 25°C. Black denotes if there was no phenotype (NP), purple if there was a comparable phenotype (CP), or red if there was a functional difference (FD). (B) Quantification of viability upon overexpression of reference or variant human cDNAs using a ubiquitous driver for genes where the variants showed a functional difference. Minimum of 3 independent crosses were set with two independent UAS-transgenic lines. 50-100 flies (a minimum of 10 if overexpression caused survival defects) were scored. Statistical analyses were performed by unpaired t-test. (C-D) Representation of optical sections of eyes and wings for variants with a functional difference using eye-specific (*GMR-GAL4*) and wing-specific (*nub-GAL4*) *drivers*, respectively at 25°C. *p<0.05, **p<0.01, ***p<0.001, ****p<0.0001.

Twelve variants in eleven human genes (*ATP2B2*^*T818M*^, *EPHA1*^*V567I*^, *GLRA2*^*N136S*^, *GRK4*^*P385A*^, *ITGA8*^*R748G*^, *IRF2BPL*^*F30L*^, *IRF2BPL*^*N701fs66\**^, *KCND3*^*R86P*^, *MINK1*^*C269R*^, *NPFFR2*^*M163I*^, *PDK2*^*R120Q*^, *TSC2*^*R1557W*^) behaved as LoF alleles. *GLRA2*^*N136S*^, *GRK4*^*P385A*^, *ITGA8*^*R748G*^, *KCND3*^*R86P*^, *MINK1*^*C269R*^, *NPFFR2*^*M163I*^, *PDK2*^*R120Q*^, *TSC2*^*R1557W*^ were annotated as LoF alleles using a ubiquitous driver because they failed to reduce the expected viability to the extent of the corresponding reference alleles upon overexpression (Figure 4B). Notably, *GLRA2*^*N136S*^ and *KCND3*^*R86P*^ were also annotated as LoF alleles in our rescued based assay, showing consistency (Figure 3). Moreover, *KCND3*^*R86P*^ and *IRF2BPL*^*N701fs66\**^ variants behaved as LoF alleles when assessed with multiple drivers. *KCND3*^*R86P*^ abolished the activity of the reference transgene to cause lethality when overexpressed with a ubiquitous driver (Figure 5B). In the wing, *KCND3*^*R86P*^ failed to produce a severe notching phenotype that is observed by expression of the reference transgene (Figure 5D). Ubiquitous or wing-specific overexpression of reference *IRF2BPL* caused lethality, whereas the *IRF2BPL*^*N701Tfs66\**^ frameshift allele (note that *IRF2BPL* is a single-exon gene) does not cause any phenotype (Figures 4B and 4D). Interestingly the missense variant, *IRF2BPL*^*F30L*^ behaved as a LoF using the wing driver but was indistinguishable using the ubiquitous driver, indicating it is likely to be a partial LoF allele (Figures 4B and 4D). In addition to *IRF2BPL* DNMs, variants in two genes (*ATP2B2 and EPHA1*) were identified as LoF alleles based on the wing-specific driver and assay. Wing-specific overexpression of reference *ATP2B2* in the developing wing disc causes a curled wing phenotype while the *ATP2B2*^*T818M*^ variant fails to do so (Figure 4D). Expression of reference *EPHA1* caused a wing size reduction and wing margin serration, whereas the *EPHA1*^*V567I*^ variant caused serrated wings of normal size (Figure 4D) indicating partial LoF.

**Figure 5:**
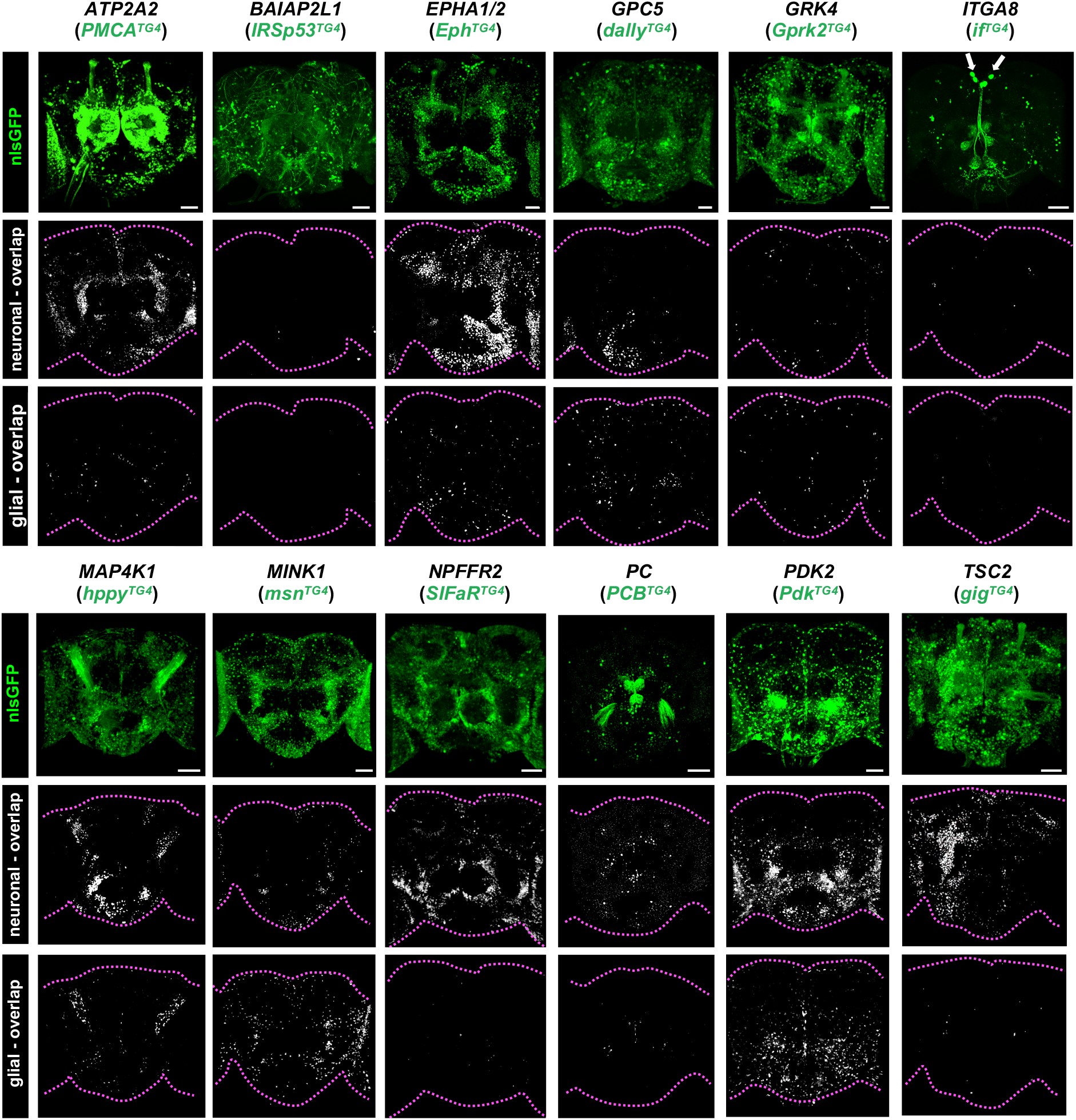
Most genes with SSC variants showing functional difference based on overexpression screening are also expressed in the *Drosophila* CNS. Representative images demonstrating the expression pattern of UAS-nlsGFP driven by *TG4* in the adult brain co-stained with neuronal (Elav) and glial (Repo) markers. White signal in the bottom two rows display overlap between GFP and the neuronal or glial marker. Scale bar = 25 μm. Dotted magenta lines outline the brain.

Seven variants (*ACE*^*Y818C*^, *GPC5*^*M133T*^, *MYH9*^*R1571Q*^, *PC*^*P1024R*^, *SLC23A1*^*L465M*^, *HTR1D*^*T99N*^, *BAIAP2L1*^*A481V*^) behaved as GoF alleles. Flies overexpressing variant forms of *ACE*, *GPC5*, *MYH9*, *PC*, and *SLC23A1* exhibited enhanced lethality when compared to reference protein (Figure 4B). The GoF nature of *MYH9*^*R1571Q*^ was also observed in a wing size-based assay (Figure 4D). *HTR1D*^*T99N*^ displayed consistent stronger phenotypes compared to reference when expressed in the eye or the wing, resulting in eye size reduction and absent wing phenotype, respectively (Figures 4C and 4D). *BAIAP2L1*^*A481V*^ caused a smaller, more crumpled wing phenotype compared to its reference allele (Figure 4D).

Intriguingly, *EPHB1*^*V916M*^ and *MAP4K1*^*M725T*^ exhibited conflicting results in the eye and wing, therefore they could not be categorized as simple LoF or GoF variants. While overexpression of reference *EPHB1* or *MAP4K1* in the developing eye causes eye size reduction phenotype, SSC variant forms of either gene result in normal eyes, indicating they behave as LoF alleles in this tissue. However, the same variant transgenes for these two genes expressed in the wing result in blistered or crumpled wings, respectively, that are phenotypically stronger than the reference alleles (Figures 4C and 4D), indicating they behave as GoF alleles in this tissue. In summary, when a scorable phenotype was present, 48% (21/44) of the SSC-DNMs tested with an overexpression strategy impact function. Furthermore, we found diverse SSC variant consequences including twelve LoF, seven GoF, and two with complex phenotypes.

While this overexpression based screening approach was not directly investigating the function of genes in the nervous-system, expression analysis revealed that most (15/19) fly genes that correspond SSCs with a functional difference identified through our overexpression assay are expressed in the adult brain and larval CNS [Figure 5, see ^19^ for *Pits* (corresponding to human *IRF2BPL*)]. While most genes are enriched in neuronal subpopulations, *msn* (*MINK1*) is enriched in glia. Interestingly, *if* (*ITGA8*) is not detected in either neurons or glia but revealed a pattern reminiscent of cells in *pars intercerebralis*, a neuroendocrine organ analogous to the mammalian hypothalamus^20^ (Figures 5 arrow and S4). Furthermore, *Eph* (*EPHA1*, *EPHA2*), was found to be enriched in glia in the larval CNS but is primarily neuronal in the adult brain (Figures 5 and S4). Taken together, most of the fly genes corresponding to SSC-DNMs we found an in vivo alteration in function using overexpression-based assays are expressed in the fly CNS, similar to hits from rescue-based studies.

### Thirty deleterious SSC variants identified by merging all functional data

In total, we found 29 missense and 1 frameshift SSC-DNMs displaying functional differences when compared to their respective reference allele (1 variant for 26 genes, 2 variants for 2 genes) (Tables 1 and S5). Approximately 53% (30/57) of the SSC-DNMs exhibited functional differences compared to the reference. Intriguingly in our study, we only found GoF variants for genes corresponding to viable TG4 fly mutants based on both rescue-based and overexpression-based assays (Figure S5A).

**Table 1:**
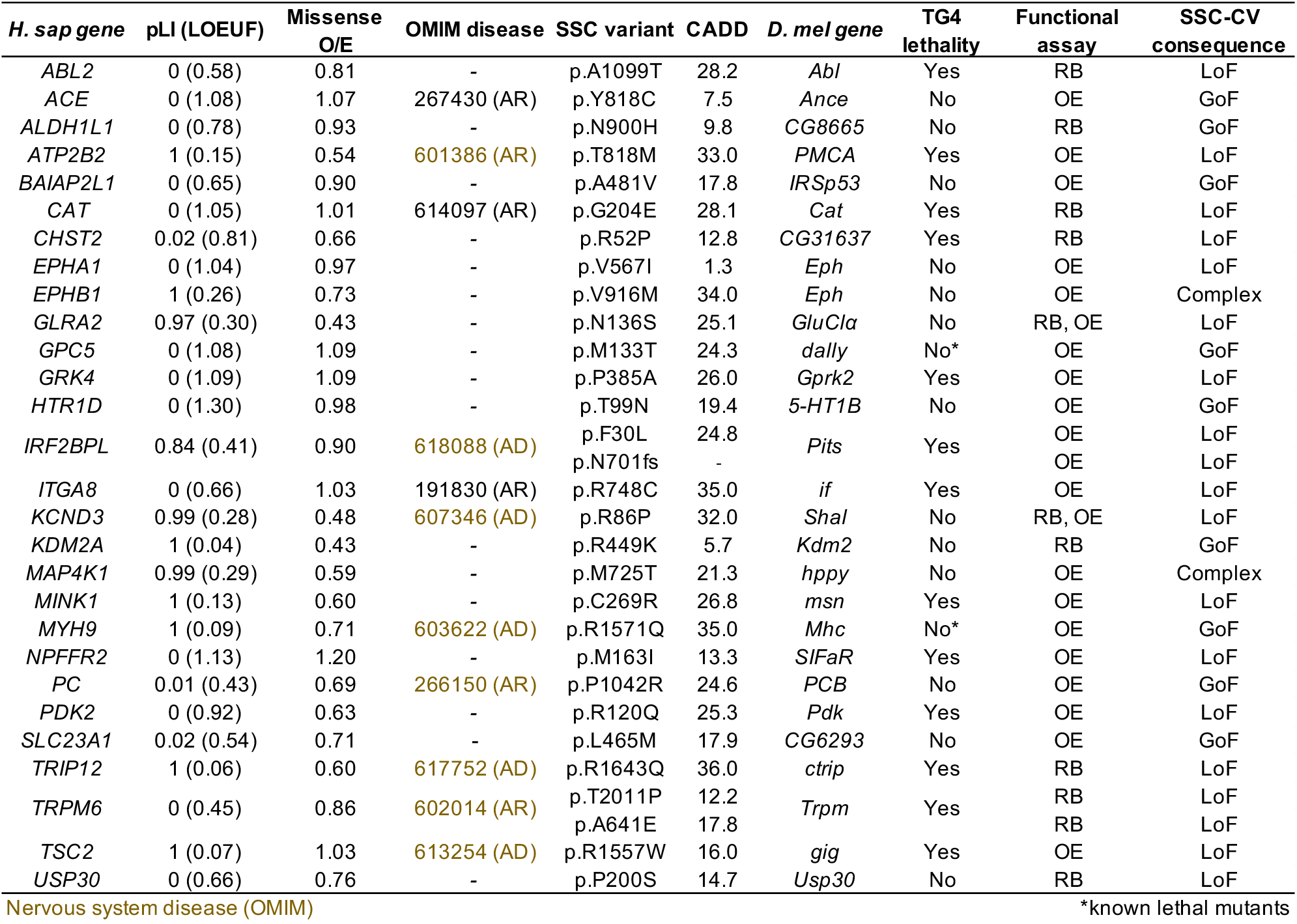
Identification of 30 SSC-DNMs with functional consequences. List of all human genes and corresponding SSC variants determined to have a functional difference across all assays in this study. RB=rescue-based, OE=overexpression, LoF=Loss-of-Function, GoF=Gain-of-Function.

When we informatically surveyed the genes and variants with functional consequences identified through our screen in comparison to other genes included in our study (variants with comparable function or those lacking a phenotype to assess) using the MARRVEL tool^24^, we did not find any significant differences in gene level metrics such as pLI (probability of Loss of function Intolerance)^25^, LOEUF [Loss of function observed/expected (o/e) upper bound fraction]^26^, missense o/e (observed/expected)^26^, pathogenicity prediction scores based on several *in silico* algorithms including SIFT^27^, PolyPhen-2^28^, and CADD^29^, or absence/presence of identical variant in gnomAD^25^ (Tables S5 and S6; Figures S5B-Z). By analyzing Gene Ontology (GO) by PantherDB^30^, ASD candidate genes from SSC with deleterious variants *in vivo* were compared to all protein coding genes. We found significant enrichment for genes with GO terms for ‘synapse (cellular component)’ and ‘ATP binding (function)’ (Figure S6). Finally, we systematically imaged the expression pattern of 41 TG4 lines generated through our study to document their expression in the adult and larval CNS in neurons or glia as a resource for the community (Figures S7 and S8).

### Rare LoF and GoF variants in *GLRA2* cause X-linked neurodevelopmental disorders with overlapping phenotypes in males and females, respectively

Many genes implicated in ASD are associated with neurodevelopmental disorders beyond autism^31,32^. Therefore we asked if genes with disruptive SSC-DNMs could be used to discover new disease-causing genes and disorders by identifying patients with rare potentially deleterious variants that have been unrecognized^33–35^. Out of 28 genes in which we identified damaging SSC-DNMs, eight are associated with Mendelian diseases with neurological presentations in OMIM^36^ (Table 1 and SI Case Histories). For one of these genes (*IRF2BPL*), we recently reported *de novo* truncating variants in *IRF2BPL* as the cause of a novel severe neurodevelopmental disorder with abnormal movements, loss of speech and seizures in collaboration with the Undiagnosed Diseases Network^19,35^. Here, we report identification of *GLRA2* as a cause of novel neurodevelopmental syndromes with overlapping features such as developmental delay (DD), intellectual disability (ID), ASD and epilepsy identified through re-analysis of clinical exome sequencing data and matchmaking through GeneMatcher^33^ (Figure 1H).

We identified rare *GLRA2* variants in eight unrelated subjects with or without autistic features. In addition to developmental and cognitive delay of variable severity, 3/8 subjects have microcephaly, 4/8 subjects have a history of epilepsy and 6/8 subjects have ocular manifestations, including congenital nystagmus that improved with age in three of them (Table 2). *GLRA2* (*Glycine Receptor Alpha 2*) is an X-linked gene that encodes a subunit of a glycine-gated chloride channel^37^. All five female subjects harbored DNMs including a recurring *GLRA2*^*T296M*^ variant found in 4/5 female subjects. This variant was also identified in a female subject in previous large-scale developmental disorder study^38^. The three male subjects had maternally inherited variants, and the mothers are not symptomatic. The CADD scores for all three male subjects are predicted to be damaging (Table 2). A maternally-inherited microdeletion of *GLRA2* has also been found in a single male patient with ASD^39^, indicating that hemizygous LoF allele of this gene in males may cause ASD. Indeed, we determined that the *GLRA2*^*N136S*^ variant present in the SSC in a male subject acts as a LoF allele based on our behavioral assay on humanized flies (Figure 3D) as well as through overexpression studies (Figure 4B).

**Table 2:** Salient features of subjects with *GLRA2* variants. Abbreviations are as follows: MRI, magnetic resonance imaging; EEG=electroencephalography, CADD=Combined Annotation Dependent Depletion.

| Subject | 1 | 2 | 3 | 4 | 5 | 6 | 7 | 8 |
| --- | --- | --- | --- | --- | --- | --- | --- | --- |
| **GLRA2 Variant**<br>(hg19, NM_001118886.2) | c.887C>T,<br>p.Thr296Met | c.887C>T,<br>p.Thr296Met | c.887C>T,<br>p.Thr296Met | c.887C>T,<br>p.Thr296Met | c.140T>C,<br>p.Phe47Ser | c.754C>T,<br>p.Arg252Cys | c.862G>A, p.<br>Ala288Thr | c.1186C>A, p.<br>Pro396Thr |
| **Inheritance** | *De novo* | *De novo* | *De novo* | *De novo* | *De Novo* | Maternal | Maternal | Maternal |
| **CADD Score** | 27 | 27 | 27 | 27 | 27.8 | 31 | 27.2 | 20.9 |
| **Gender** | Female | Female | Female | Female | Female | Male | Male | Male |
| **Age at most recent**<br>**evaluation (years)** | 6.7 | 6.5 | 5.5 | 0.5 | 6.7 | 0.9 | 7 | 34 |
| **Developmental**<br>**delay/intellectual**<br>**disability** | Yes | Yes | Yes | Yes | Yes | Yes | Yes, with<br>regression | Yes |
| **Hypotonia/incoordination** | No | No | Yes, ataxic gait | Yes | No | Yes | Yes, ataxia | Yes |
| **Autism spectrum**<br>**disorder** | No | Yes | No | N/A | No | N/A | Yes | Yes |
| **Inattention/hyperactivity** | Yes | Yes | No | N/A | Yes | N/A | No | No |
| **Sleep disturbance** | No | Yes | No | No | Yes | Yes | No | No |
| **Microcephaly** | No | Yes | Yes | Yes | No | No | No | No |
| **Ocular features** | Myopia,<br>astigmatism and<br>nystagmus<br>(improved with<br>age) | Nystagmus<br>(improved with<br>age) | Alternating<br>exotropia,<br>borderline<br>opsoclonus | None | Strabismus,<br>nystagmus<br>(improved with<br>age) | Strabismus | Myopia | Myopia,<br>astigmatism |
| **Epilepsy** | No | Yes | No | Yes | Yes | No | Yes | No |
| **EEG findings** | Slow background<br>suggestive of mild<br>encephalopathy | Bilateral<br>synchronized high<br>amplitude spikes,<br>Epileptic potentials | Normal | Slow background,<br>infantile spasms,<br>multifocal spikes<br>during sleep | Infantile spasms,<br>then normal<br>interictal EEG | Not performed | Generalized<br>slowing and<br>generalized<br>epileptiform<br>discharges<br>associated with<br>myoclonic jerks | Not performed |
| **Brain MRI findings** | Normal | Delayed<br>myelination, a<br>small arachnoid<br>cyst | Mild cortical<br>atrophy, thinning<br>of corpus callosum | Normal | Cortical and white<br>matter atrophy,<br>including vermian<br>atrophy | Not performed | Minimally<br>increased T2<br>signal intensity on<br>the occipital lobes | Normal |

To further understand the functional consequences of variants found in our *GLRA2* cohort, we generated additional transgenic flies to assay the function of p.T296M (found in female subjects) and p.R252C (found in a male subject) variants (Figures 6A and S9A). By overexpressing reference and variant *GLRA2* using a ubiquitous driver, we found that *GLRA2*^*R252C*^ behaves as a LoF allele (Figure 6B), similar to *GLRA2*^*N136S*^ (Figure 4B). In contrast, this assay did not distinguish *GLRA2*^*T296M*^ from the reference (Figure 6B). Given the recurrent nature of this variant, as well as structural prediction that the residue has a potential role in obstruction of the ion pore in the closed conformation (Figure S9G)^40,41^, we further tested *GLRA2*^*T296M*^ and other alleles using additional GAL4 drivers. Using a *pnr-GAL4* that is expressed in the dorsocentral stripe in the notum, we found that *GLRA2*^*T296M*^, but not the reference or other *GLRA2* variant tested, causes lethality when expressed at high levels (Figure S9B). When we expressed *GLRA2*^*T296M*^ at lower levels by manipulating the temperature, we found that this variant induces the formation of melanized nodules in the thorax, a phenotype that we never observe when the reference or other *GLRA2* variants are overexpressed (Figures 6C, 6D and S9B).

**Figure 6:**
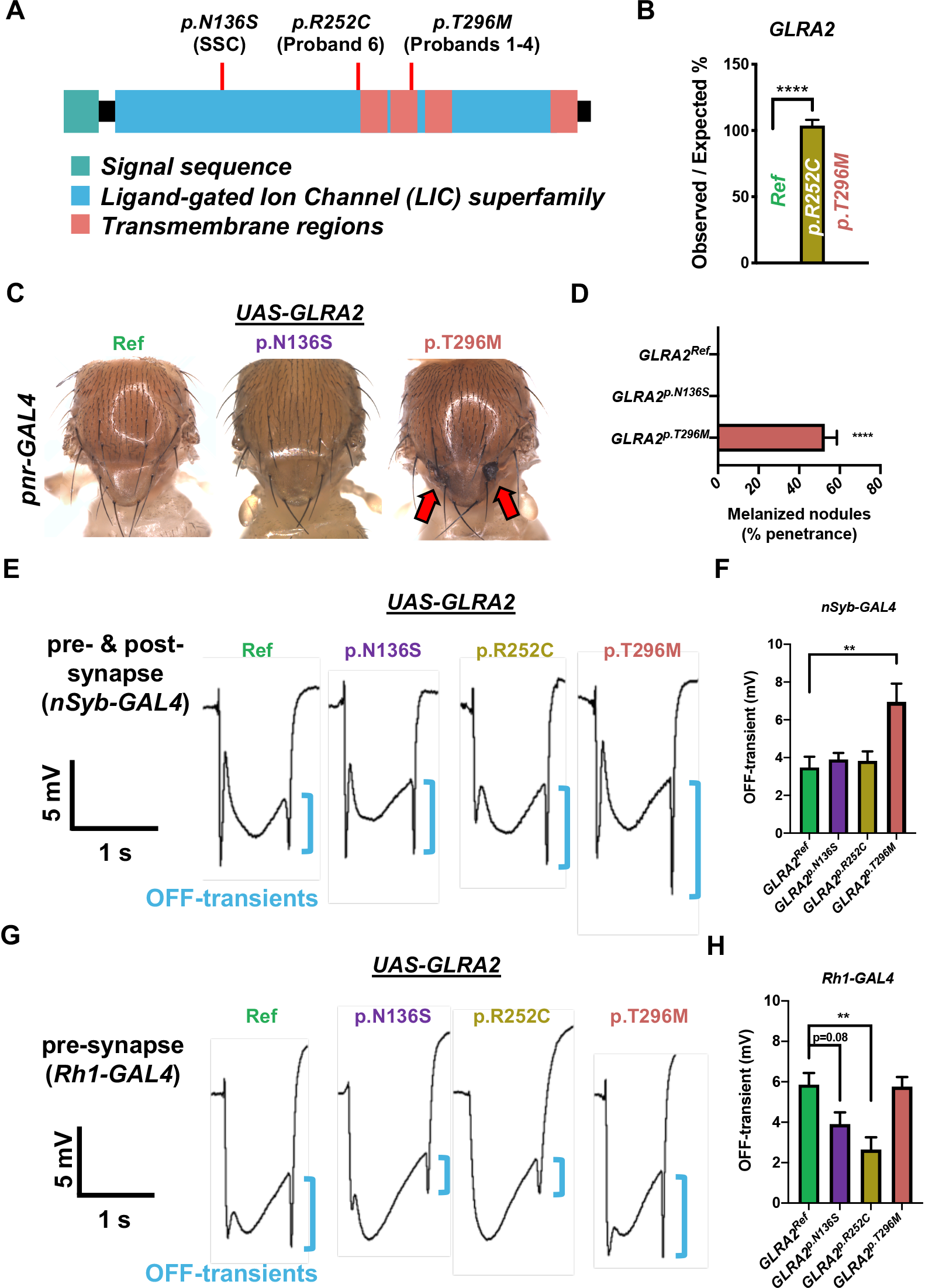
*GLRA2*^*T296M*^ found in female probands acts as a GoF allele while *GLRA2*^*R252C*^ and GLRA2^*N136S*^ found in male probands behave as LoF alleles. (A) Schematic diagram of domain structure of GLRA2 and the relative positions of subject variants functionally assessed in *Drosophila*. (B) Mendelian ratios upon overexpression the *GLRA2* reference or variant human cDNAs using a ubiquitous driver (*tub-GAL4*). (C-D) Representative images and quantification of melanized nodules formed on the notum of flies expressing *GLRA2*^*T296M*^ driven by a dorsocentral throax-specific (*pnr-GAL4*) driver at 25°C. (E-H) Representative traces of ERG and quantification of “*OFF*”-transient amplitude (blue bracket) in animals expressing GLRA2 panneuronally (both pre-synaptic photoreceptors and post-synaptic laminar neurons, *nSyb-GAL4*) or only in the pre-synaptic photoreceptors (*Rh1-GAL4*).

To further examine the functional consequences of overexpression of reference and variant *GLRA2* in the nervous system, we performed electroretinogram (ERG) recordings on the fly eye expressing human *GLRA2* using two distinct drivers. Panneuronal driver (*nSyb-GAL4*)^42^ allows one to express *GLRA2* in both pre-synaptic photoreceptors and post-synaptic neurons in the nervous system. Using this driver, we found a significant increase in amplitudes of “*OFF*” transients with *GLRA2*^*T296M*^ (Figures 6E, 6F, S9C, and S9D). This indicates an increase in synaptic transmission^43^, supporting the finding in the notum that p.T296M behaves as a GoF allele. Interestingly, when we limited the expression of *GLRA2* to pre-synaptic photoreceptors using *Rh1-GAL4*^44^, we did not observe any functional difference between *GLRA2*^*T296M*^ and the reference allele. However, with this driver, we were able to discern that both *GLRA2*^*R252C*^ seen in Subject 6 and *GLRA2*^*N136S*^ found in an SSC subject behave as LoF alleles based on observing a decrease in amplitude of *“OFF*” transients, indicating a decrease in synaptic transmission (Figures 6G, 6H, S9E, S9F). Hence, we have identified a cohort of subjects with deleterious variants in an X-linked gene *GLRA2* and shown that a recurrent missense DNM in females acts as a GoF allele, whereas alleles in male subjects behave as LoF alleles.

## Discussion

In this study, we generated >300 fly strains that allow functional studies of human variants and homologous fly genes *in vivo*. These reagents can be used to study many coding variants that are being identified through next-generation sequencing efforts in the human genomics field in diverse disease cohorts beyond ASD. Our screen elucidated 30 SSC variants with functional differences compared to reference, which was over half (~53%) of the genes in which we were able to perform a comparative functional assay.

Our screen was part of a larger effort to characterize the functional consequences of missense *de novo* changes from the SSC dataset using different strategies. One approach was based on proteomics by performing yeast-two-hybrid assays on 109 SSC-DNMs found in patients, showing 20% of protein-protein interactions that are found in reference proteins are disrupted by variants^45^. Another group reported that ~70% of 37 SSC-DNMs knocked-in to homologous *C. elegans* genes caused scorable phenotypes^46^. These studies are complementary to each other because while some variants have been identified as deleterious by more than one approach (e.g. *GLRA2*^*N136S*^ identified in both worm and fly screens), others are uniquely identified in one study, some of which could be due to technical limitations. For example, our approach of utilizing human cDNA transgenes allowed us to test variant function regardless of residue conservation in *Drosophila*. Of the 29 disruptive missense SSC-DNMs identified through our study, 14 affect residues that are conserved in flies and 10 in worms.

To take a rather unbiased approach, our gene prioritization was only based on gene-level conservation and tool availability (e.g. intronic MiMIC lines, full length human cDNA) rather than based on gene level constraints and variant level pathogenicity prediction scores. Hence, our study subset, although somewhat limited, can be considered as a random sample of ASD-implicated genes and variants. Interestingly, we could not find any significant difference in pathogenicity prediction for disruptive variants *in vivo*. Of the 29 missense SSC-DNMs that had functional consequences in our assays, four were not predicted as damaging variants (CADD<10), nine had moderate scores (CADD: 10-20), and 16 were predicted to be disruptive (CADD>20). Understanding how variants that are not predicted to be damaging based on state-of-the-art informatics programs impact protein function may provide guidance to improve the accuracy of *in silico* tools.

To study the functional consequences of SSC-DNMs, we took two conceptually different approaches (rescue-based humanization strategy and overexpression-based strategy). Indeed, the two approaches were complementary as only two variants (*GLRA2*^*N136S*^ and *KCND3*^*R86P*^) were detected in both screens, showing consistent LoF effects using both approaches. Interestingly, *GRK4*, *NPFFR2*, and *PDK2* SSC-DNMs were found as LoF variants when overexpressed ubiquitously, yet these variants were able to rescue lethality in a similar manner to their respective reference alleles (Figure 4 and Table S5). This suggests that these variants are partial LoF alleles and different drivers and assays have different sensitivity. Moreover, two disruptive SSC-DNMs, *EPHB1*^*V916M*^ and *MAP4K1*^*M725T*^, behaved as complex alleles, displaying discordant phenotypes in the eye and wing. This suggests that these variants may behave in a context-dependent manner, acting as a GoF allele in one tissue while behaving as a LoF allele in another. Thus, one functional assay may not be enough to reveal the full nature of pathogenic mechanisms, and some disease-associated variants may act differently in different tissues or cell types.

Starting from a single *de novo* hemizygous missense variant that we identified as a LoF allele in *GLRA2* (p.N136S), we identified a cohort of subjects with overlapping neurodevelopmental phenotypes carrying LoF or GoF variants in this gene. Interestingly, female subjects harbored DNMs and male subjects carried maternally inherited variants in this X-linked gene, which undergoes random X-inactivation in females but not in males^47^. The X-linked status of *GLRA2* may mean that variants causing reduced *GLRA2* activity lead to disease in males but can be tolerated in heterozygous females. This is supported by non-symptomatic mothers of male probands who had maternally inherited alleles (Subjects 6-8). In contrast, GoF variants in this channel could be overrepresented in females since hyperactivation of this channel may cause neurological defects^48^. While the exact mechanism of how the p.T296M variant affects GLRA2 function remains unclear, the presence of melanized nodules in flies expressing this variant are indicative of an innate immune response^49^, potentially as a result of leaky ion channel function^50^. Fittingly, our structural analysis revealed that the p.T296 residue is adjacent to a critical amino acid that is likely important for keeping the ion pore in a conformationally closed state (Figure S9G).

In summary, we utilized a model organism-based *in vivo* functional genomics approach to study the functional consequence of rare genetic events in a common neurological disorder, ASD. In addition to garnering variant functional data for ASD subjects in the SSC, we leveraged this information to identify and document a novel rare neurological condition through matchmaking and collaboration. Such bi-directional communication and collaboration between bench scientists and clinicians greatly facilitate the functional studies of human variants found in common diseases such as ASD, and can also lead to novel discoveries that have impact on rare disease research.

## Supporting information

Extended Case Histories

## Acknowledgments

We express our appreciation to the patients and families for their participation in this study. We thank Pradnya Bhadane, Nora Duran, Mark Durham, Shelley Gibson, Yuchun He, Mei-Chu Huang, Matthew Lagarde, Wen-Wen Lin, and Dr. Karen Schulze for technical or administrative assistance. We thank Dr. Hugo Bellen for insightful scientific discussions valuable comments on this manuscript. We sincerely thank the late Dr. Kenneth Scott for access to many of the human cDNAs used in this study. This work was primarily supported by a Simons Foundation Autism Research Initiative (SFARI) Functional Screen Award (368479) to M.F.W. and S.Y. Generation of human cDNA transgenic lines were in part supported by NIH/ORIP (R24OD022005). Confocal microscopy is supported in part by NIH/NICHD (U54HD083092) to the Intellectual and Developmental Disabilities Research Center (IDDRC) Neurovisualization Core at BCM. P.C.M. is supported by CIHR (MFE-164712) and the Stand by Eli Foundation. J.A. is supported by NIH/NINDS (F32NS110174). R.M. is supported by NIH/NIGMS (T32GM07526-43), and through BCM Chao physician-scientist award. C.M.L. is part of the BCM Medical Scientist Training Program and McNair MD/PhD Student Scholars, supported by the McNair Medical Institute at the Robert and Janice McNair Foundation and NIH F30 Award (F30MH118804). H.T.C. received support by the American Academy of Neurology and CNCDP-K12. T.S.B. is supported by the Netherlands Organization for Scientific Research (ZonMW Veni, Grant 91617021), an Erasmus MC Fellowship 2017 and Erasmus MC Human Disease Model Award 2018. R.G. and A.V. received support from The DECODE-EE project (Health Research Call 2018, Tuscany Region). A.B. and L.P. received funding specifically appointed to Department of Medical Sciences from the Italian Ministry for Education, University and Research (Ministero dell’Istruzione, dell’Università e della Ricerca—MIUR) under the programme ‘Dipartimenti di Eccellenza 2018–2022’ Project code D15D18000410001.

## Author Contributions

M.F.W. and S.Y. conceived, designed the project. P.C.M., J.A., S.L.D., and J.M.H. designed and conducted most fly experiments and analyzed the data. V.H.B. and Y.C. performed cloning and mutagenesis. H.K.G. performed cloning and coordination. S.J. performed immunostaining and confocal microscopy. X.L. performed structural analysis and generated reagents. N.L. D.B., Y.C., P.L., B.H., H.P., C.M.L., H.T.C., H.C., N.A.H., O.K., and S.N.M contributed to reagent generation and some fly experiments. R.M., A.G., E.M.C.S., R.G., A.V., C.N.M., T.S.B., M.R.V., M.W., M.V., G.L., I.S., N.C., J.A.M., P.B.A. R.K., L.P., and A.B. reported and described *GLRA2* subjects. L.Z.R. and J.A.R. aided in subject match-making. P.C.M., J.A., S.L.D., J.M.H., R.M., M.F.W. and S.Y. wrote the manuscript.

## STAR Methods

### Additional gene/variant prioritization details

We prioritized genes and coding de novo missense mutations (DNMs) identified in the Simons Simplex Collection (SSC) from the initial study using exome sequencing^4^. In this cohort, 1,708 ASD proband-specific *de novo* missense or in-frame indels were identified through WES (Figure 1A), corresponding to 1,519 unique human genes. Of these, 920 fly genes corresponding to 1,032 human genes were identified. 487 human genes had no or weak ortholog candidates in *Drosophila* based on multiple ortholog prediction algorithms scores (cut off: DIOPT < 4/16)^51^ (DIOPT Version 8 accessed January 2020) (Supplementary Table 1). By overlapping these 920 *Drosophila* genes with available fly lines containing MiMIC transposons within coding introns that permit targeting of all annotated protein isoforms (1,732 insertions)^13,14^, we identified reagents for 122 fly genes corresponding to 143 human genes and 179 ASD proband variants from the SSC.

Of the 122 fly genes we selected to work on, we were able to successfully generate 109 TG4 lines through RMCE of MiMIC elements through genetic crosses (see below for specific methods)^15^. To generate UAS human reference transgenic (Ref-Tg) and human SSC candidate variant transgenic (SSC-Tg), we obtained human ORF (open reading frame) collections from the Mammalian Gene Collection^52^ or commercial sources. We generated SSC-Tg cDNAs by site-directed mutagenesis protocols (see below for specific methods). The reference and mutagenized ORF were shuttled into to the pUASg.HA-attB destination vector (Figure 1F)^53^. Transgenes were integrated into a precise location in the fly genome using ϕC31 transgenesis technology. Out of the 143 human genes and 179 SSC-DNMs of interest that we attempted to generate, we were successful in generating 194 UAS-cDNA (106 Ref-Tgs; 88 SSC-Tgs) flies (Figure 1G, Tables S2 and S3).

We were able to make a complete set of TG4, UAS-Ref-Tgs and UAS-SSC-Tgs lines for 65 fly genes corresponding to 74 human genes and 79 variants (some fly genes correspond to multiple human genes, and multiple SSC variants are found for a small subset of human genes), which were critical to test the variant function using a rescue-based humanization strategy. For an additional 42 human genes we generated 42 TG4 mutants. For 32 human genes we generated Ref-Tgs and for 9 genes we generated SSC-Tgs. In summary, 303 Drosophila stocks were generated for this project as a resource for the community, and these stocks are available from the Bloomington *Drosophila* Stock Center or in the process of being transferred and registered at BDSC.

### Generation of TG4 lines

All *TG4* alleles in this study were generated by ϕC31-mediated recombination-mediated cassette exchange of MiMIC (Minos mediated integration cassette) insertion lines^13,14,17^. Conversion of the original MiMIC element was performed via genetic by crossing *UAS-2xEGFP, hs-Cre,vas-dϕC31, Trojan T2A-GAL4* triplet flies to each MiMIC strain and following a crossing scheme^15^. 73 TG4 lines were described previously but not extensively characterized^16^, while 35 lines were generated specifically for this study.

### Generation of UAS-human cDNA lines

The majority of reference human cDNA clones were obtained in either pDONR221 or pDONR223 donor vectors. The LR clonase II (ThermoFisher) enzyme was used to shuttle ORFs into the p.UASg-HA.attB destination vector via Gateway™ cloning. Some ORFs that were not Gateway compatible were obtained from additional sources (Table S2), amplified with flanking *attB* sites and cloned into pDONR223 plasmid using BP clonase II (ThermoFisher). Sequence-verified variants were generated in the DONR vectors by either site-directed mutagenesis (SDM) via or High-Throughput Mutagenesis (HiTM) as previously described^54^. SDM was performed with primers generated using NEBaseChanger (Table S3) with the Q5® mutagenesis kit (NEB). Sequence-verified reference and variant ORFs in the pUASg-HA.attB destination plasmid were microinjected into ~200 embryos in one three *attP* docking sites (*attP86Fb*, *VK00037* or *VK00033*) docking sites by ϕC31 mediated transgenesis^53,55^. The docking site of choice were selected based on the genomic locus of the corresponding fly gene. In principal, VK00037 docking site on the 2^nd^ chromosome was used for human genes that correspond to fly genes on the X, 3^rd^ or 4^th^ chromosome, whereas VK00033 or attP86Fb docking site on the 3^rd^ chromosome was used for human genes that correspond to fly genes on the 2^nd^ chromosome.

### Fly husbandry

Unless otherwise noted, all flies used in experiments were grown in a temperature and humidity-controlled incubator at 25°C and 50% humidity on a 12-hour light/dark cycle.

Some experiments were conducted at different temperatures that are specifically indicated in the text and figures. Stocks were reared on standard fly food (water, yeast, soy flour, cornmeal, agar, corn syrup, and propionic acid) at room temperature (~22°C) and routinely maintained.

### Fly stocks used that were not generated here

*tub-GAL4* (*y*^*1*^ *w*^*^; *P{w[+mC]=tubP-GAL4}LL7/TM3, Sb*^*1*^ *Ser*^*1*^) BDSC_5138, *GMR-GAL4* (*w^*^; P{w[+mC]=GAL4-ninaE.GMR}12*) BDSC_1104, *nub-GAL4* (*P{GawB}nubbin-AC-62*)^56^, *nSyb-GAL4* (*y*^*1*^ *w*^*;^; *P{nSyb-GAL4.S}3*) BDSC_51635, *Rh1-GAL4* (*P{ry[+t7.2]=rh1-GAL4}3, ry[506])*BDSC_8691, *pnr-GAL4* (*y*^*1*^ *w*^*1118*^; *P{w[+mW.hs]=GawB}pnr[MD237]/TM3, P{w[+mC]=UAS-y.C}MC2, Ser*^*1*^) BDSC_3039, *UAS-LacZ* (*w^*^; P{w[+mC]=UAS-lacZ.Exel}2*) BDSC _8529, UAS-nlsGFP (*w*^*1118*^; *P{w[+mC]=UAS-GFP.nls}14*) BDSC_4775.

### Electroretinograms (ERG)

ERG recordings on adult flies were performed on *nSyb-GAL4*^42^ and *Rh1-GAL4*^44^ driven *UAS-GLRA2* at 5 days post-eclosion raised at 25°C in 12h light/12h dark cycle as previously described^57^ using LabChart software (AD instruments). 4-10 flies were examined for each genotype. Recording was repeated at least 3 times per fly. Quantification and statistical analysis was performed using ANOVA followed by Bonferroni’s multiple comparison test using Prism 8.0.

### Complementation test of lethality in TG4 lines

Out of the 109 TG4 mutants generated, 64 TG4 mutants were homozygous lethal. Because lethality can be caused by disruption of the gene of interest or due to second site lethal mutations carried on the same chromosome, we performed complementation test using standard methodology. For genes on the 2^nd^ and 3^rd^ chromosome, female heterozygous TG4 lines balanced with either *SM6a* or *TM3, Sb, Ser*, respectively, were crossed with male flies carrying a corresponding deficiency (Df) that covers the gene of interest (see Supplementary table 4). Three independent crosses were set at 25°C for each TG4 line and we determined if any *TG4* flies survived to the adult stage *in trans* with their corresponding Df (*TG4/Df*). If viable, a second Df line covering the same gene was used to validate this finding to make sure the complementation is not due to some problematic Df lines. If TG4 was viable over two independent Df lines, we ascribed the lethality to a second site mutation on the *TG4* chromosome. *TG4* that remained lethal in trans with a Df line are be considered to be disrupting an essential gene in flies. For five genes on the X-chromosome of the fly, complementation was performed by first rescuing hemizygous TG4 males with a duplication (Dp) line obtained from BDSC (Table S4), and crossing these rescued flies to female TG4/FM7 flies. If TG4/Y; Dp/+ lines were viable, we ascribed the lethality of TG4 to the gene of interest. All Df and Dp lines were obtained from BDSC, and the specific stock used in our analysis are listed in Supplementary Table 4.

Through this experiment, we found 64 *TG4* mutant lines that were homozygous lethal, and 47 remained lethal when *in trans* with a corresponding deficiency line (Figure S1A, Table S4). The 47 essential genes in *D. mel* corresponded to 60 SSC related human genes (Figure S1B). The lethality of 17 *TG4* lines corresponding to 18 human genes were due to a second site lethal mutation, potentially present in the original MiMIC line, or introduced during RMCE which has been reported previously^14^. These TG4 lines together with viable TG4 lines are likely associated with non-essential genes in *Drosophila*.

### Rescue of lethality in TG4 lines by UAS-human cDNA transgenes

In order to assess the ability of human reference or SSC variant cDNAs to rescue lethality observed in *TG4* mutants in essential genes, we first double balanced all Df lines that fail to complement a lethal TG4 line with UAS-reference or variant cDNA lines. For genes on the 2^nd^ chromosome, we generated *Df/CyO; UAS-cDNA/(TM3, Sb, Ser)* stocks. For genes on the 3^rd^ chromosome, we generated *UAS-cDNA/(CyO); Df/TM3, Sb, Ser.* Heterozygous *TG4/Balancer* females were crossed to double balanced Df/Balancer, UAS-human cDNA males at multiple temperatures (18°C, 22°C, 25°C, 29°C) to determine rescue of lethality to adult stage. A minimum of two independent crosses were conducted at each temperature. For the five genes on the X-chromosome of the fly, we attempted rescue by crossing female *TG4/FM7* flies to UAS-cDNA/(SM6a) males to generate hemizygous TG4 males that expresses human cDNA (TG4/Y; UAS-cDNA/+) to test their viability. Statistical analysis was performed using Two-way ANOVA followed by Sidak’s multiple comparison test across temperature and genotype.

### Lifespan assays

Lifespan analysis was performed as previously described^58^. Briefly, newly eclosed flies were separated by genotype and sex and incubated at 25°C. Flies were transferred into a fresh vial every two days and survival was determined once a day. 11-49 flies were tested per group. Statistical analysis was performed using Log-rank (Mantel-Cox) test.

### Behavioral assays

Of 48 *TG4* mutants that were viable when *in trans* with a corresponding deficiency, 17 lines exhibited lethality in homozygous states, indicating the presence of a second site lethal mutation. Out of 31 *TG4* mutants that were homozygous viable, we prioritized to study 21 *TG4* mutants based on reagent availability. Courtship assay was performed as previously described^22^. *TG4* lines were backcrossed to *Canton-S* strain in order to eliminate known courtship deficiencies present in the *y*^*1*^ *w*^*^ background, which all TG4 lines are initially generated on. Collection of socially naïve adults was performed by isolating pupae in 16 x 100 polystyrene vials containing approximately 1 ml of fly food. After eclosion, flies were anesthetized briefly with CO2 to ensure they were healthy and lacking wing damage. Anesthetized flies were returned to their vials and allowed 24 hours to recover before testing. Courtship assays were performed in a 6 well acrylic plate with 40mm circular wells, with a depth of 3mm and a slope of 11 degrees, as per the chamber design in Simon et al, 2010^59^. One Canton-S virgin female (6-10 days post-eclosion), and one *TG4* mutant male fly (3-5 days post-eclosion) with or without UAS-human cDNAs were simultaneously introduced into the chamber via aspiration. Recordings were taken using a Basler 1920UM, 1.9MP, 165FPS, USB3 Monochromatic camera using the BASLER Pylon module, with an adjusted capturer rate of 33 fps (frames per second). Conversion of captured images into a movie file was performed via a custom MatLab script, and tracking of flies in the movie was performed using the Caltech Flytracker^60^. Machine learning assessment of courtship was performed using JAABA^61^ using classifiers that scored at 95% or higher accuracy during ground-truthing trials. At least 10 animals were tested per genotype. Analysis of data was performed using Excel (Microsoft) and Prism (GraphPad). A ROUT (Q=1%) test was performed in Prism to identify outliers. Determination of significance in behavior tests was performed using the Kruskal-Wallis one-way analysis of variance and the Dunn’s multiple comparison test. P-values of 0.05 or less were considered significant.

### Overexpression assays to assess lethality and morphological phenotypes

To detect any differences in the phenotypes induced by overexpression of reference and variant human cDNA in order to assess variant function, we crossed UAS-human cDNAs with reference or variant alleles to ubiquitous (*tub-GAL4*)^62^, wing (*nub-GAL4*)^56^ or eye (*GMR-GAL4*)^63^ specific drivers. In the ubiquitous expression screen, 3-4 virgin females of *tub-GAL4/TM3 Sb* flies were crossed to 2-4 males of the UAS-cDNA reference and variant at 25°C. After 3-4 days, the parents were transferred into new vials, and the new vial was placed at 29°C while the old vial was kept at 25°C, allowing us to test two temperatures simultaneously. The parents were discarded after 3-5 days. Flies were collected after most of the pupae eclosed. The total number of flies were counted and scored with the genotype of interest (i.e. *tub-GAL4>UAS-cDNA*) as well as all other genotypes, (i.e. genotypes with balancers). A minimum of 10 flies were scored per experiment, though for the majority of crosses 50-100 flies were scored in this primary analysis. Viability was calculated by taking the % of observed/expected based on Mendelian ratio, and any UAS-cDNA with survival less than 70% was recorded as having scorable phenotype (lethal or semi-lethal). All of lines showing a phenotype at 29°C also showed phenotypes at 25°C, so subsequent experiments were performed at 25°C. To validate our hits, we performed the same viability assay, except each UAS-cDNA was tested at least three times to statistically validate that there is a difference between reference and variant. In addition, two independent UAS-cDNA transgenic lines established from the same construct were tested for each reference and variant. A variant was considered to have functional consequence (true hit) if both transgenic lines showed the same phenotype. In the cases where the difference is rather minor (e.g. <20% difference between survival), this was considered within the variation of the experiment paradigm, and the variant phenotype was documented. Functional study using wing or eye drivers were performed using similar strategies, but morphological phenotypes were scored instead of lethality.

### Imaging of adult fly morphology

*Drosophila* eyes, wings and nota (dorsal thorax) were imaged after flies were frozen at - 20°C for at least 24 hours. Wings for some flies were dissected in 70% EtOH and mounted onto slides for imaging. Images were obtained with the Leica MZ16 stereomicroscope equipped with Optronics MicroFire Camera and Image Pro Plus 7.0 software to extend the depth-of-field for Z-stack images.

### Expression analysis of TG4 lines in larval and adult brains

All *TG4* lines are crossed with UAS nlsGFP (3^rd^ chromosome) at room temperature. The brains of GFP positive third instar larvae and 3-5 days old adult flies were dissected in 1X phosphate-buffered saline (PBS). Adult brains were fixed immediately in 4% paraformaldehyde (PFA) and incubated at 4°C overnight (o/n) on a shaker. Next day these brains were post-fixed with 4% PFA with 2% Triton-X in PBS (PBST), kept in a vacuum container for an hour to get rid of the air from the tracheal tissue also make the tissue more permissive. Fixative was replaced every 10 minutes during this post-fixation step. Larval brains were fixed for 50 minutes on a rotator at room temperature. After thorough washing with PBS with 0.2% Triton (PBTX) both adult and larval brains were incubated with primary antibodies overnight (o/n) at 4°C on shaker. The sample were extensively washed with 0.2% PBTX before secondary antibodies were applied at room temperature for 2 hours. Samples were thoroughly washed with PBST and mounted on a glass slide using Vectasheild (Vector Labs, H-1000-10). Primary antibodies used: Mouse anti-repo (DSHB: 8D12) 1:50, Rat anti-elav (DSHB: 7E8A10) 1:100, Goat anti-GFP (ABCAM: ab6662) 1:500. Secondary antibodies used: Anti-mouse-647 (Jackson ImmunoResearch: 715-605-151) 1:250, Anti-rat-Cy3 (Jackson ImmunoResearch: 712-165-153) 1:500. The samples were scanned using a laser confocal microscope (Zeiss LSM 880), and images were processed using ZEN (Zeiss) and Imaris (Oxford Instruments) software.

### GeneOntology (GO) analysis

GO analysis was performed based on the PANTHER (Protein Analysis Through Evolutionary Relationships) system (http://www.pantherdb.org; date last accessed October 31, 2020^30^. Statistical analysis was performed by using the default PANTHER Overrepresentation Test (Released 20200728), Annotation Version and Release Date: GO Ontology database DOI: 10.5281/zenodo.4033054 Released 2020-09-10 which used the Fisher’s Exact test with a false discovery rate p<0.05.

### Patient recruitment and consent

Affected individuals were investigated by their referring physicians at local sites. Prior to research studies, informed consent was obtained according to the institutional review boards (IRB) and ethnics committees of each institution. Individuals who were ascertained in diagnostic testing procedures (and/or their legal guardians) gave clinical written informed consent for testing, and their permission for inclusion of their anonymized data in this cohort series. This was obtained using standard forms at each local site by the responsible referring physicians.

### Exome sequencing and identification of GLRA2 variants

Subjects 1, 2 and 4 had clinical exome sequencing at GeneDx (Gaithersburg, MD, United States), at the Praxis für Humangenetik Tubingen (Tubingen, Germany), and at Baylor Genetics (Houston, TX, United States), respectively. Subject 3 WES was performed at the Meyer Children’s Hospital, University of Florence, in the context of the DESIRE program and as previously described^64^. Briefly, the SureSelectXT Clinical Research Exome kit (Agilent Technologies, Santa Clara, CA) was used for library preparation and target enrichment, and paired-end sequencing was performed using Illumina sequencer (NextSeq550, Illumina, San Diego, CA, USA) to obtain an average coverage of above 80x, with 97.6% of target bases covered at least 10x. Reads were aligned to the GRCh37/hg19 human genome reference assembly by the BWA software package, and the GATK suite was used for base quality score recalibration, realignment of insertion/deletions (InDels), and variant calling^65,66^. Variant annotation and filtering pipeline included available software (VarSeq, Golden Helix, Inc v1.4.6), focusing on non-synonymous/splice site variants with minor allele frequency (MAF) lower than 0.01 in the GnomAD database^26^ (http://gnomad.broadinstitute.org/), an internal healthy control database and pre-computed genomic variants score from dbNSFP^67^. Subject 5 had exome sequencing at Lyon Universiy Hospital (Lyon, France). The SeqCap EZ Medexome kit (Roche, Pleasanton, CA, USA) was used for library preparation and target enrichment before paired-end sequencing using an Illumina instrument (NextSeq500, Illulina, San Diego, CA, USA). A mean depth of coverage of 133x was obtained with 99.0% of target bases covered at least 10x. Reads were aligned to the GRCh37/hg19 human genome reference assembly by the BWA software package, and the GATK suite was used for base quality score recalibration, realignment of insertion/deletions (InDels), and variant calling^65,66^. Variant annotation was performed with SnpEFF and filtering pipeline focused on non-synonymous/splice site variants with minor allele frequency (MAF) lower than 0.01 in the GnomAD database^26^ (http://gnomad.broadinstitute.org/).

Subject 6 WES was performed at the Erasmus MC as previously described^68^. In brief, exome-coding DNA was captured with the Agilent SureSelect Clinical Research Exome (CRE) kit (v2). Sequencing was performed on an Illumina HiSeq 4000 platform with 150-bp paired-end reads. Reads were aligned to hg19 using BWA (BWA-MEM v0.7.13) and variants were called using the GATK Haplotype Caller^65^ v3.7 (https://www.broadinstitute.org/gatk/). Detected variants were annotated, filtered and prioritized using the Bench lab NGS v5.0.2 platform (Agilent technologies). Subject 7 WES and data processing were performed by the Genomics Platorm at the Broad Institute of MIT and Harvard with an Illumina Nextera or Twist exome capture (~38 Mb target), and sequenced (150 bp paired reads) to cover >80% of targets at 20x and a mean target coverage of >100x. WES data was processed through a pipeline based on Picard and mapping done using the BWA aligner to the human genome build 38. Variants were called using Genome Analysis Toolkit (GATK) HaplotypeCaller package version 3.5^65^ (https://www.broadinstitute.org/gatk/). Subject 8 WES was performed in collaboration with the Autism Sequencing Consortium (ASC) at the Broad Institute on Illumina HiSeq sequencers using the Illumina Nextera exome capture kit. Exome sequencing data was processed through a pipeline based on Picard and mapping done using the BWA aligner to the human genome build 37 (hg19). Variants were called using Genome Analysis Toolkit (GATK) HaplotypeCaller package version 3.4^65^(https://www.broadinstitute.org/gatk/). Variant call accuracy was estimated using the GATK Variant Quality Score Recalibration (VQSR) approach. High-quality variants with an effect on the coding sequence or affecting splice site regions were filtered against public databases (dbSNP150 and gnomAD V.2.0) to retain (i) private and clinically associated variants; and (ii) annotated variants with an unknown frequency or having minor allele frequency <0.1%, and occurring with a frequency <2% in an in-house database including frequency data from > 1,500 population-matched WES. The functional impact of variants was analyzed by CADD V.1.3, Mendelian Clinically Applicable Pathogenicity V.1.0^29,69^, and using InterVar V.0.1.6 to obtain clinical interpretation according to American College of Medical Genetics and Genomics/Association for Molecular Pathology 2015 guidelines^70^. GeneMatcher^33^ (https://genematcher.org/) assisted in the recruitment of Subjects 2, 3 and 5-8.

### Western blot

Five heads of *nSyb-GAL4 UAS-GLRA2* reference and variant flies aged for 5 days post eclosion were lysed in 30µL NETN buffer (50mM Tris pH 7.5, 150mM NaCl, 0.5% NP-40, 1 mM EDTA) with an electric douncer for 10 seconds for three times on ice. 30µL of 2x Laemmli Sample Buffer (Bio-Rad) with 10% 2-mercaptoethanol was added to the lysis and incubated on ice for 10 min. Samples were boiled at 95°C and spun at 14,000 RPM for 5 minutes at 4 °C. The soluble fraction was loaded onto a standard SDS-PAGE gel. PVDF (polyvinylidene difluoride) membrane activated for 1 minute with 100% methanol. After running and wet transfer, the membrane was blocked in 5% skim milk for 1 hour. The membrane was incubated (overnight, shaking, at 4°C) with mouse anti-HA (HA.11, 1:1,000, 901501, BioLegend) and mouse anti-Actin (C4) (1:50,000, MAB1501, EMD Millipore) primary antibodies in 3% BSA (bovine serum albumin), followed by 10 minute washes (3 times) with 1% Triton-X in Tris-buffered saline (TBST). We incubated this with goat anti-mouse HRP-conjugated (1:15000, 115-035-146, Jackson ImmunoResearch) secondary antibody in skim milk. The membrane was washed three times with 1% TBST and detected with Western Lightning™ Chemiluminescence Reagent Plus (perkinelmerNEL104001EA) ECL solution using the Bio-Rad ChemiDoc MP imaging system.

### Structural biological analysis of GLRA2 patient variants

Protein residues that corresponds to GLRA2 patient variants were mapped onto the crystal protein structure of GLRA1 protein in Protein Data Bank (PBD, ID: 4X5T)^71^ using the PyMOL (https://pymol.org/)^72^ because GLRA1 and GLRA2 are highly homologous proteins (85% similarity, 78% identity and 3% gaps) based on DIOPT^51^.

### Image generation

Cartoon images in Figure 1H, S1A and 2G were generated with BioRender.com.

