## Supplementary material for "*Drosophila* functional screening of *de novo* variants in autism uncovers deleterious variants and facilitates discovery of rare neurodevelopmental diseases": Table S1

**Supplemental Table 1: Conserved genes from Simon's Simplex Collection**

| Human Gene | HGNC | FlyBaseID | Fly Gene |
|---|---|---|---|
| *A2M* | 7 | FBgn0041181 | *Tep3* |
| *A2ML1* | 23336 | FBgn0041180 | *Tep4* |
| *AASS* | 17366 | FBgn0286198 | *LKRSDH* |
| *ABCA1* | 29 | FBgn0083956 | *CG34120* |
| *ABCA13* | 14638 | FBgn0083956 | *CG34120* |
| *ABCA2* | 32 | FBgn0031171 | *CG1801* |
| *ABCA7* | 37 | FBgn0083956 | *CG34120* |
| *ABCA8* | 38 | FBgn0034493 | *CG8908* |
| *ABCB6* | 47 | FBgn0038376 | *Hmt-1* |
| *ABCC4* | 55 | FBgn0038740 | *CG4562* |
| *ABCC5* | 56 | FBgn0039644 | *rdog* |
| *ABCG1* | 73 | FBgn0020762 | *Atet* |
| *ABCG2* | 74 | FBgn0003996 | *w* |
| *ABHD12* | 15868 | FBgn0034419 | *CG15111* |
| *ABI2* | 24011 | FBgn0020510 | *Abi* |
| *ABL1* | 76 | FBgn0000017 | *Abl* |
| *ABL2* | 77 | FBgn0000017 | *Abl* |
| *ABR* | 81 | FBgn0025836 | *RhoGAP1A* |
| *ACACB* | 85 | FBgn0033246 | *ACC* |
| *ACE* | 2707 | FBgn0012037 | *Ance* |
| *ACHE* | 108 | FBgn0000024 | *Ace* |
| *ACP2* | 123 | FBgn0000032 | *Acph-1* |
| *ACTN4* | 166 | FBgn0000667 | *Actn* |
| *ACTR6* | 24025 | FBgn0011741 | *Arp6* |
| *ACTRT3* | 24022 | FBgn0000045 | *Act79B* |
| *ADAM18* | 196 | FBgn0259110 | *mmd* |
| *ADAMTS7* | 223 | FBgn0029791 | *CG4096* |
| *ADAMTSL1* | 14632 | FBgn0051619 | *nolo* |
| *ADAMTSL4* | 19706 | FBgn0032252 | *loh* |
| *ADCY5* | 236 | FBgn0263131 | *CG43373* |
| *ADD3* | 245 | FBgn0263391 | *hts* |
| *ADRBK2* | 290 | FBgn0260798 | *Gprk1* |
| *AEBP2* | 24051 | FBgn0086655 | *jing* |
| *AGAP1* | 16922 | FBgn0028509 | *CenG1A* |
| *AGAP2* | 16921 | FBgn0028509 | *CenG1A* |
| *AGK* | 21869 | FBgn0260750 | *Mulk* |
| *AGO1* | 3262 | FBgn0262739 | *AGO1* |
| *AGTRAP* | 13539 | FBgn0052638 | *CG32638* |
| *AK1* | 361 | FBgn0022709 | *Adk1* |
| *AKAP1* | 367 | FBgn0263987 | *spoon* |
| *AKAP9* | 379 | FBgn0086690 | *Plp* |
| *AKR1B15* | 37281 | FBgn0086254 | *CG6084* |
| *AKR1C2* | 385 | FBgn0086254 | *CG6084* |

| | | | |
|---|---|---|---|
| *AKR1D1* | 388 | FBgn0086254 | *CG6084* |
| *AKT2* | 392 | FBgn0010379 | *Akt1* |
| *ALDH18A1* | 9722 | FBgn0037146 | *CG7470* |
| *ALDH1L1* | 3978 | FBgn0032945 | *CG8665* |
| *ALDH3A1* | 405 | FBgn0010548 | *Aldh-III* |
| *ALDH5A1* | 408 | FBgn0039349 | *Ssadh* |
| *ALS2* | 443 | FBgn0037116 | *Als2* |
| *AMPD2* | 469 | FBgn0052626 | *AMPdeam* |
| *AMY2B* | 478 | FBgn0000079 | *Amy-p* |
| *ANGPT2* | 485 | FBgn0087011 | *CG41520* |
| *ANK2* | 493 | FBgn0011747 | *Ank* |
| *ANK3* | 494 | FBgn0011747 | *Ank* |
| *ANKRD17* | 23575 | FBgn0043884 | *mask* |
| *ANO3* | 14004 | FBgn0036235 | *CG6938* |
| *ANO6* | 25240 | FBgn0036235 | *CG6938* |
| *ANP32D* | 16676 | FBgn0034282 | *Mapmodulin* |
| *AP1S2* | 560 | FBgn0039132 | *AP-1sigma* |
| *AP2S1* | 565 | FBgn0043012 | *AP-2sigma* |
| *AP3B2* | 567 | FBgn0003210 | *rb* |
| *AP3D1* | 568 | FBgn0001087 | *g* |
| *APAF1* | 576 | FBgn0263864 | *Dark* |
| *APBA1* | 578 | FBgn0052677 | *X11Lbeta* |
| *APC2* | 24036 | FBgn0015589 | *Apc* |
| *APOB* | 603 | FBgn0087002 | *apolpp* |
| *AQPEP* | 26904 | FBgn0051445 | *CG31445* |
| *AREL1* | 20363 | FBgn0031384 | *CG4238* |
| *ARHGAP21* | 23725 | FBgn0031118 | *RhoGAP19D* |
| *ARHGAP30* | 27414 | FBgn0032821 | *CdGAPr* |
| *ARHGEF10L* | 25540 | FBgn0263706 | *CG43658* |
| *ARHGEF11* | 14580 | FBgn0023172 | *RhoGEF2* |
| *ARHGEF16* | 15515 | FBgn0261547 | *Exn* |
| *ARHGEF4* | 684 | FBgn0264707 | *RhoGEF3* |
| *ARID1B* | 18040 | FBgn0261885 | *osa* |
| *ARID2* | 18037 | FBgn0042085 | *Bap170* |
| *ARMC3* | 30964 | FBgn0033794 | *CG13326* |
| *ARPP21* | 16968 | FBgn0004875 | *enc* |
| *ARX* | 18060 | FBgn0000061 | *al* |
| *ASL* | 746 | FBgn0032076 | *Argl* |
| *ASPM* | 19048 | FBgn0000140 | *asp* |
| *ATAD5* | 25752 | FBgn0036574 | *elg1* |
| *ATG2B* | 20187 | FBgn0044452 | *Atg2* |
| *ATIC* | 794 | FBgn0039241 | *CG11089* |
| *ATP10A* | 13542 | FBgn0032120 | *CG33298* |
| *ATP11C* | 13554 | FBgn0030746 | *CG9981* |
| *ATP12A* | 13816 | FBgn0002921 | *Atpalpha* |

| | | | |
|---|---|---|---|
| *ATP1A4* | 14073 | FBgn0002921 | *Atpalpha* |
| *ATP2A1* | 811 | FBgn0263006 | *SERCA* |
| *ATP2B2* | 815 | FBgn0259214 | *PMCA* |
| *ATP2B4* | 817 | FBgn0259214 | *PMCA* |
| *ATP4A* | 819 | FBgn0002921 | *Atpalpha* |
| *ATP6V1A* | 851 | FBgn0265262 | *Vha68-1* |
| *ATP8B2* | 13534 | FBgn0037989 | *ATP8B* |
| *B9D1* | 24123 | FBgn0038342 | *B9d1* |
| *BAIAP2L1* | 21649 | FBgn0052082 | *IRSp53* |
| *BAZ1A* | 960 | FBgn0027620 | *Acf* |
| *BAZ2B* | 963 | FBgn0033636 | *tou* |
| *BCCIP* | 978 | FBgn0038183 | *CG9286* |
| *BCHE* | 983 | FBgn0000024 | *Ace* |
| *BCORL1* | 25657 | FBgn0036814 | *CG14073* |
| *BEST3* | 17105 | FBgn0040238 | *Best1* |
| *BIRC6* | 13516 | FBgn0266717 | *Bruce* |
| *BLK* | 1057 | FBgn0262733 | *Src64B* |
| *BMP1* | 1067 | FBgn0004885 | *tok* |
| *BNC1* | 1081 | FBgn0000459 | *disco* |
| *BRD1* | 1102 | FBgn0033155 | *Br140* |
| *BRD3* | 1104 | FBgn0004656 | *fs(1)h* |
| *BRD4* | 13575 | FBgn0004656 | *fs(1)h* |
| *BRF1* | 11551 | FBgn0038499 | *Brf* |
| *BRPF1* | 14255 | FBgn0033155 | *Br140* |
| *BRPF3* | 14256 | FBgn0033155 | *Br140* |
| *BRSK2* | 11405 | FBgn0036544 | *sff* |
| *BTBD2* | 15504 | FBgn0262871 | *lute* |
| *C10orf137* | 24640 | FBgn0035923 | *CG6511* |
| *C19orf70* | 33702 | FBgn0036726 | *QIL1* |
| *C2CD3* | 24564 | FBgn0052425 | *CG32425* |
| *C2orf42* | 26056 | FBgn0039663 | *CG2321* |
| *C4orf27* | 26051 | FBgn0037377 | *CG1218* |
| *C9orf156* | 30967 | FBgn0033229 | *CG12822* |
| *CA3* | 1374 | FBgn0027844 | *CAH1* |
| *CACNA1C* | 1390 | FBgn0001991 | *Ca-alpha1D* |
| *CACNA1D* | 1391 | FBgn0001991 | *Ca-alpha1D* |
| *CACNA1E* | 1392 | FBgn0263111 | *cac* |
| *CACNA1G* | 1394 | FBgn0264386 | *Ca-alpha1T* |
| *CACNA1H* | 1395 | FBgn0264386 | *Ca-alpha1T* |
| *CACNA1S* | 1397 | FBgn0001991 | *Ca-alpha1D* |
| *CACNA2D1* | 1399 | FBgn0261041 | *stj* |
| *CAD* | 1424 | FBgn0003189 | *r* |
| *CAMK2A* | 1460 | FBgn0264607 | *CaMKII* |
| *CAMSAP1* | 19946 | FBgn0263197 | *Patronin* |
| *CAP2* | 20039 | FBgn0261458 | *capt* |

| | | | |
|---|---|---|---|
| *CAPN10* | 1477 | FBgn0260450 | *CalpC* |
| *CAPN12* | 13249 | FBgn0025866 | *CalpB* |
| *CAPRIN2* | 21259 | FBgn0042134 | *Capr* |
| *CARS* | 1493 | FBgn0027091 | *CysRS* |
| *CARS2* | 25695 | FBgn0033900 | *CysRS-m* |
| *CASK* | 1497 | FBgn0013759 | *CASK* |
| *CAT* | 1516 | FBgn0000261 | *Cat* |
| *CBL* | 1541 | FBgn0020224 | *Cbl* |
| *CCBL2* | 33238 | FBgn0037955 | *Kyat* |
| *CCDC22* | 28909 | FBgn0036671 | *CG9951* |
| *CCDC28B* | 28163 | FBgn0031395 | *CG10874* |
| *CCDC65* | 29937 | FBgn0050259 | *CG30259* |
| *CCDC88C* | 19967 | FBgn0283724 | *Girdin* |
| *CCNB1* | 1579 | FBgn0000405 | *CycB* |
| *CCNB3* | 18709 | FBgn0015625 | *CycB3* |
| *CCNJL* | 25876 | FBgn0010317 | *CycJ* |
| *CCT4* | 1617 | FBgn0032444 | *CCT4* |
| *CCT6B* | 1621 | FBgn0027329 | *CCT6* |
| *CD151* | 1630 | FBgn0036769 | *Tsp74F* |
| *CDC34* | 1734 | FBgn0036516 | *CG7656* |
| *CDC42BPB* | 1738 | FBgn0023081 | *gek* |
| *CDK13* | 1733 | FBgn0037093 | *Cdk12* |
| *CDK18* | 8751 | FBgn0005640 | *Eip63E* |
| *CDK19* | 19338 | FBgn0015618 | *Cdk8* |
| *CEBPG* | 1837 | FBgn0036126 | *Irbp18* |
| *CEP135* | 29086 | FBgn0036480 | *Cep135* |
| *CERS6* | 23826 | FBgn0040918 | *schlank* |
| *CES2* | 1864 | FBgn0027584 | *CG4757* |
| *CHD3* | 1918 | FBgn0262519 | *Mi-2* |
| *CHD4* | 1919 | FBgn0262519 | *Mi-2* |
| *CHD7* | 20626 | FBgn0266557 | *kis* |
| *CHD8* | 20153 | FBgn0266557 | *kis* |
| *CHD9* | 25701 | FBgn0266557 | *kis* |
| *CHKB* | 1938 | FBgn0032955 | *CG2201* |
| *CHRNB1* | 1961 | FBgn0000038 | *nAChRbeta1* |
| *CHST2* | 1970 | FBgn0051637 | *CG31637* |
| *CHSY1* | 17198 | FBgn0030662 | *CG9220* |
| *CIC* | 14214 | FBgn0262582 | *cic* |
| *CISH* | 1984 | FBgn0033266 | *Socs44A* |
| *CIT* | 1985 | FBgn0002466 | *sti* |
| *CLASP1* | 17088 | FBgn0021760 | *chb* |
| *CLCN7* | 2025 | FBgn0033755 | *ClC-b* |
| *CLCNKB* | 2027 | FBgn0051116 | *ClC-a* |
| *CLIP2* | 2586 | FBgn0020503 | *CLIP-190* |
| *CMAS* | 18290 | FBgn0052220 | *Csas* |

| | | | |
|---|---|---|---|
| *CNGB1* | 2151 | FBgn0266346 | *CngB* |
| *CNOT1* | 7877 | FBgn0085436 | *Not1* |
| *CNOT4* | 7880 | FBgn0051716 | *Cnot4* |
| *CNOT6* | 14099 | FBgn0011725 | *twin* |
| *CNPY2* | 13529 | FBgn0263260 | *sel* |
| *CNTN6* | 2176 | FBgn0037240 | *Cont* |
| *CNTNAP4* | 18747 | FBgn0013997 | *Nrx-IV* |
| *COL4A3BP* | 2205 | FBgn0027569 | *cert* |
| *CPA4* | 15740 | FBgn0029804 | *CG3097* |
| *CPT1B* | 2329 | FBgn0261862 | *whd* |
| *CPT1C* | 18540 | FBgn0261862 | *whd* |
| *CR1* | 2334 | FBgn0032797 | *Hasp* |
| *CREBBP* | 2348 | FBgn0261617 | *nej* |
| *CREBL2* | 2350 | FBgn0032202 | *REPTOR-BP* |
| *CRHR1* | 2357 | FBgn0033744 | *Dh44-R2* |
| *CROCC* | 21299 | FBgn0039152 | *Root* |
| *CRTC3* | 26148 | FBgn0036746 | *Crtc* |
| *CRY2* | 2385 | FBgn0016054 | *phr6-4* |
| *CSAD* | 18966 | FBgn0000153 | *b* |
| *CSNK2A1* | 2457 | FBgn0264492 | *CkIIalpha* |
| *CTCF* | 13723 | FBgn0035769 | *CTCF* |
| *CTNNB1* | 2514 | FBgn0000117 | *arm* |
| *CUL5* | 2556 | FBgn0039632 | *Cul5* |
| *CYP3A43* | 17450 | FBgn0038037 | *Cyp9f2* |
| *CYP4F12* | 18857 | FBgn0015032 | *Cyp4c3* |
| *CYP4F3* | 2646 | FBgn0033395 | *Cyp4p2* |
| *CYP4Z1* | 20583 | FBgn0005670 | *Cyp4d1* |
| *DAP3* | 2673 | FBgn0034727 | *mRpS29* |
| *DARS2* | 25538 | FBgn0051739 | *AspRS-m* |
| *DBR1* | 15594 | FBgn0035838 | *ldbr* |
| *DCAF11* | 20258 | FBgn0034527 | *CG9945* |
| *DCAF12L1* | 29395 | FBgn0037980 | *DCAF12* |
| *DCAF5* | 20224 | FBgn0250755 | *CG42233* |
| *DCLK1* | 2700 | FBgn0261387 | *CG17528* |
| *DDR2* | 2731 | FBgn0053531 | *Ddr* |
| *DDX20* | 2743 | FBgn0011802 | *Gem3* |
| *DDX23* | 17347 | FBgn0032690 | *CG10333* |
| *DENND4A* | 24321 | FBgn0025864 | *Crag* |
| *DENND5B* | 28338 | FBgn0035229 | *pns* |
| *DGAT1* | 2843 | FBgn0004797 | *mdy* |
| *DGCR14* | 16817 | FBgn0023506 | *Es2* |
| *DHX9* | 2750 | FBgn0002774 | *mle* |
| *DICER1* | 17098 | FBgn0039016 | *Dcr-1* |
| *DIDO1* | 2680 | FBgn0082831 | *pps* |
| *DIS3L2* | 28648 | FBgn0035111 | *Dis3l2* |

| | | | |
|---|---|---|---|
| *DLC1* | 2897 | FBgn0285955 | *cv-c* |
| *DLGAP2* | 2906 | FBgn0259978 | *vlc* |
| *DLX3* | 2916 | FBgn0000157 | *Dll* |
| *DMPK* | 2933 | FBgn0023081 | *gek* |
| *DMXL2* | 2938 | FBgn0023458 | *Rbcn-3A* |
| *DNAAF1* | 30539 | FBgn0023090 | *dtr* |
| *DNAH10* | 2941 | FBgn0013813 | *Dhc98D* |
| *DNAH11* | 2942 | FBgn0013812 | *Dhc93AB* |
| *DNAH17* | 2946 | FBgn0013812 | *Dhc93AB* |
| *DNAH2* | 2948 | FBgn0001313 | *kl-2* |
| *DNAH5* | 2950 | FBgn0037726 | *CG9492* |
| *DNAH7* | 18661 | FBgn0013810 | *Dhc36C* |
| *DNAH9* | 2953 | FBgn0013812 | *Dhc93AB* |
| *DNAJB11* | 14889 | FBgn0031256 | *shv* |
| *DNAJB5* | 14887 | FBgn0031322 | *CG5001* |
| *DNAJB6* | 14888 | FBgn0034091 | *mrj* |
| *DNAJC13* | 30343 | FBgn0015477 | *Rme-8* |
| *DOCK1* | 2987 | FBgn0015513 | *mbc* |
| *DOCK4* | 19192 | FBgn0264324 | *spg* |
| *DOCK7* | 19190 | FBgn0031216 | *Zir* |
| *DOHH* | 28662 | FBgn0261479 | *nero* |
| *DOM3Z* | 2992 | FBgn0030793 | *CG9125* |
| *DPP6* | 3010 | FBgn0263780 | *CG17684* |
| *DPYSL2* | 3014 | FBgn0023023 | *CRMP* |
| *DPYSL3* | 3015 | FBgn0023023 | *CRMP* |
| *DST* | 1090 | FBgn0013733 | *shot* |
| *DUOX2* | 13273 | FBgn0283531 | *Duox* |
| *DUS1L* | 30086 | FBgn0031238 | *CG3645* |
| *DUSP14* | 17007 | FBgn0039742 | *CG15528* |
| *DYNC1H1* | 2961 | FBgn0261797 | *Dhc64C* |
| *DYSF* | 3097 | FBgn0266757 | *mfr* |
| *DYTN* | 23279 | FBgn0015926 | *dah* |
| *DZIP1L* | 26551 | FBgn0039201 | *CG13617* |
| *EBAG9* | 3123 | FBgn0052536 | *CG32536* |
| *EBF3* | 19087 | FBgn0001319 | *kn* |
| *ECD* | 17029 | FBgn0000543 | *ecd* |
| *ECE2* | 13275 | FBgn0031081 | *Nep3* |
| *ECHDC3* | 23489 | FBgn0034191 | *CG6984* |
| *EEF1A2* | 3192 | FBgn0284245 | *eEF1alpha1* |
| *EFNA2* | 3222 | FBgn0040324 | *Ephrin* |
| *EFR3A* | 28970 | FBgn0086784 | *stmA* |
| *EFR3B* | 29155 | FBgn0086784 | *stmA* |
| *EGFR* | 3236 | FBgn0003731 | *Egfr* |
| *EHD2* | 3243 | FBgn0016693 | *Past1* |
| *EIF2AK3* | 3255 | FBgn0037327 | *PEK* |

| | | | |
|---|---|---|---|
| *EIF3B* | 3280 | FBgn0034237 | *eIF3b* |
| *EIF3G* | 3274 | FBgn0038796 | *eIF3g2* |
| *EIF3J* | 3270 | FBgn0027619 | *eIF3j* |
| *EIF4A1* | 3282 | FBgn0001942 | *eIF4A* |
| *EIF4G1* | 3296 | FBgn0023213 | *eIF4G1* |
| *EIF4G2* | 3297 | FBgn0010488 | *NAT1* |
| *ELAVL3* | 3314 | FBgn0086675 | *fne* |
| *ELOVL1* | 14418 | FBgn0051522 | *CG31522* |
| *EMC4* | 28032 | FBgn0037199 | *CG11137* |
| *EME2* | 27289 | FBgn0033549 | *mms4* |
| *EP300* | 3373 | FBgn0261617 | *nej* |
| *EP400* | 11958 | FBgn0020306 | *dom* |
| *EPB42* | 3381 | FBgn0031975 | *Tg* |
| *EPHA1* | 3385 | FBgn0025936 | *Eph* |
| *EPHB1* | 3392 | FBgn0025936 | *Eph* |
| *EPRS* | 3418 | FBgn0005674 | *GluProRS* |
| *EPT1* | 29361 | FBgn0053116 | *CG33116* |
| *ERBB2IP* | 15842 | FBgn0033984 | *Lap1* |
| *ERP44* | 18311 | FBgn0030734 | *CG9911* |
| *ETV6* | 3495 | FBgn0000097 | *aop* |
| *EVL* | 20234 | FBgn0000578 | *ena* |
| *EXD2* | 20217 | FBgn0037901 | *Exd2* |
| *EXTL1* | 3515 | FBgn0265974 | *ttv* |
| *EYA1* | 3519 | FBgn0000320 | *eya* |
| *FAAH2* | 26440 | FBgn0033717 | *CG8839* |
| *FABP4* | 3559 | FBgn0037913 | *fabp* |
| *FAF2* | 24666 | FBgn0025608 | *Faf2* |
| *FAM136A* | 25911 | FBgn0034362 | *CG5323* |
| *FAM151A* | 25032 | FBgn0031968 | *CG7231* |
| *FAM177B* | 34395 | FBgn0029937 | *CG8300* |
| *FAM214B* | 25666 | FBgn0033638 | *CG9005* |
| *FAM92B* | 24781 | FBgn0032428 | *CG6405* |
| *FAP* | 3590 | FBgn0259175 | *ome* |
| *FAT1* | 3595 | FBgn0261574 | *kug* |
| *FAT2* | 3596 | FBgn0261574 | *kug* |
| *FAT3* | 23112 | FBgn0261574 | *kug* |
| *FBN3* | 18794 | FBgn0035798 | *frac* |
| *FBXL6* | 13603 | FBgn0033609 | *Fbl6* |
| *FBXO11* | 13590 | FBgn0037760 | *FBXO11* |
| *FBXO43* | 28521 | FBgn0017551 | *Rca1* |
| *FCGBP* | 13572 | FBgn0029167 | *Hml* |
| *FERMT1* | 15889 | FBgn0035498 | *Fit1* |
| *FEZF2* | 13506 | FBgn0031375 | *erm* |
| *FGD3* | 16027 | FBgn0035761 | *RhoGEF4* |
| *FGGY* | 25610 | FBgn0035484 | *CG11594* |

| | | | |
|---|---|---|---|
| *FLII* | 3750 | FBgn0000709 | *fliI* |
| *FNBP1L* | 20851 | FBgn0035533 | *Cip4* |
| *FOXP3* | 6106 | FBgn0262477 | *FoxP* |
| *FRMD3* | 24125 | FBgn0032225 | *CG5022* |
| *FRYL* | 29127 | FBgn0016081 | *fry* |
| *FSHB* | 3964 | FBgn0063368 | *Gpb5* |
| *FTSJ3* | 17136 | FBgn0030720 | *CG8939* |
| *G3BP2* | 30291 | FBgn0015778 | *rin* |
| *G6PC2* | 28906 | FBgn0031463 | *G6P* |
| *GAB3* | 17515 | FBgn0016794 | *dos* |
| *GABRA1* | 4075 | FBgn0030707 | *CG8916* |
| *GABRB2* | 4082 | FBgn0010240 | *Lcch3* |
| *GABRB3* | 4083 | FBgn0010240 | *Lcch3* |
| *GAPVD1* | 23375 | FBgn0030286 | *Gapvd1* |
| *GAS2L2* | 24846 | FBgn0029881 | *pigs* |
| *GATA4* | 4173 | FBgn0003117 | *pnr* |
| *GCLC* | 4311 | FBgn0040319 | *Gclc* |
| *GCM2* | 4198 | FBgn0014179 | *gcm* |
| *GCN1L1* | 4199 | FBgn0039959 | *CG17514* |
| *GDI1* | 4226 | FBgn0004868 | *Gdi* |
| *GGT7* | 4259 | FBgn0030796 | *CG4829* |
| *GLMN* | 14373 | FBgn0050496 | *CG30496* |
| *GLOD4* | 14111 | FBgn0031143 | *CG1532* |
| *GLRA2* | 4327 | FBgn0024963 | *GluClalpha* |
| *GLYCTK* | 24247 | FBgn0031428 | *CG9886* |
| *GNAI1* | 4384 | FBgn0001104 | *Galphai* |
| *GNAO1* | 4389 | FBgn0001122 | *Galphao* |
| *GNAS* | 4392 | FBgn0001123 | *Galphas* |
| *GNPDA2* | 21526 | FBgn0031717 | *Oscillin* |
| *GOLGA4* | 4427 | FBgn0034854 | *Golgin245* |
| *GON4L* | 25973 | FBgn0085444 | *mute* |
| *GORASP2* | 17500 | FBgn0036919 | *Grasp65* |
| *GPC5* | 4453 | FBgn0263930 | *dally* |
| *GPCPD1* | 26957 | FBgn0031566 | *CG2818* |
| *GPN1* | 17030 | FBgn0040346 | *CG3704* |
| *GPR112* | 18992 | FBgn0039818 | *CG11318* |
| *GPR179* | 31371 | FBgn0051760 | *CG31760* |
| *GPS1* | 4549 | FBgn0027057 | *CSN1b* |
| *GPSM1* | 17858 | FBgn0040080 | *pins* |
| *GRAMD1B* | 29214 | FBgn0085423 | *GramD1B* |
| *GRIA1* | 4571 | FBgn0264000 | *GluRIB* |
| *GRIK5* | 4583 | FBgn0039916 | *Ekar* |
| *GRIN2B* | 4586 | FBgn0053513 | *Nmdar2* |
| *GRK4* | 4543 | FBgn0261988 | *Gprk2* |
| *GRM5* | 4597 | FBgn0050361 | *mtt* |

| | | | |
|---|---|---|---|
| *GRM6* | 4598 | FBgn0019985 | *mGluR* |
| *GRM7* | 4599 | FBgn0019985 | *mGluR* |
| *GSAP* | 28042 | FBgn0010309 | *pigeon* |
| *GSKIP* | 20343 | FBgn0037156 | *CG11523* |
| *GTF3C1* | 4664 | FBgn0032517 | *CG7099* |
| *GUCY1A3* | 4685 | FBgn0013972 | *Gycalpha99B* |
| *GYG2* | 4700 | FBgn0265191 | *Gyg* |
| *H2AFV* | 20664 | FBgn0001197 | *His2Av* |
| *HADHA* | 4801 | FBgn0028479 | *Mtpalpha* |
| *HADHB* | 4803 | FBgn0025352 | *Thiolase* |
| *HCN2* | 4846 | FBgn0263397 | *Ih* |
| *HCN4* | 16882 | FBgn0263397 | *Ih* |
| *HDAC1* | 4852 | FBgn0015805 | *HDAC1* |
| *HDAC3* | 4854 | FBgn0025825 | *HDAC3* |
| *HDAC9* | 14065 | FBgn0041210 | *HDAC4* |
| *HDGFRP2* | 14680 | FBgn0039743 | *CG7946* |
| *HDLBP* | 4857 | FBgn0027835 | *Dp1* |
| *HECTD1* | 20157 | FBgn0032208 | *Ufd4* |
| *HECW2* | 29853 | FBgn0261931 | *CG42797* |
| *HELQ* | 18536 | FBgn0002899 | *mus301* |
| *HERC2* | 4868 | FBgn0031107 | *HERC2* |
| *HEXIM1* | 24953 | FBgn0038251 | *Hexim* |
| *HEY1* | 4880 | FBgn0027788 | *Hey* |
| *HIGD2A* | 28311 | FBgn0030743 | *CG9921* |
| *HIST1H2BJ* | 4761 | FBgn0053884 | *His2B:CG33884* |
| *HIST1H3G* | 4772 | FBgn0053845 | *His3:CG33845* |
| *HMGCS2* | 5008 | FBgn0010611 | *Hmgs* |
| *HNRNPF* | 5039 | FBgn0259139 | *glo* |
| *HNRNPUL1* | 17011 | FBgn0050122 | *CG30122* |
| *HNRNPUL2* | 25451 | FBgn0050122 | *CG30122* |
| *HPCA* | 5144 | FBgn0013303 | *Nca* |
| *HSD17B2* | 5211 | FBgn0262112 | *sro* |
| *HSD3B1* | 5217 | FBgn0036698 | *CG7724* |
| *HSP90AA1* | 5253 | FBgn0001233 | *Hsp83* |
| *HSPA4* | 5237 | FBgn0026418 | *Hsc70Cb* |
| *HSPB2* | 5247 | FBgn0001225 | *Hsp26* |
| *HTR1D* | 5289 | FBgn0004573 | *5-HT1B* |
| *ICA1* | 5343 | FBgn0037050 | *ICA69* |
| *IGF2R* | 5467 | FBgn0051072 | *Lerp* |
| *IKBKAP* | 5959 | FBgn0037926 | *Elp1* |
| *IKBKB* | 5960 | FBgn0024222 | *IKKbeta* |
| *INCENP* | 6058 | FBgn0260991 | *Incenp* |
| *INIP* | 24994 | FBgn0259720 | *CG42374* |
| *INPP5B* | 6077 | FBgn0023508 | *Ocrl* |
| *INSR* | 6091 | FBgn0283499 | *InR* |

| | | | |
|---|---|---|---|
| *INTS10* | 25548 | FBgn0035462 | *IntS10* |
| *INTS6* | 14879 | FBgn0261383 | *IntS6* |
| *INTS7* | 24484 | FBgn0036038 | *defl* |
| *IREB2* | 6115 | FBgn0024958 | *Irp-1A* |
| *IRF2BPL* | 14282 | FBgn0030400 | *Pits* |
| *ITGA2B* | 6138 | FBgn0001250 | *if* |
| *ITGA7* | 6143 | FBgn0004456 | *mew* |
| *ITGA8* | 6144 | FBgn0001250 | *if* |
| *ITGAV* | 6150 | FBgn0001250 | *if* |
| *ITGB3* | 6156 | FBgn0004657 | *mys* |
| *ITPR1* | 6180 | FBgn0010051 | *Itp-r83A* |
| *ITPR3* | 6182 | FBgn0010051 | *Itp-r83A* |
| *ITSN2* | 6184 | FBgn0023388 | *Dap160* |
| *JAK1* | 6190 | FBgn0004864 | *hop* |
| *JPH3* | 14203 | FBgn0032129 | *jp* |
| *JUP* | 6207 | FBgn0000117 | *arm* |
| *KANK1* | 19309 | FBgn0027596 | *Kank* |
| *KANSL1* | 24565 | FBgn0262527 | *nsl1* |
| *KAT6B* | 17582 | FBgn0034975 | *enok* |
| *KCNC1* | 6233 | FBgn0003386 | *Shaw* |
| *KCND3* | 6239 | FBgn0005564 | *Shal* |
| *KCNH3* | 6252 | FBgn0011589 | *Elk* |
| *KCNH8* | 18864 | FBgn0011589 | *Elk* |
| *KCNJ15* | 6261 | FBgn0032706 | *Irk3* |
| *KCNJ3* | 6264 | FBgn0039081 | *Irk2* |
| *KCTD20* | 21052 | FBgn0035107 | *mri* |
| *KCTD3* | 21305 | FBgn0037758 | *CG9467* |
| *KDELR2* | 6305 | FBgn0267330 | *KdelR* |
| *KDM1A* | 29079 | FBgn0260397 | *Su(var)3-3* |
| *KDM2A* | 13606 | FBgn0037659 | *Kdm2* |
| *KDM2B* | 13610 | FBgn0037659 | *Kdm2* |
| *KDM5B* | 18039 | FBgn0031759 | *lid* |
| *KDM6B* | 29012 | FBgn0260749 | *Utx* |
| *KDR* | 6307 | FBgn0032006 | *Pvr* |
| *KIAA0100* | 28960 | FBgn0035420 | *hob* |
| *KIAA0195* | 28983 | FBgn0022153 | *l(2)k05819* |
| *KIAA1009* | 21107 | FBgn0261610 | *CG42699* |
| *KIAA1161* | 19918 | FBgn0261575 | *tobi* |
| *KIAA1432* | 17686 | FBgn0028500 | *Rich* |
| *KIF13B* | 14405 | FBgn0019968 | *Khc-73* |
| *KIF18A* | 29441 | FBgn0004379 | *Klp67A* |
| *KIF1A* | 888 | FBgn0267002 | *unc-104* |
| *KIF21A* | 19349 | FBgn0032243 | *Klp31E* |
| *KIFC3* | 6326 | FBgn0002924 | *ncd* |
| *KIRREL2* | 18816 | FBgn0003285 | *rst* |

| | | | |
|---|---|---|---|
| *KIRREL3* | 23204 | FBgn0003285 | *rst* |
| *KLF5* | 6349 | FBgn0040765 | *luna* |
| *KLHL4* | 6355 | FBgn0030114 | *CG17754* |
| *KLK1* | 6357 | FBgn0037677 | *CG12951* |
| *KMT2C* | 13726 | FBgn0023518 | *trr* |
| *KMT2D* | 7133 | FBgn0023518 | *trr* |
| *L3MBTL1* | 15905 | FBgn0002441 | *l(3)mbt* |
| *LAMA1* | 6481 | FBgn0261563 | *wb* |
| *LAMA2* | 6482 | FBgn0261563 | *wb* |
| *LAMA3* | 6483 | FBgn0002526 | *LanA* |
| *LAMA4* | 6484 | FBgn0002526 | *LanA* |
| *LAMB1* | 6486 | FBgn0261800 | *LanB1* |
| *LAMB2* | 6487 | FBgn0261800 | *LanB1* |
| *LAMC1* | 6492 | FBgn0267348 | *LanB2* |
| *LAMC3* | 6494 | FBgn0267348 | *LanB2* |
| *LAMP3* | 14582 | FBgn0032949 | *Lamp1* |
| *LCA5* | 31923 | FBgn0036687 | *CG6652* |
| *LETM1* | 6556 | FBgn0284252 | *Letm1* |
| *LHFPL3* | 6589 | FBgn0262624 | *Tmhs* |
| *LIMK1* | 6613 | FBgn0283712 | *LIMK1* |
| *LIPT1* | 29569 | FBgn0265178 | *CG44243* |
| *LIPT2* | 37216 | FBgn0037251 | *CG9804* |
| *LIX1* | 18581 | FBgn0032230 | *lft* |
| *LLGL1* | 6628 | FBgn0002121 | *l(2)gl* |
| *LNX1* | 6657 | FBgn0001263 | *inaD* |
| *LPHN2* | 18582 | FBgn0033313 | *Cirl* |
| *LPIN2* | 14450 | FBgn0263593 | *Lpin* |
| *LRCH4* | 6691 | FBgn0032633 | *Lrch* |
| *LRP1* | 6692 | FBgn0053087 | *LRP1* |
| *LRP11* | 16936 | FBgn0027550 | *CG6495* |
| *LRP2* | 6694 | FBgn0261260 | *mgl* |
| *LRPAP1* | 6701 | FBgn0037756 | *CG8507* |
| *LRRC40* | 26004 | FBgn0028487 | *f-cup* |
| *LRRK1* | 18608 | FBgn0038816 | *Lrrk* |
| *LSG1* | 25652 | FBgn0266284 | *Ns3* |
| *LTK* | 6721 | FBgn0040505 | *Alk* |
| *LYSMD1* | 32070 | FBgn0285913 | *red* |
| *LZTR1* | 6742 | FBgn0040344 | *Lztr1* |
| *MACF1* | 13664 | FBgn0013733 | *shot* |
| *MADD* | 6766 | FBgn0030613 | *Rab3-GEF* |
| *MAGEA3* | 6801 | FBgn0037481 | *MAGE* |
| *MAGEA8* | 6806 | FBgn0037481 | *MAGE* |
| *MAGEC1* | 6812 | FBgn0037481 | *MAGE* |
| *MAGEE1* | 24934 | FBgn0037481 | *MAGE* |
| *MAN2B1* | 6826 | FBgn0027611 | *LManII* |

| | | | |
|---|---|---|---|
| *MANBA* | 6831 | FBgn0037215 | *beta-Man* |
| *MAP2K3* | 6843 | FBgn0261524 | *lic* |
| *MAP2K7* | 6847 | FBgn0010303 | *hep* |
| *MAP3K10* | 6849 | FBgn0030018 | *slpr* |
| *MAP4* | 6862 | FBgn0266579 | *tau* |
| *MAP4K1* | 6863 | FBgn0263395 | *hppy* |
| *MAPK13* | 6875 | FBgn0015765 | *p38a* |
| *MAPK8IP1* | 6882 | FBgn0040281 | *Aplip1* |
| *MAPK8IP3* | 6884 | FBgn0024187 | *syd* |
| *MARK3* | 6897 | FBgn0260934 | *par-1* |
| *MAST2* | 19035 | FBgn0267390 | *dop* |
| *MBD2* | 6917 | FBgn0027950 | *MBD-like* |
| *MBNL1* | 6923 | FBgn0265487 | *mbl* |
| *MCCC2* | 6937 | FBgn0042083 | *Mccc2* |
| *MCF2L2* | 30319 | FBgn0050440 | *CG30440* |
| *MCM6* | 6949 | FBgn0025815 | *Mcm6* |
| *MCOLN3* | 13358 | FBgn0262516 | *Trpml* |
| *MCTP2* | 25636 | FBgn0034389 | *Mctp* |
| *MDN1* | 18302 | FBgn0033661 | *CG13185* |
| *MED12L* | 16050 | FBgn0001324 | *kto* |
| *MED23* | 2372 | FBgn0034795 | *MED23* |
| *MED24* | 22963 | FBgn0035851 | *MED24* |
| *MEGF11* | 29635 | FBgn0027594 | *drpr* |
| *MEGF8* | 3233 | FBgn0031981 | *Megf8* |
| *METTL17* | 19280 | FBgn0032168 | *CG13126* |
| *MFN1* | 18262 | FBgn0029870 | *Marf* |
| *MGAT4B* | 7048 | FBgn0036446 | *CG9384* |
| *MIB1* | 21086 | FBgn0263601 | *mib1* |
| *MICAL2* | 24693 | FBgn0053208 | *Mical* |
| *MINK1* | 17565 | FBgn0010909 | *msn* |
| *MKL1* | 14334 | FBgn0052296 | *Mrtf* |
| *MKS1* | 7121 | FBgn0030395 | *Mks1* |
| *MLH1* | 7127 | FBgn0011659 | *Mlh1* |
| *MMP15* | 7161 | FBgn0035049 | *Mmp1* |
| *MMP3* | 7173 | FBgn0035049 | *Mmp1* |
| *MMP8* | 7175 | FBgn0035049 | *Mmp1* |
| *MMS22L* | 21475 | FBgn0023513 | *CG14803* |
| *MNS1* | 29636 | FBgn0037581 | *CG7352* |
| *MOCOS* | 18234 | FBgn0002641 | *mal* |
| *MPI* | 7216 | FBgn0286506 | *Mpi* |
| *MPO* | 7218 | FBgn0011828 | *Pxn* |
| *MPP6* | 18167 | FBgn0250785 | *vari* |
| *MROH7* | 24802 | FBgn0040236 | *c11.1* |
| *MRPL50* | 16654 | FBgn0028648 | *mRpL50* |
| *MRPS5* | 14498 | FBgn0044510 | *mRpS5* |

| | | | |
|---|---|---|---|
| *MRRF* | 7234 | FBgn0035980 | *mRRF1* |
| *MSH6* | 7329 | FBgn0036486 | *Msh6* |
| *MTERFD2* | 28785 | FBgn0031419 | *CG15390* |
| *MTF2* | 29535 | FBgn0003044 | *Pcl* |
| *MTMR2* | 7450 | FBgn0025742 | *mtm* |
| *MTMR8* | 16825 | FBgn0028497 | *CG3530* |
| *MTMR9* | 14596 | FBgn0035945 | *CG5026* |
| *MTOR* | 3942 | FBgn0021796 | *Tor* |
| *MTR* | 7468 | FBgn0032726 | *CG10621* |
| *MUC2* | 7512 | FBgn0029167 | *Hml* |
| *MUC5B* | 7516 | FBgn0029167 | *Hml* |
| *MUC6* | 7517 | FBgn0029167 | *Hml* |
| *MYB* | 7545 | FBgn0002914 | *Myb* |
| *MYBBP1A* | 7546 | FBgn0001341 | *Mybbp1A* |
| *MYH10* | 7568 | FBgn0265434 | *zip* |
| *MYH3* | 7573 | FBgn0264695 | *Mhc* |
| *MYH9* | 7579 | FBgn0265434 | *zip* |
| *MYO15A* | 7594 | FBgn0263705 | *Myo10A* |
| *MYO1E* | 7599 | FBgn0086347 | *Myo31DF* |
| *MYO5A* | 7602 | FBgn0261397 | *didum* |
| *MYO5C* | 7604 | FBgn0261397 | *didum* |
| *MYO7B* | 7607 | FBgn0000317 | *ck* |
| *MYOCD* | 16067 | FBgn0052296 | *Mrtf* |
| *MYT1L* | 7623 | FBgn0263772 | *CG43689* |
| *N4BP1* | 29850 | FBgn0259742 | *CG42360* |
| *NAA40* | 25845 | FBgn0039687 | *Naa40* |
| *NACA* | 7629 | FBgn0086904 | *Nacalpha* |
| *NAV2* | 15997 | FBgn0263873 | *sick* |
| *NBEAL2* | 31928 | FBgn0263110 | *CG43367* |
| *NCAPD2* | 24305 | FBgn0039680 | *Cap-D2* |
| *NCOA1* | 7668 | FBgn0041092 | *tai* |
| *NCOR1* | 7672 | FBgn0265523 | *Smr* |
| *NCOR2* | 7673 | FBgn0265523 | *Smr* |
| *NDRG3* | 14462 | FBgn0043070 | *MESK2* |
| *NDUFA12* | 23987 | FBgn0031436 | *ND-B17.2* |
| *NDUFA13* | 17194 | FBgn0029868 | *ND-B16.6* |
| *NF1* | 7765 | FBgn0015269 | *Nf1* |
| *NFASC* | 29866 | FBgn0264975 | *Nrg* |
| *NFIL3* | 7787 | FBgn0016076 | *vri* |
| *NFXL1* | 18726 | FBgn0035518 | *CG15011* |
| *NID2* | 13389 | FBgn0026403 | *Ndg* |
| *NISCH* | 18006 | FBgn0033996 | *CG11807* |
| *NLGN1* | 14291 | FBgn0083963 | *Nlg3* |
| *NLGN3* | 14289 | FBgn0083963 | *Nlg3* |
| *NMT2* | 7858 | FBgn0020392 | *Nmt* |

| | | | |
|---|---|---|---|
| *NOS3* | 7876 | FBgn0011676 | *Nos* |
| *NOTCH1* | 7881 | FBgn0004647 | *N* |
| *NOTCH3* | 7883 | FBgn0004647 | *N* |
| *NPFFR2* | 4525 | FBgn0038880 | *SIFaR* |
| *NR2F1* | 7975 | FBgn0003651 | *svp* |
| *NR4A2* | 7981 | FBgn0014859 | *Hr38* |
| *NSUN7* | 25857 | FBgn0266099 | *CG44836* |
| *NTN1* | 8029 | FBgn0015773 | *NetA* |
| *NTN5* | 25208 | FBgn0015774 | *NetB* |
| *NUAK1* | 14311 | FBgn0262617 | *Nuak1* |
| *NUB1* | 17623 | FBgn0031161 | *CG15445* |
| *NUDCD1* | 24306 | FBgn0030342 | *CG10347* |
| *NUP133* | 18016 | FBgn0039004 | *Nup133* |
| *NUP188* | 17859 | FBgn0033766 | *Nup188* |
| *NUP210* | 30052 | FBgn0266580 | *Gp210* |
| *NUP214* | 8064 | FBgn0010660 | *Nup214* |
| *OBSCN* | 15719 | FBgn0005666 | *bt* |
| *ONECUT1* | 8138 | FBgn0028996 | *onecut* |
| *OPLAH* | 8149 | FBgn0034733 | *CG4752* |
| *ORC4* | 8490 | FBgn0023181 | *Orc4* |
| *OTOGL* | 26901 | FBgn0029167 | *Hml* |
| *P4HA2* | 8547 | FBgn0039776 | *PH4alphaEFB* |
| *PACS2* | 23794 | FBgn0020647 | *KrT95D* |
| *PAK6* | 16061 | FBgn0025743 | *mbt* |
| *PAOX* | 20837 | FBgn0037606 | *CG8032* |
| *PAPD4* | 26776 | FBgn0260780 | *wisp* |
| *PAPL* | 33781 | FBgn0030245 | *CG1637* |
| *PAPOLA* | 14981 | FBgn0015949 | *hrg* |
| *PBRM1* | 30064 | FBgn0039227 | *polybromo* |
| *PBX4* | 13403 | FBgn0000611 | *exd* |
| *PC* | 8636 | FBgn0027580 | *PCB* |
| *PCDH15* | 14674 | FBgn0039709 | *Cad99C* |
| *PCNT* | 16068 | FBgn0086690 | *Plp* |
| *PCNX* | 19740 | FBgn0003048 | *pcx* |
| *PDE6C* | 8787 | FBgn0085370 | *Pde11* |
| *PDGFRB* | 8804 | FBgn0032006 | *Pvr* |
| *PDIA2* | 14180 | FBgn0286818 | *Pdi* |
| *PDIA4* | 30167 | FBgn0033663 | *ERp60* |
| *PDIA6* | 30168 | FBgn0025678 | *CaBP1* |
| *PDK2* | 8810 | FBgn0017558 | *Pdk* |
| *PDLIM1* | 2067 | FBgn0265991 | *Zasp52* |
| *PDS5A* | 29088 | FBgn0260012 | *pds5* |
| *PDSS2* | 23041 | FBgn0037044 | *Pdss2* |
| *PDZD2* | 18486 | FBgn0000008 | *a* |
| *PEAR1* | 33631 | FBgn0027594 | *drpr* |

| | | | |
|---|---|---|---|
| *PELI1* | 8827 | FBgn0025574 | *Pli* |
| *PEPD* | 8840 | FBgn0000455 | *Dip-C* |
| *PEX5L* | 30024 | FBgn0023516 | *Pex5* |
| *PGA5* | 8887 | FBgn0032049 | *Bace* |
| *PGM2* | 8906 | FBgn0033377 | *Pgm2a* |
| *PGRMC1* | 16090 | FBgn0030703 | *MSBP* |
| *PHACTR2* | 20956 | FBgn0052264 | *CG32264* |
| *PHF19* | 24566 | FBgn0003044 | *Pcl* |
| *PHF3* | 8921 | FBgn0082831 | *pps* |
| *PHIP* | 15673 | FBgn0011785 | *BRWD3* |
| *PHRF1* | 24351 | FBgn0037344 | *CG2926* |
| *PIAS4* | 17002 | FBgn0003612 | *Su(var)2-10* |
| *PIEZO2* | 26270 | FBgn0264953 | *Piezo* |
| *PIK3C2A* | 8971 | FBgn0015278 | *Pi3K68D* |
| *PIK3R2* | 8980 | FBgn0020622 | *Pi3K21B* |
| *PIKFYVE* | 23785 | FBgn0028741 | *fab1* |
| *PITPNM3* | 21043 | FBgn0003218 | *rdgB* |
| *PITX1* | 9004 | FBgn0020912 | *Ptx1* |
| *PIWIL4* | 18444 | FBgn0004872 | *piwi* |
| *PKM* | 9021 | FBgn0267385 | *PyK* |
| *PKN1* | 9405 | FBgn0020621 | *Pkn* |
| *PLCG2* | 9066 | FBgn0003416 | *sl* |
| *PLD4* | 23792 | FBgn0032923 | *CG9248* |
| *PLD5* | 26879 | FBgn0032923 | *CG9248* |
| *PLEKHA8* | 30037 | FBgn0030641 | *CG6299* |
| *PLEKHG2* | 29515 | FBgn0050115 | *GEFmeso* |
| *PLOD2* | 9082 | FBgn0036147 | *Plod* |
| *PLOD3* | 9083 | FBgn0036147 | *Plod* |
| *PLXDC1* | 20945 | FBgn0028331 | *l(1)G0289* |
| *PLXNA2* | 9100 | FBgn0025741 | *PlexA* |
| *PLXNB1* | 9103 | FBgn0025740 | *PlexB* |
| *PLXNB2* | 9104 | FBgn0025740 | *PlexB* |
| *PLXNB3* | 9105 | FBgn0025740 | *PlexB* |
| *PM20D1* | 26518 | FBgn0039052 | *CG6733* |
| *PNISR* | 21222 | FBgn0051211 | *CG31211* |
| *PNPLA7* | 24768 | FBgn0003656 | *sws* |
| *POGLUT1* | 22954 | FBgn0086253 | *rumi* |
| *POGZ* | 18801 | FBgn0033998 | *row* |
| *POLA2* | 30073 | FBgn0005696 | *DNApol-alpha73* |
| *POLD1* | 9175 | FBgn0263600 | *DNApol-delta* |
| *POLG* | 9179 | FBgn0004406 | *tam* |
| *POLQ* | 9186 | FBgn0002905 | *mus308* |
| *POLR2M* | 14862 | FBgn0038692 | *Gdn1* |
| *POU3F4* | 9217 | FBgn0086680 | *vvl* |
| *POU6F1* | 9224 | FBgn0261588 | *pdm3* |

| | | | |
|---|---|---|---|
| *PPAP2C* | 9230 | FBgn0041087 | *wun2* |
| *PPARGC1B* | 30022 | FBgn0037248 | *srl* |
| *PPFIA1* | 9245 | FBgn0046704 | *Liprin-alpha* |
| *PPIP5K2* | 29035 | FBgn0027279 | *l(1)G0196* |
| *PPM1A* | 9275 | FBgn0086361 | *alph* |
| *PPM1B* | 9276 | FBgn0086361 | *alph* |
| *PPP1R3A* | 9291 | FBgn0036862 | *Gbs-76A* |
| *PPP1R9A* | 14946 | FBgn0010905 | *Spn* |
| *PPP2R1B* | 9303 | FBgn0260439 | *Pp2A-29B* |
| *PPP2R5D* | 9312 | FBgn0042693 | *wrd* |
| *PPP5C* | 9322 | FBgn0005777 | *PpD3* |
| *PPP6R2* | 19253 | FBgn0035688 | *fmt* |
| *PPRC1* | 30025 | FBgn0037248 | *srl* |
| *PRCP* | 9344 | FBgn0032864 | *CG2493* |
| *PRKAA1* | 9376 | FBgn0023169 | *AMPKalpha* |
| *PRKAR1B* | 9390 | FBgn0259243 | *Pka-R1* |
| *PRKCA* | 9393 | FBgn0003091 | *Pkc53E* |
| *PRKD1* | 9407 | FBgn0038603 | *PKD* |
| *PRMT8* | 5188 | FBgn0037834 | *Art1* |
| *PRPF18* | 17351 | FBgn0027784 | *Prp18* |
| *PRPF8* | 17340 | FBgn0033688 | *Prp8* |
| *PRPS1L1* | 9463 | FBgn0036030 | *CG6767* |
| *PRPSAP2* | 9467 | FBgn0039790 | *CG2246* |
| *PRRC2C* | 24903 | FBgn0261710 | *nocte* |
| *PRUNE2* | 25209 | FBgn0035488 | *CG11593* |
| *PSD4* | 19096 | FBgn0051158 | *Efa6* |
| *PSEN1* | 9508 | FBgn0284421 | *Psn* |
| *PSMC4* | 9551 | FBgn0028686 | *Rpt3* |
| *PSMC5* | 9552 | FBgn0020369 | *Rpt6* |
| *PSMD8* | 9566 | FBgn0028693 | *Rpn12* |
| *PSMG4* | 21108 | FBgn0033781 | *CG13319* |
| *PTAR1* | 30449 | FBgn0027296 | *temp* |
| *PTEN* | 9588 | FBgn0026379 | *Pten* |
| *PTGES* | 9599 | FBgn0053178 | *CG33178* |
| *PTK7* | 9618 | FBgn0004839 | *otk* |
| *PTPLAD1* | 24175 | FBgn0032524 | *Hacd2* |
| *PTPN11* | 9644 | FBgn0000382 | *csw* |
| *PTPN7* | 9659 | FBgn0016641 | *PTP-ER* |
| *PTPRF* | 9670 | FBgn0000464 | *Lar* |
| *PTPRK* | 9674 | FBgn0267486 | *Ptp36E* |
| *PTPRZ1* | 9685 | FBgn0004369 | *Ptp99A* |
| *PUS7* | 26033 | FBgn0035901 | *Pus7* |
| *PXDN* | 14966 | FBgn0011828 | *Pxn* |
| *PXDNL* | 26359 | FBgn0011828 | *Pxn* |
| *QTRTD1* | 25771 | FBgn0036000 | *CG3434* |

| | | | |
|---|---|---|---|
| *RAB43* | 19983 | FBgn0015793 | *Rab19* |
| *RAD51* | 9817 | FBgn0003479 | *spn-A* |
| *RAD9B* | 21700 | FBgn0025807 | *Rad9* |
| *RALGAPA1* | 17770 | FBgn0039466 | *CG5521* |
| *RAN* | 9846 | FBgn0020255 | *Ran* |
| *RANBP17* | 14428 | FBgn0053180 | *Ranbp16* |
| *RANBP2* | 9848 | FBgn0039302 | *Nup358* |
| *RAPGEF4* | 16626 | FBgn0085421 | *Epac* |
| *RASAL3* | 26129 | FBgn0261570 | *CG42684* |
| *RBM12* | 9898 | FBgn0035235 | *CG7879* |
| *RBM27* | 29243 | FBgn0002044 | *swm* |
| *RBM8A* | 9905 | FBgn0033378 | *tsu* |
| *RBMS3* | 13427 | FBgn0052423 | *shep* |
| *RCOR2* | 27455 | FBgn0261573 | *CoRest* |
| *RFC5* | 9973 | FBgn0032244 | *RfC3* |
| *RFWD3* | 25539 | FBgn0036660 | *CG13025* |
| *RFX2* | 9983 | FBgn0020379 | *Rfx* |
| *RFX7* | 25777 | FBgn0037445 | *CG9727* |
| *RGL1* | 30281 | FBgn0026376 | *Rgl* |
| *RGN* | 9989 | FBgn0038257 | *smp-30* |
| *RHOT2* | 21169 | FBgn0039140 | *Miro* |
| *RHPN2* | 19974 | FBgn0026374 | *Rhp* |
| *RIC8A* | 29550 | FBgn0028292 | *ric8a* |
| *RIC8B* | 25555 | FBgn0028292 | *ric8a* |
| *RICTOR* | 28611 | FBgn0031006 | *rictor* |
| *RIMS2* | 17283 | FBgn0053547 | *Rim* |
| *RIN2* | 18750 | FBgn0085443 | *spri* |
| *RLTPR* | 27089 | FBgn0033212 | *LRR* |
| *RNF123* | 21148 | FBgn0038296 | *CG6752* |
| *RNF151* | 23235 | FBgn0028847 | *CG9014* |
| *RNF25* | 14662 | FBgn0033884 | *CG13344* |
| *RNF4* | 10067 | FBgn0037384 | *dgrn* |
| *ROBO4* | 17985 | FBgn0002543 | *robo2* |
| *RORB* | 10259 | FBgn0000448 | *Hr3* |
| *ROS1* | 10261 | FBgn0003366 | *sev* |
| *RPF1* | 30350 | FBgn0032408 | *CG6712* |
| *RPH3A* | 17056 | FBgn0030230 | *Rph* |
| *RPL21* | 10313 | FBgn0032987 | *RpL21* |
| *RPS6KA1* | 10430 | FBgn0262866 | *S6kII* |
| *RPTOR* | 30287 | FBgn0029840 | *raptor* |
| *RRAGC* | 19902 | FBgn0033272 | *RagC-D* |
| *RRP8* | 29030 | FBgn0034422 | *CG7137* |
| *RTCA* | 17981 | FBgn0025630 | *Rtca* |
| *RUFY2* | 19761 | FBgn0051064 | *CG31064* |
| *RUVBL1* | 10474 | FBgn0040078 | *pont* |

| | | | |
|---|---|---|---|
| *RYR1* | 10483 | FBgn0011286 | *RyR* |
| *RYR2* | 10484 | FBgn0011286 | *RyR* |
| *RYR3* | 10485 | FBgn0011286 | *RyR* |
| *SASH1* | 19182 | FBgn0051163 | *SKIP* |
| *SBF1* | 10542 | FBgn0025802 | *Sbf* |
| *SBF2* | 2135 | FBgn0025802 | *Sbf* |
| *SCARB2* | 1665 | FBgn0010435 | *emp* |
| *SCN11A* | 10583 | FBgn0285944 | *para* |
| *SCN1A* | 10585 | FBgn0285944 | *para* |
| *SCN2A* | 10588 | FBgn0285944 | *para* |
| *SCN3A* | 10590 | FBgn0285944 | *para* |
| *SCN4A* | 10591 | FBgn0285944 | *para* |
| *SCN5A* | 10593 | FBgn0285944 | *para* |
| *SDC3* | 10660 | FBgn0010415 | *Sdc* |
| *SDHA* | 10680 | FBgn0261439 | *SdhA* |
| *SDK1* | 19307 | FBgn0021764 | *sdk* |
| *SDK2* | 19308 | FBgn0021764 | *sdk* |
| *SEC14L5* | 29032 | FBgn0031814 | *retm* |
| *SEC16B* | 30301 | FBgn0052654 | *Sec16* |
| *SEC24A* | 10703 | FBgn0033460 | *Sec24AB* |
| *SEC24D* | 10706 | FBgn0262126 | *Sec24CD* |
| *SEMA4C* | 10731 | FBgn0011260 | *Sema2a* |
| *SEMA4G* | 10735 | FBgn0011260 | *Sema2a* |
| *SEMA5A* | 10736 | FBgn0284221 | *Sema5c* |
| *SESN2* | 20746 | FBgn0034897 | *Sesn* |
| *SETBP1* | 15573 | FBgn0005386 | *ash1* |
| *SETD1B* | 29187 | FBgn0040022 | *Set1* |
| *SETD2* | 18420 | FBgn0030486 | *Set2* |
| *SETD5* | 25566 | FBgn0036398 | *upSET* |
| *SETX* | 445 | FBgn0035842 | *CG7504* |
| *SF1* | 12950 | FBgn0025571 | *SF1* |
| *SF3B1* | 10768 | FBgn0031266 | *Sf3b1* |
| *SFPQ* | 10774 | FBgn0004227 | *nonA* |
| *SGSM1* | 29410 | FBgn0052506 | *CG32506* |
| *SGSM3* | 25228 | FBgn0038304 | *CG12241* |
| *SH2D3C* | 16884 | FBgn0031762 | *CG9098* |
| *SH3RF3* | 24699 | FBgn0040294 | *POSH* |
| *SHOX* | 10853 | FBgn0085396 | *CG34367* |
| *SHPRH* | 19336 | FBgn0035689 | *CG7376* |
| *SIK1* | 11142 | FBgn0025625 | *Sik2* |
| *SIN3A* | 19353 | FBgn0022764 | *Sin3A* |
| *SIN3B* | 19354 | FBgn0022764 | *Sin3A* |
| *SKIL* | 10897 | FBgn0085450 | *Snoo* |
| *SLC12A4* | 10913 | FBgn0261794 | *kcc* |
| *SLC12A9* | 17435 | FBgn0032689 | *CG10413* |

| | | | |
|---|---|---|---|
| *SLC16A10* | 17027 | FBgn0001296 | *kar* |
| *SLC16A3* | 10924 | FBgn0023549 | *Mct1* |
| *SLC16A5* | 10926 | FBgn0023549 | *Mct1* |
| *SLC17A1* | 10929 | FBgn0039886 | *CG2003* |
| *SLC17A3* | 10931 | FBgn0010497 | *dmGlut* |
| *SLC22A11* | 18120 | FBgn0086365 | *Orct2* |
| *SLC22A24* | 28542 | FBgn0038719 | *CG16727* |
| *SLC22A6* | 10970 | FBgn0019952 | *Orct* |
| *SLC22A9* | 16261 | FBgn0264907 | *CG44098* |
| *SLC23A1* | 10974 | FBgn0037807 | *CG6293* |
| *SLC25A29* | 20116 | FBgn0032219 | *CG4995* |
| *SLC26A5* | 9359 | FBgn0036770 | *Prestin* |
| *SLC26A7* | 14467 | FBgn0036770 | *Prestin* |
| *SLC2A10* | 13444 | FBgn0029932 | *CG4607* |
| *SLC2A9* | 13446 | FBgn0028561 | *sut3* |
| *SLC30A5* | 19089 | FBgn0037875 | *ZnT86D* |
| *SLC35C2* | 17117 | FBgn0035449 | *CG14971* |
| *SLC35E1* | 20803 | FBgn0031183 | *CG14621* |
| *SLC39A12* | 20860 | FBgn0036461 | *Zip71B* |
| *SLC41A3* | 31046 | FBgn0053181 | *CG33181* |
| *SLC4A1* | 11027 | FBgn0036043 | *CG8177* |
| *SLC4A2* | 11028 | FBgn0036043 | *CG8177* |
| *SLC4A4* | 11030 | FBgn0259111 | *Ndae1* |
| *SLC6A1* | 11042 | FBgn0039915 | *Gat* |
| *SLC6A13* | 11046 | FBgn0039915 | *Gat* |
| *SLC6A8* | 11055 | FBgn0011603 | *ine* |
| *SLC8A2* | 11069 | FBgn0013995 | *Calx* |
| *SLC9A3* | 11073 | FBgn0040297 | *Nhe2* |
| *SLCO1C1* | 13819 | FBgn0036732 | *Oatp74D* |
| *SLCO4A1* | 10953 | FBgn0051634 | *Oatp26F* |
| *SMAD2* | 6768 | FBgn0025800 | *Smox* |
| *SMAD4* | 6770 | FBgn0011655 | *Med* |
| *SMC3* | 2468 | FBgn0015615 | *SMC3* |
| *SMC6* | 20466 | FBgn0266282 | *jnj* |
| *SMG6* | 17809 | FBgn0039260 | *Smg6* |
| *SMYD1* | 20986 | FBgn0011566 | *Smyd3* |
| *SNAP91* | 14986 | FBgn0086372 | *lap* |
| *SNAPC3* | 11136 | FBgn0260398 | *Pbp49* |
| *SND1* | 30646 | FBgn0035121 | *Tudor-SN* |
| *SNRK* | 30598 | FBgn0033915 | *CG8485* |
| *SNX5* | 14969 | FBgn0032005 | *Snx6* |
| *SOGA2* | 29121 | FBgn0031869 | *CG18304* |
| *SOGA3* | 21494 | FBgn0031869 | *CG18304* |
| *SON* | 11183 | FBgn0037716 | *Son* |
| *SOX10* | 11190 | FBgn0024288 | *Sox100B* |

| | | | |
|---|---|---|---|
| *SOX6* | 16421 | FBgn0039938 | *Sox102F* |
| *SP3* | 11208 | FBgn0039169 | *Spps* |
| *SP7* | 17321 | FBgn0020378 | *Sp1* |
| *SPAG9* | 14524 | FBgn0024187 | *syd* |
| *SPATA5* | 18119 | FBgn0032450 | *CG5776* |
| *SPECC1L* | 29022 | FBgn0025633 | *CG13366* |
| *SPEG* | 16901 | FBgn0053519 | *Unc-89* |
| *SPEN* | 17575 | FBgn0016977 | *spen* |
| *SPG11* | 11226 | FBgn0034786 | *CG13531* |
| *SPTAN1* | 11273 | FBgn0250789 | *alpha-Spec* |
| *SPTBN1* | 11275 | FBgn0250788 | *beta-Spec* |
| *SRBD1* | 25521 | FBgn0051156 | *CG31156* |
| *SRCAP* | 16974 | FBgn0020306 | *dom* |
| *SRF* | 11291 | FBgn0004101 | *bs* |
| *SRPR* | 11307 | FBgn0010391 | *Gtp-bp* |
| *SRRM2* | 16639 | FBgn0035253 | *CG7971* |
| *SSPO* | 21998 | FBgn0029167 | *Hml* |
| *SSR1* | 11323 | FBgn0028327 | *l(1)G0320* |
| *SSRP1* | 11327 | FBgn0010278 | *Ssrp* |
| *STAG1* | 11354 | FBgn0020616 | *SA* |
| *STAG3* | 11356 | FBgn0020616 | *SA* |
| *STAT1* | 11362 | FBgn0016917 | *Stat92E* |
| *STK11* | 11389 | FBgn0038167 | *Lkb1* |
| *STK36* | 17209 | FBgn0001079 | *fu* |
| *STRIP2* | 22209 | FBgn0035437 | *Strip* |
| *STT3A* | 6172 | FBgn0031149 | *Stt3A* |
| *STUB1* | 11427 | FBgn0027052 | *STUB1* |
| *SULF2* | 20392 | FBgn0040271 | *Sulf1* |
| *SUPT16H* | 11465 | FBgn0002183 | *dre4* |
| *SUV420H1* | 24283 | FBgn0025639 | *Hmt4-20* |
| *SV2A* | 20566 | FBgn0029896 | *CG3168* |
| *SV2B* | 16874 | FBgn0029896 | *CG3168* |
| *SYMPK* | 22935 | FBgn0037371 | *Sym* |
| *SYNE1* | 17089 | FBgn0261836 | *Msp300* |
| *SYNGAP1* | 11497 | FBgn0261570 | *CG42684* |
| *SYT14* | 23143 | FBgn0261086 | *Syt14* |
| *SYT7* | 11514 | FBgn0039900 | *Syt7* |
| *TACO1* | 24316 | FBgn0032205 | *CG4957* |
| *TAF6* | 11540 | FBgn0010417 | *Taf6* |
| *TAF9* | 11542 | FBgn0000617 | *e(y)1* |
| *TANC2* | 30212 | FBgn0041096 | *rols* |
| *TAOK3* | 18133 | FBgn0031030 | *Tao* |
| *TATDN2* | 28988 | FBgn0038877 | *CG3308* |
| *TBC1D22B* | 21602 | FBgn0038855 | *CG5745* |
| *TBC1D25* | 8092 | FBgn0034009 | *CG8155* |

| | | | |
|---|---|---|---|
| *TBC1D31* | 30888 | FBgn0035073 | *CG16896* |
| *TBC1D8B* | 24715 | FBgn0037074 | *CG7324* |
| *TBCB* | 1989 | FBgn0034451 | *TBCB* |
| *TBCK* | 28261 | FBgn0029736 | *CG4041* |
| *TBL1XR1* | 29529 | FBgn0263933 | *ebi* |
| *TCERG1L* | 23533 | FBgn0261641 | *CG42724* |
| *TCF3* | 11633 | FBgn0267821 | *da* |
| *TCF7L1* | 11640 | FBgn0085432 | *pan* |
| *TDP1* | 18884 | FBgn0260817 | *gkt* |
| *TECRL* | 27365 | FBgn0035471 | *Sc2* |
| *TEFM* | 26223 | FBgn0037184 | *CG14450* |
| *TEKT1* | 15534 | FBgn0035638 | *Tektin-C* |
| *TENM2* | 29943 | FBgn0004449 | *Ten-m* |
| *TENM4* | 29945 | FBgn0004449 | *Ten-m* |
| *TET1* | 29484 | FBgn0263392 | *Tet* |
| *TET2* | 25941 | FBgn0263392 | *Tet* |
| *TFEB* | 11753 | FBgn0263112 | *Mitf* |
| *TGM3* | 11779 | FBgn0031975 | *Tg* |
| *TH* | 11782 | FBgn0005626 | *ple* |
| *THBS3* | 11787 | FBgn0031850 | *Tsp* |
| *THSD4* | 25835 | FBgn0032252 | *loh* |
| *TIAL1* | 11804 | FBgn0005649 | *Rox8* |
| *TIPIN* | 30750 | FBgn0032698 | *CG10336* |
| *TKTL2* | 25313 | FBgn0037607 | *CG8036* |
| *TLE3* | 11839 | FBgn0001139 | *gro* |
| *TLE4* | 11840 | FBgn0001139 | *gro* |
| *TLK2* | 11842 | FBgn0283657 | *Tlk* |
| *TLN2* | 15447 | FBgn0260442 | *rhea* |
| *TLR10* | 15634 | FBgn0036978 | *Toll-9* |
| *TMEM120A* | 21697 | FBgn0040384 | *CG32795* |
| *TMEM201* | 33719 | FBgn0034447 | *CG7744* |
| *TMEM214* | 25983 | FBgn0053129 | *CG33129* |
| *TMEM8A* | 17205 | FBgn0039290 | *CG13654* |
| *TMF1* | 11870 | FBgn0029912 | *CG4557* |
| *TNIK* | 30765 | FBgn0010909 | *msn* |
| *TNKS* | 11941 | FBgn0027508 | *Tnks* |
| *TNPO3* | 17103 | FBgn0031456 | *Tnpo-SR* |
| *TOP1MT* | 29787 | FBgn0004924 | *Top1* |
| *TOP3B* | 11993 | FBgn0026015 | *Top3beta* |
| *TPK1* | 17358 | FBgn0037942 | *CG14721* |
| *TPR* | 12017 | FBgn0013756 | *Mtor* |
| *TPST1* | 12020 | FBgn0086674 | *Tpst* |
| *TRAK1* | 29947 | FBgn0262872 | *milt* |
| *TRAK2* | 13206 | FBgn0262872 | *milt* |
| *TRAPPC10* | 11868 | FBgn0038303 | *SIDL* |

| | | | |
|---|---|---|---|
| *TRAPPC11* | 25751 | FBgn0286567 | *gry* |
| *TRAPPC9* | 30832 | FBgn0261787 | *brun* |
| *TRDMT1* | 2977 | FBgn0028707 | *Mt2* |
| *TREH* | 12266 | FBgn0003748 | *Treh* |
| *TRIO* | 12303 | FBgn0024277 | *trio* |
| *TRIP11* | 12305 | FBgn0027287 | *Gmap* |
| *TRIP12* | 12306 | FBgn0260794 | *ctrip* |
| *TRPM1* | 7146 | FBgn0265194 | *Trpm* |
| *TRPM6* | 17995 | FBgn0265194 | *Trpm* |
| *TRPM7* | 17994 | FBgn0265194 | *Trpm* |
| *TRPV4* | 18083 | FBgn0036414 | *nan* |
| *TRPV5* | 3145 | FBgn0036414 | *nan* |
| *TRRAP* | 12347 | FBgn0053554 | *Nipped-A* |
| *TSC2* | 12363 | FBgn0005198 | *gig* |
| *TSNARE1* | 26437 | FBgn0036341 | *Syx13* |
| *TSPAN4* | 11859 | FBgn0029837 | *Tsp5D* |
| *TSPYL5* | 29367 | FBgn0014879 | *Set* |
| *TSR2* | 25455 | FBgn0039404 | *CG14543* |
| *TSSK2* | 11401 | FBgn0038630 | *CG14305* |
| *TTC17* | 25596 | FBgn0029713 | *CG11436* |
| *TTLL5* | 19963 | FBgn0051108 | *TTLL5* |
| *TTN* | 12403 | FBgn0005666 | *bt* |
| *TUBA1A* | 20766 | FBgn0003886 | *alphaTub85E* |
| *TUBB4B* | 20771 | FBgn0284243 | *betaTub56D* |
| *TUBB8* | 20773 | FBgn0003890 | *betaTub97EF* |
| *TUBGCP4* | 16691 | FBgn0026431 | *Grip75* |
| *TUBGCP5* | 18600 | FBgn0026433 | *Grip128* |
| *TULP4* | 15530 | FBgn0039530 | *Tusp* |
| *TXLNA* | 30685 | FBgn0039379 | *CG5886* |
| *TXNDC5* | 21073 | FBgn0030329 | *prtp* |
| *TXNRD1* | 12437 | FBgn0037170 | *Trxr-2* |
| *U2SURP* | 30855 | FBgn0034572 | *CG9346* |
| *UBB* | 12463 | FBgn0003943 | *Ubi-p63E* |
| *UBE3A* | 12496 | FBgn0061469 | *Ube3a* |
| *UBE3C* | 16803 | FBgn0034989 | *CG3356* |
| *UBE4A* | 12499 | FBgn0028467 | *CG11070* |
| *UBQLN3* | 12510 | FBgn0031057 | *Ubqn* |
| *UBR3* | 30467 | FBgn0260970 | *Ubr3* |
| *UBR4* | 30313 | FBgn0011230 | *poe* |
| *UBXN2A* | 27265 | FBgn0033179 | *p47* |
| *UBXN6* | 14928 | FBgn0034372 | *Gint3* |
| *UGGT1* | 15663 | FBgn0014075 | *Uggt* |
| *UNC45B* | 14304 | FBgn0010812 | *unc-45* |
| *UNC80* | 26582 | FBgn0039536 | *unc80* |
| *UPF3A* | 20332 | FBgn0034923 | *Upf3* |

| | | | |
|---|---|---|---|
| *UQCRC2* | 12586 | FBgn0250814 | *UQCR-C2* |
| *URB1* | 17344 | FBgn0038968 | *CG12499* |
| *USP24* | 12623 | FBgn0005632 | *faf* |
| *USP30* | 20065 | FBgn0029819 | *Usp30* |
| *USP34* | 20066 | FBgn0039214 | *puf* |
| *USP45* | 20080 | FBgn0029763 | *Usp16-45* |
| *USP46* | 20075 | FBgn0039025 | *Usp12-46* |
| *USP7* | 12630 | FBgn0030366 | *Usp7* |
| *USP9X* | 12632 | FBgn0005632 | *faf* |
| *UTP20* | 17897 | FBgn0034734 | *CG4554* |
| *VNN1* | 12705 | FBgn0052750 | *CG32750* |
| *VPS13A* | 1908 | FBgn0033194 | *Vps13* |
| *VPS13D* | 23595 | FBgn0052113 | *Vps13D* |
| *VPS18* | 15972 | FBgn0000482 | *dor* |
| *VPS25* | 28122 | FBgn0022027 | *Vps25* |
| *VPS39* | 20593 | FBgn0038593 | *Vps39* |
| *VPS4A* | 13488 | FBgn0283469 | *Vps4* |
| *WDFY3* | 20751 | FBgn0043362 | *bchs* |
| *WDFY4* | 29323 | FBgn0043362 | *bchs* |
| *WDR26* | 21208 | FBgn0037094 | *CG7611* |
| *WDR37* | 31406 | FBgn0038617 | *CG12333* |
| *WDR4* | 12756 | FBgn0029857 | *wuho* |
| *WDR45* | 28912 | FBgn0037648 | *CG11975* |
| *WDR55* | 25971 | FBgn0037943 | *CG14722* |
| *XPNPEP2* | 12823 | FBgn0038072 | *CG6225* |
| *XRN2* | 12836 | FBgn0031868 | *Rat1* |
| *XYLT2* | 15517 | FBgn0015360 | *oxt* |
| *YIPF5* | 24877 | FBgn0032465 | *Yip1d1* |
| *YTHDC1* | 30626 | FBgn0027616 | *Ythdc1* |
| *YTHDC2* | 24721 | FBgn0030833 | *CG8915* |
| *ZC3H12B* | 17407 | FBgn0038769 | *CG10889* |
| *ZC3H3* | 28972 | FBgn0035900 | *ZC3H3* |
| *ZC3H4* | 17808 | FBgn0003575 | *su(sable)* |
| *ZDHHC22* | 20106 | FBgn0042133 | *CG18810* |
| *ZEB2* | 14881 | FBgn0004606 | *zfh1* |
| *ZFHX3* | 777 | FBgn0004607 | *zfh2* |
| *ZFPM2* | 16700 | FBgn0003963 | *ush* |
| *ZFYVE26* | 20761 | FBgn0037897 | *CG5270* |
| *ZFYVE9* | 6775 | FBgn0026369 | *Sara* |
| *ZMAT5* | 28046 | FBgn0051922 | *CG31922* |
| *ZMIZ2* | 22229 | FBgn0026160 | *tna* |
| *ZMYND8* | 9397 | FBgn0039863 | *CG1815* |
| *ZRANB3* | 25249 | FBgn0031655 | *Marcal1* |
| *ZSWIM8* | 23528 | FBgn0085430 | *CG34401* |

in *Drosophila mela*

| DIOPT Score |
|---|
| 6 |
| 5 |
| 15 |
| 4 |
| 7 |
| 4 |
| 4 |
| 4 |
| 15 |
| 14 |
| 6 |
| 10 |
| 8 |
| 12 |
| 14 |
| 9 |
| 10 |
| 13 |
| 13 |
| 10 |
| 9 |
| 13 |
| 11 |
| 15 |
| 5 |
| 5 |
| 11 |
| 9 |
| 10 |
| 11 |
| 11 |
| 15 |
| 6 |
| 13 |
| 9 |
| 14 |
| 12 |
| 12 |
| 13 |
| 11 |
| 5 |
| 10 |
| 5 |

| |
|---|
| 8 |
| 13 |
| 15 |
| 14 |
| 9 |
| 15 |
| 12 |
| 13 |
| 13 |
| 5 |
| 7 |
| 7 |
| 10 |
| 7 |
| 8 |
| 6 |
| 12 |
| 14 |
| 12 |
| 13 |
| 7 |
| 9 |
| 8 |
| 4 |
| 5 |
| 13 |
| 7 |
| 4 |
| 10 |
| 10 |
| 5 |
| 4 |
| 12 |
| 13 |
| 9 |
| 9 |
| 7 |
| 12 |
| 11 |
| 10 |
| 14 |
| 15 |
| 12 |
| 10 |
| 6 |

| |
|---|
| 7 |
| 14 |
| 12 |
| 13 |
| 5 |
| 13 |
| 12 |
| 12 |
| 8 |
| 13 |
| 10 |
| 14 |
| 10 |
| 6 |
| 12 |
| 15 |
| 6 |
| 11 |
| 9 |
| 13 |
| 8 |
| 9 |
| 14 |
| 14 |
| 13 |
| 12 |
| 4 |
| 14 |
| 5 |
| 6 |
| 11 |
| 14 |
| 11 |
| 8 |
| 12 |
| 13 |
| 9 |
| 9 |
| 9 |
| 13 |
| 5 |
| 14 |
| 9 |
| 12 |
| 12 |

| |
|---|
| 4 |
| 5 |
| 5 |
| 15 |
| 15 |
| 11 |
| 14 |
| 9 |
| 13 |
| 13 |
| 10 |
| 14 |
| 12 |
| 11 |
| 10 |
| 8 |
| 13 |
| 12 |
| 7 |
| 11 |
| 12 |
| 12 |
| 4 |
| 12 |
| 11 |
| 12 |
| 13 |
| 6 |
| 12 |
| 11 |
| 10 |
| 9 |
| 9 |
| 14 |
| 5 |
| 8 |
| 15 |
| 12 |
| 4 |
| 12 |
| 14 |
| 15 |
| 5 |
| 11 |
| 10 |

| |
|---|
| 13 |
| 13 |
| 9 |
| 13 |
| 12 |
| 12 |
| 10 |
| 14 |
| 10 |
| 12 |
| 11 |
| 4 |
| 12 |
| 12 |
| 9 |
| 12 |
| 7 |
| 8 |
| 13 |
| 15 |
| 9 |
| 14 |
| 14 |
| 8 |
| 5 |
| 5 |
| 5 |
| 14 |
| 14 |
| 14 |
| 14 |
| 12 |
| 10 |
| 11 |
| 13 |
| 6 |
| 15 |
| 12 |
| 13 |
| 13 |
| 15 |
| 15 |
| 15 |
| 7 |
| 11 |

| |
|---|
| 9 |
| 6 |
| 4 |
| 4 |
| 13 |
| 7 |
| 14 |
| 8 |
| 11 |
| 11 |
| 11 |
| 13 |
| 14 |
| 14 |
| 14 |
| 11 |
| 14 |
| 13 |
| 10 |
| 15 |
| 13 |
| 14 |
| 9 |
| 10 |
| 11 |
| 11 |
| 12 |
| 14 |
| 12 |
| 15 |
| 8 |
| 5 |
| 8 |
| 8 |
| 14 |
| 14 |
| 11 |
| 15 |
| 11 |
| 4 |
| 12 |
| 13 |
| 11 |
| 11 |
| 13 |

| |
|---|
| 14 |
| 14 |
| 13 |
| 12 |
| 11 |
| 11 |
| 10 |
| 12 |
| 14 |
| 9 |
| 12 |
| 10 |
| 4 |
| 6 |
| 13 |
| 13 |
| 14 |
| 8 |
| 15 |
| 10 |
| 9 |
| 14 |
| 6 |
| 8 |
| 15 |
| 10 |
| 14 |
| 12 |
| 8 |
| 9 |
| 5 |
| 9 |
| 10 |
| 12 |
| 11 |
| 13 |
| 4 |
| 12 |
| 13 |
| 4 |
| 4 |
| 13 |
| 9 |
| 5 |
| 14 |

| |
|---|
| 14 |
| 13 |
| 4 |
| 11 |
| 14 |
| 4 |
| 13 |
| 10 |
| 12 |
| 6 |
| 9 |
| 12 |
| 13 |
| 14 |
| 7 |
| 10 |
| 14 |
| 9 |
| 15 |
| 13 |
| 4 |
| 13 |
| 12 |
| 10 |
| 15 |
| 12 |
| 14 |
| 10 |
| 14 |
| 11 |
| 10 |
| 11 |
| 14 |
| 15 |
| 13 |
| 4 |
| 7 |
| 14 |
| 12 |
| 12 |
| 11 |
| 7 |
| 11 |
| 12 |
| 4 |

| |
|---|
| 4 |
| 4 |
| 12 |
| 13 |
| 11 |
| 11 |
| 7 |
| 13 |
| 15 |
| 14 |
| 8 |
| 7 |
| 14 |
| 11 |
| 9 |
| 8 |
| 15 |
| 15 |
| 8 |
| 15 |
| 13 |
| 11 |
| 7 |
| 12 |
| 9 |
| 4 |
| 12 |
| 9 |
| 11 |
| 8 |
| 10 |
| 4 |
| 10 |
| 14 |
| 15 |
| 5 |
| 4 |
| 12 |
| 8 |
| 15 |
| 9 |
| 5 |
| 7 |
| 13 |
| 10 |

| |
|---|
| 13 |
| 12 |
| 14 |
| 7 |
| 11 |
| 10 |
| 12 |
| 12 |
| 10 |
| 8 |
| 14 |
| 11 |
| 13 |
| 11 |
| 12 |
| 10 |
| 10 |
| 9 |
| 9 |
| 13 |
| 12 |
| 11 |
| 12 |
| 5 |
| 5 |
| 11 |
| 14 |
| 14 |
| 14 |
| 13 |
| 11 |
| 13 |
| 9 |
| 10 |
| 13 |
| 14 |
| 5 |
| 14 |
| 13 |
| 12 |
| 12 |
| 11 |
| 13 |
| 4 |
| 11 |

| |
|---|
| 12 |
| 5 |
| 11 |
| 4 |
| 10 |
| 8 |
| 7 |
| 9 |
| 8 |
| 8 |
| 4 |
| 12 |
| 13 |
| 11 |
| 8 |
| 5 |
| 7 |
| 12 |
| 12 |
| 11 |
| 13 |
| 15 |
| 7 |
| 15 |
| 4 |
| 9 |
| 12 |
| 7 |
| 11 |
| 7 |
| 11 |
| 14 |
| 13 |
| 8 |
| 14 |
| 11 |
| 6 |
| 13 |
| 10 |
| 15 |
| 8 |
| 8 |
| 5 |
| 7 |
| 15 |

| |
|---|
| 15 |
| 11 |
| 12 |
| 8 |
| 6 |
| 6 |
| 6 |
| 10 |
| 14 |
| 12 |
| 11 |
| 13 |
| 10 |
| 14 |
| 5 |
| 14 |
| 14 |
| 11 |
| 13 |
| 12 |
| 14 |
| 13 |
| 11 |
| 14 |
| 14 |
| 11 |
| 14 |
| 15 |
| 8 |
| 12 |
| 10 |
| 12 |
| 14 |
| 7 |
| 4 |
| 5 |
| 11 |
| 4 |
| 15 |
| 14 |
| 5 |
| 13 |
| 4 |
| 12 |
| 14 |

| |
|---|
| 14 |
| 13 |
| 11 |
| 11 |
| 15 |
| 14 |
| 15 |
| 15 |
| 4 |
| 6 |
| 7 |
| 6 |
| 10 |
| 8 |
| 13 |
| 9 |
| 11 |
| 13 |
| 4 |
| 12 |
| 10 |
| 8 |
| 7 |
| 7 |
| 5 |
| 15 |
| 11 |
| 9 |
| 7 |
| 14 |
| 5 |
| 10 |
| 9 |
| 13 |
| 15 |
| 11 |
| 13 |
| 10 |
| 7 |
| 14 |
| 11 |
| 10 |
| 11 |
| 12 |
| 13 |

| |
|---|
| 11 |
| 12 |
| 10 |
| 8 |
| 9 |
| 13 |
| 8 |
| 12 |
| 4 |
| 6 |
| 13 |
| 14 |
| 14 |
| 11 |
| 15 |
| 10 |
| 4 |
| 10 |
| 14 |
| 14 |
| 4 |
| 11 |
| 14 |
| 5 |
| 13 |
| 10 |
| 14 |
| 13 |
| 14 |
| 8 |
| 14 |
| 11 |
| 6 |
| 12 |
| 4 |
| 6 |
| 7 |
| 5 |
| 15 |
| 13 |
| 4 |
| 12 |
| 14 |
| 4 |
| 8 |

| |
|---|
| 14 |
| 15 |
| 8 |
| 7 |
| 12 |
| 15 |
| 8 |
| 11 |
| 6 |
| 11 |
| 10 |
| 11 |
| 13 |
| 14 |
| 12 |
| 13 |
| 8 |
| 8 |
| 6 |
| 13 |
| 12 |
| 10 |
| 10 |
| 6 |
| 5 |
| 8 |
| 12 |
| 15 |
| 11 |
| 14 |
| 8 |
| 8 |
| 8 |
| 4 |
| 8 |
| 13 |
| 15 |
| 6 |
| 15 |
| 15 |
| 14 |
| 15 |
| 4 |
| 11 |
| 6 |

| |
|---|
| 11 |
| 4 |
| 11 |
| 13 |
| 13 |
| 12 |
| 6 |
| 11 |
| 12 |
| 13 |
| 14 |
| 10 |
| 6 |
| 14 |
| 11 |
| 14 |
| 15 |
| 13 |
| 12 |
| 15 |
| 15 |
| 11 |
| 14 |
| 4 |
| 6 |
| 8 |
| 13 |
| 15 |
| 15 |
| 15 |
| 5 |
| 11 |
| 13 |
| 7 |
| 14 |
| 14 |
| 12 |
| 4 |
| 11 |
| 5 |
| 7 |
| 13 |
| 13 |
| 11 |
| 15 |

| |
|---|
| 11 |
| 13 |
| 12 |
| 14 |
| 13 |
| 12 |
| 9 |
| 14 |
| 6 |
| 7 |
| 13 |
| 13 |
| 11 |
| 11 |
| 14 |
| 15 |
| 12 |
| 7 |
| 15 |
| 15 |
| 12 |
| 14 |
| 13 |
| 13 |
| 11 |
| 10 |
| 10 |
| 9 |
| 14 |
| 5 |
| 13 |
| 9 |
| 5 |
| 11 |
| 14 |
| 14 |
| 10 |
| 14 |
| 11 |
| 15 |
| 15 |
| 11 |
| 12 |
| 13 |
| 15 |

| |
|---|
| 14 |
| 14 |
| 12 |
| 5 |
| 14 |
| 14 |
| 10 |
| 8 |
| 10 |
| 11 |
| 9 |
| 9 |
| 11 |
| 5 |
| 15 |
| 10 |
| 11 |
| 14 |
| 5 |
| 12 |
| 12 |
| 5 |
| 5 |
| 13 |
| 11 |
| 4 |
| 10 |
| 11 |
| 8 |
| 7 |
| 13 |
| 14 |
| 12 |
| 14 |
| 15 |
| 14 |
| 12 |
| 8 |
| 12 |
| 4 |
| 13 |
| 13 |
| 5 |
| 15 |
| 15 |

| |
|---|
| 13 |
| 7 |
| 7 |
| 5 |
| 6 |
| 6 |
| 5 |
| 6 |
| 4 |
| 13 |
| 14 |
| 13 |
| 6 |
| 4 |
| 5 |
| 5 |
| 15 |
| 14 |
| 4 |
| 8 |
| 13 |
| 15 |
| 8 |
| 14 |
| 6 |
| 4 |
| 12 |
| 12 |
| 5 |
| 15 |
| 8 |
| 12 |
| 12 |
| 14 |
| 11 |
| 9 |
| 12 |
| 9 |
| 13 |
| 11 |
| 11 |
| 4 |
| 6 |
| 6 |
| 5 |

| |
|---|
| 9 |
| 7 |
| 5 |
| 14 |
| 11 |
| 11 |
| 6 |
| 11 |
| 12 |
| 15 |
| 15 |
| 13 |
| 8 |
| 10 |
| 14 |
| 6 |
| 7 |
| 14 |
| 13 |
| 14 |
| 13 |
| 4 |
| 12 |
| 8 |
| 13 |
| 15 |
| 15 |
| 13 |
| 14 |
| 8 |
| 8 |
| 7 |
| 14 |
| 5 |
| 9 |
| 9 |
| 11 |
| 11 |
| 13 |
| 12 |
| 11 |
| 11 |
| 5 |
| 13 |
| 13 |

| |
|---|
| 12 |
| 13 |
| 13 |
| 14 |
| 14 |
| 6 |
| 12 |
| 6 |
| 14 |
| 8 |
| 9 |
| 12 |
| 13 |
| 12 |
| 8 |
| 4 |
| 10 |
| 4 |
| 15 |
| 13 |
| 11 |
| 14 |
| 10 |
| 14 |
| 12 |
| 12 |
| 11 |
| 14 |
| 8 |
| 11 |
| 7 |
| 13 |
| 9 |
| 11 |
| 13 |
| 12 |
| 14 |
| 10 |
| 14 |
| 14 |
| 13 |
| 12 |
| 13 |
| 11 |
| 15 |

| |
|---|
| 13 |
| 14 |
| 13 |
| 14 |
| 12 |
| 9 |
| 10 |
| 12 |
| 11 |
| 12 |
| 7 |
| 10 |
| 13 |
| 13 |
| 6 |
| 6 |
| 5 |
| 13 |
| 8 |
| 8 |
| 11 |
| 9 |
| 8 |
| 12 |
| 5 |
| 14 |
| 9 |
| 13 |
| 12 |
| 13 |
| 5 |
| 13 |
| 6 |
| 14 |
| 15 |
| 14 |
| 6 |
| 12 |
| 15 |
| 8 |
| 13 |
| 15 |
| 14 |
| 15 |
| 10 |

| |
|---|
| 14 |
| 12 |
| 4 |
| 14 |
| 14 |
| 11 |
| 14 |
| 13 |
| 14 |
| 14 |
| 12 |
| 13 |
| 14 |
| 15 |
| 14 |
| 14 |
| 13 |
| 15 |
| 7 |
| 13 |
| 14 |
| 13 |
| 4 |
| 14 |
| 6 |
| 14 |
| 14 |
| 15 |
| 9 |
| 9 |
| 10 |
| 9 |
| 5 |
| 6 |
| 10 |
| 12 |
| 5 |
| 12 |
| 11 |
| 13 |
| 13 |
| 13 |
| 4 |
| 13 |
