## Supplementary material for "*Drosophila* functional screening of *de novo* variants in autism uncovers deleterious variants and facilitates discovery of rare neurodevelopmental diseases": Table S2

**Supplemental Table 2: TG4 mutant and UAS (Reference and Variant) transgenic generation**

| Human Gene | HGNCID | FlyBaseID | Fly Gene | DIOPT Score | MiMIC | TG4 generated | UAS-ref generated | *H. sap.* Transcript | HA tagged (Y/N) | # SSC variants | UAS- var generated (this study) | Lethal? |
|---|---|---|---|---|---|---|---|---|---|---|---|---|
| *ABCC4* | 55 | FBgn0038740 | *CG4562* | 14 | MI12484 | Yes (Lee et al., 2018) | Yes - This study | NM_001105515 | N | 1 | 1 | Y |
| *ABCC5* | 56 | FBgn0039644 | *CG4562* | 14 | MI12484 | Yes (Lee et al., 2018) | Yes - This study | NM_005688 | N | 2 | 2 | Y |
| *ABL2* | 77 | FBgn0000017 | *Abl* | 10 | MI00347 | Yes (Lee et al., 2018) | Yes - This study | NM_001100108 | N | 1 | 1 | Y |
| *ACE* | 2707 | FBgn0012037 | *Ance* | 10 | MI05748 | Yes - This study | Yes - This study | NM_000789 | N | 1 | 1 | N |
| *ACHE* | 108 | FBgn0000024 | *Ace* | 9 | MI07345 | Yes - This study | Yes - This study | NM_015831 | N | 2 | 2 | Y |
| *AK1* | 361 | FBgn0022709 | *Adk1* | 13 | MI13455 | Yes (Lee et al., 2018) | Yes - This study | NM_000476 | N | 1 | 1 | N |
| *ALDH18A1* | 9722 | FBgn0037146 | *CG7470* | 15 | MI13272 | Yes (Lee et al., 2018) | Yes - This study | NM_002860.3 | Y | 1 | 1 | Y |
| *ALDH1L1* | 3978 | FBgn0032945 | *CG8665* | 14 | MI09533 | Yes (Lee et al., 2018) | Yes - This study | NM_001270364.1 | N | 1 | 1 | N |
| *ALDH3A1* | 405 | FBgn0010548 | *Aldh-III* | 9 | MI00204 | Yes - This study | Yes - This study | NM_014244 | Y | 1 | 1 | Y |
| *ARHGAP21* | 23725 | FBgn0031118 | *RhoGAP19D* | 7 | MI07163 | Yes (Lee et al., 2018) | NG | N/A | - | 1 | 0 | N |
| *ARID1B* | 18040 | FBgn0261885 | *osa* | 12 | MI02292 | Yes (Lee et al., 2018) | NG | N/A | - | 3 | 0 | Y |
| *ATP10A* | 13542 | FBgn0032120 | *CG33298* | 12 | MI08659 | Yes - This study | Yes - This study | NM_024490.3 | Y | 3 | 0 | N |
| *ATP2B2* | 815 | FBgn0259214 | *PMCA* | 12 | MI12515 | Yes (Lee et al., 2018) | Yes - This study | NM_001001331.3 | N | 1 | 1 | Y |
| *ATP2B4* | 817 | FBgn0259214 | *PMCA* | 13 | MI12515 | Yes (Lee et al., 2018) | Yes - This study | NM_001001396 | N | 1 | 0 | Y |
| *BAIAP2L1* | 21649 | FBgn0052082 | *IRSp53* | 8 | MI12856 | Yes - This study | Yes - This study | NM_018842 | N | 1 | 1 | N |
| *BCHE* | 983 | FBgn0000024 | *Ace* | 10 | MI07345 | Yes - This study | Yes - This study | NM_000055 | Y | 1 | 1 | Y |
| *BEST3* | 17105 | FBgn0040238 | *Best1* | 12 | MI05442 | Yes - This study | Yes - This study | NM_032735.2 | N | 1 | 1 | N |
| *BMP1* | 1067 | FBgn0004885 | *tok* | 11 | MI06118 | Yes - This study | Yes - This study | NM_001655 | Y | 1 | 1 | Y |
| *CACNA1C* | 1390 | FBgn0001991 | *Ca-alpha1D* | 12 | MI11167 | Failed conversion | NG | N/A | - | 1 | 0 | - |
| *CACNA1S* | 1397 | FBgn0001991 | *Ca-alpha1D* | 13 | MI11167 | Failed conversion | NG | N/A | - | 2 | 0 | - |
| *CAMK2A* | 1460 | FBgn0264607 | *CaMKII* | 9 | MI03976 | Failed conversion | Yes - This study | NM_015981 | N | 1 | 1 | - |
| *CARS* | 1493 | FBgn0027091 | *CysRS* | 15 | MI15366 | Failed conversion | Yes - This study | NM_001751.5 | Y | 1 | 1 | - |
| *CASK* | 1497 | FBgn0013759 | *CASK* | 11 | MI01748 | Yes - This study | Yes - This study | NM_001126054.2 | Y | 1 | 0 | N |
| *CAT* | 1516 | FBgn0000261 | *Cat* | 14 | MI04522 | Yes (Lee et al., 2018) | Yes - This study | NM_001003696 | Y | 1 | 1 | Y |
| *CEP135* | 29086 | FBgn0036480 | *Cep135* | 12 | MI02509 | Yes (Lee et al., 2018) | Yes - This study | NM_025009.4 | Y | 1 | 1 | N |
| *CHST2* | 1970 | FBgn0051637 | *CG31637* | 8 | MI03598 | Yes (Lee et al., 2018) | Yes - This study | NM_004267.4 | Y | 1 | 1 | Y |
| *CLCNKB* | 2027 | FBgn0051116 | *ClC-a* | 5 | MI05423 | Yes (Lee et al., 2018) | Yes - This study | NM_000085.4 | Y | 1 | 1 | Y |
| *CLIP2* | 2586 | FBgn0020503 | *CLIP-190* | 11 | MI02784 | Yes (Lee et al., 2018) | Yes - This study | NM_003388.4 | Y | 1 | 1 | N |
| *CRHR1* | 2357 | FBgn0033744 | *Dh44-R2* | 9 | MI09411 | Yes (Lee et al., 2018) | NG | N/A | - | 1 | 0 | N |
| *CSAD* | 18966 | FBgn0000153 | *b* | 13 | MI10547 | Yes (Lee et al., 2018) | Yes - This study | NM_015989 | Y | 1 | 1 | N |
| *CTNNB1* | 2514 | FBgn0000117 | *arm* | 14 | MI08675 | Yes (Lee et al., 2018) | Yes - This study | NM_001098209 | N | 2 | 1 | Y |
| *DDR2* | 2731 | FBgn0053531 | *Ddr* | 13 | MI06226 | Yes - This study | Yes - This study | NM_001014796 | N | 1 | 0 | N |
| *DLC1* | 2897 | FBgn0285955 | *cv-c* | 9 | MI03552 | Yes - This study | Yes - This study | NM_182643 | N | 1 | 0 | N |
| *DNAH7* | 18661 | FBgn0013810 | *Dhc36C* | 13 | MI05945 | Failed conversion | NG | N/A | - | 1 | 0 | - |
| *DOCK1* | 2987 | FBgn0015513 | *mbc* | 13 | MI06051 | Yes (Lee et al., 2018) | NG | N/A | - | 1 | 0 | Y |
| *DOCK4* | 19192 | FBgn0264324 | *spg* | 10 | MI09702 | Yes (Lee et al., 2018) | NG | N/A | - | 1 | 0 | N |
| *DPP6* | 3010 | FBgn0263780 | *CG17684* | 9 | MI13487 | Yes (Lee et al., 2018) | Yes - This study | NM_001039350.2 | Y | 1 | 0 | N |
| *DPYSL2* | 3014 | FBgn0023023 | *CRMP* | 10 | MI06892 | Yes - This study | Yes - This study | NM_001386.5 | Y | 1 | 1 | N |
| *DPYSL3* | 3015 | FBgn0023023 | *CRMP* | 11 | MI06892 | Yes - This study | Yes - This study | NM_001197294.1 | Y | 1 | 1 | N |
| *DST* | 1090 | FBgn0013733 | *shot* | 11 | MI14321 | Yes (Lee et al., 2018) | NG | N/A | - | 2 | 0 | N |
| *ELAVL3* | 3314 | FBgn0086675 | *fne* | 10 | MI09399 | Yes (Lee et al., 2018) | Yes - This study | NM_001420 | N | 2 | 1 | N |
| *EP400* | 11958 | FBgn0020306 | *dom* | 10 | MI08014 | Yes (Lee et al., 2018) | Yes - This study | NM_015409.4 | N | 1 | 0 | Y |
| *EPHA1* | 3385 | FBgn0025936 | *Eph* | 6 | MI05205 | Yes (Lee et al., 2018) | Yes - This study | NM_020384 | Y | 1 | 1 | N |
| *EPHB1* | 3392 | FBgn0025936 | *Eph* | 13 | MI05205 | Yes (Lee et al., 2018) | Yes - This study | NM_004441 | N | 1 | 1 | N |
| *EPT1* | 29361 | FBgn0053116 | *CG33116* | 14 | MI01935 | Failed conversion | Yes - This study | NM_033505 | N | 1 | 1 | - |
| *EXD2* | 20217 | FBgn0037901 | *Exd2* | 14 | MI03063 | Failed conversion | Yes - This study | NM_001193363.1 | Y | 1 | 1 | - |
| *FGGY* | 25610 | FBgn0035484 | *CG11594* | 14 | MI04075 | Yes - This study | Yes - This study | NM_001278224.1 | N | 1 | 0 | N |
| *FRYL* | 29127 | FBgn0016081 | *fry* | 14 | MI12326 | Yes (Lee et al., 2018) | NG | N/A | - | 1 | 0 | Y |
| *GCLC* | 4311 | FBgn0040319 | *Gclc* | 14 | MI04005 | Yes (Lee et al., 2018) | Yes - This study | NM_001498.4 | Y | 1 | 1 | N |
| *GLRA2* | 4327 | FBgn0024963 | *GluClalpha* | 10 | MI14426 | Yes - This study | Yes - This study | NM_002063 | Y | 2 | 1 | N |
| *GNAO1* | 4389 | FBgn0001122 | *Galphao* | 14 | MI00833 | Yes - This study | Yes - This study | NM_138736.2 | Y | 1 | 0 | N |
| *GPC5* | 4453 | FBgn0263930 | *dally* | 14 | MI01424 | Yes (Lee et al., 2018) | Yes - This study | NM_004466.5 | Y | 1 | 1 | N |
| *GRIA1* | 4571 | FBgn0264000 | *GluRIB* | 11 | MI00476 | Yes (Lee et al., 2018) | Yes - This study | NM_000827.3 | Y | 1 | 1 | N |
| *GRIK5* | 4583 | FBgn0039916 | *Ekar* | 7 | MI02500 | Failed conversion | Yes - This study | NM_002088.4 | N | 1 | 0 | - |
| *GRIN2B* | 4586 | FBgn0053513 | *Nmdar2* | 11 | MI09281 | Yes (Lee et al., 2018) | NG | N/A | - | 2 | 0 | N |

| | | | | | | | | | | | | |
|---|---|---|---|---|---|---|---|---|---|---|---|---|
| *GRK4* | 4543 | FBgn0261988 | *Gprk2* | 12 | MI01907 | Yes (Lee et al., 2018) | Yes - This study | NM_001004056 | Y | 1 | 1 | Y |
| *HTR1D* | 5289 | FBgn0263116 | *5-HT1B* | 4 | MI05213 | Yes - This study | Yes - This study | NM_000864 | Y | 1 | 1 | N |
| *IGF2R* | 5467 | FBgn0051072 | *Lerp* | 8 | MI09611 | Yes (Lee et al., 2018) | Yes - This study | NM_000876.2 | Y | 1 | 0 | N |
| *IRF2BPL* | 14282 | FBgn0030400 | *Pits* | 11 | MI02926 | Yes (Lee et al., 2018) | Yes - (Marcogliese et al., 2018) | NM_024496.3 | N | 2 | 2 | Y |
| *ITGA2B* | 6138 | FBgn0001250 | *if* | 10 | MI12214 | Yes (Lee et al., 2018) | Failed injection | N/A | - | 1 | 0 | Y |
| *ITGA8* | 6144 | FBgn0001250 | *if* | 12 | MI12214 | Yes (Lee et al., 2018) | Yes - This study | NM_003638.1 | Y | 1 | 1 | Y |
| *ITPR3* | 6182 | FBgn0010051 | *Itp-r83A* | 11 | MI15600 | Yes - This study | NG | N/A | N | 2 | 0 | N |
| *JUP* | 6207 | FBgn0000117 | *arm* | 10 | MI08675 | Yes (Lee et al., 2018) | Yes - This study | NM_002230.2 | Y | 1 | 1 | Y |
| *KCND3* | 6239 | FBgn0005564 | *Shal* | 12 | MI00446 | Yes - This study | Yes - This study | NM_172198.2 | Y | 2 | 1 | N |
| *KCNH3* | 6252 | FBgn0011589 | *Elk* | 11 | MI02485 | Failed conversion | NG | N/A | - | 1 | 0 | - |
| *KCNH8* | 18864 | FBgn0011589 | *Elk* | 12 | MI02485 | Failed conversion | Yes - This study | NM_144633.2 | Y | 1 | 0 | - |
| *KDM2A* | 13606 | FBgn0037659 | *Kdm2* | 13 | MI08481 | Yes - This study | Yes - This study | NM_012308.2 | N | 1 | 1 | N |
| *KDM2B* | 13610 | FBgn0037659 | *Kdm2* | 11 | MI08481 | Yes - This study | NG | N/A | - | 1 | 0 | N |
| *KDR* | 6307 | FBgn0032006 | *Pvr* | 10 | MI04181 | Yes - This study | Yes - This study | NM_002253 | N | 1 | 1 | Y |
| *KMT2C* | 13726 | FBgn0263667 | *Lpt* | 7 | MI09585 | Yes (Lee et al., 2018) | NG | N/A | - | 3 | 0 | Y |
| *LAMA1* | 6481 | FBgn0261563 | *wb* | 9 | MI07688 | Yes (Lee et al., 2018) | NG | N/A | - | 1 | 0 | Y |
| *LAMA2* | 6482 | FBgn0261563 | *wb* | 8 | MI07688 | Yes (Lee et al., 2018) | Yes - This study | NM_000426.3 | N | 1 | 0 | Y |
| *LRCH4* | 6691 | FBgn0032633 | *Lrch* | 7 | MI05027 | Yes (Lee et al., 2018) | Yes - This study | NM_002319.4 | N | 1 | 1 | Y |
| *LRP1* | 6692 | FBgn0053087 | *LRP1* | 11 | MI03128 | Yes (Lee et al., 2018) | NG | N/A | - | 2 | 0 | Y |
| *MACF1* | 13664 | FBgn0013733 | *shot* | 10 | MI14321 | Yes (Lee et al., 2018) | NG | N/A | - | 1 | 0 | N |
| *MADD* | 6766 | FBgn0030613 | *Rab3-GEF* | 15 | MI01069 | Yes (Lee et al., 2018) | Yes - This study | NM_130471 | N | 1 | 1 | N |
| *MANBA* | 6831 | FBgn0037215 | *beta-Man* | 15 | MI03739 | Yes - This study | Yes - This study | NM_005908 | N | 1 | 0 | N |
| *MAP4K1* | 6863 | FBgn0263395 | *hppy* | 6 | MI03637 | Yes (Lee et al., 2018) | Yes - This study | NM_007181.5 | Y | 1 | 1 | N |
| *MBNL1* | 6923 | FBgn0265487 | *mbl* | 10 | MI00139 | Yes (Lee et al., 2018) | Yes - This study | NM_021038 | Y | 1 | 1 | Y |
| *MEGF11* | 29635 | FBgn0027594 | *drpr* | 11 | MI07659 | Yes - This study | Yes - This study | NM_032445 | N | 2 | 1 | N |
| *MINK1* | 17565 | FBgn0010909 | *msn* | 12 | MI09440 | Yes (Lee et al., 2018) | Yes - This study | NM_153827.4 | Y | 1 | 1 | Y |
| *MYH3* | 7573 | FBgn0264695 | *Mhc* | 9 | MI03941 | Yes (Lee et al., 2018) | Yes - This study | NM_002470.3 | N | 1 | 0 | N |
| *MYH9* | 7579 | FBgn0265434 | *Mhc* | 9 | MI03941 | Yes (Lee et al., 2018) | Yes - This study | NM_002473.4 | Y | 1 | 1 | N |
| *MYO7B* | 7607 | FBgn0000317 | *ck* | 8 | MI10140 | Yes (Lee et al., 2018) | NG | N/A | - | 3 | 0 | Y |
| *NCOR1* | 7672 | FBgn0265523 | *Smr* | 10 | MI12579 | Yes - This study | Yes - This study | NM_006311.3 | N | 1 | 1 | Y |
| *NID2* | 13389 | FBgn0026403 | *Ndg* | 11 | MI15397 | Yes - This study | Yes - This study | NM_007361 | N | 1 | 0 | N |
| *NLGN1* | 14291 | FBgn0083963 | *Nlg3* | 11 | MI00445 | Yes (Lee et al., 2018) | Yes - This study | NM_014932.2 | N | 1 | 1 | N |
| *NLGN3* | 14289 | FBgn0083963 | *Nlg3* | 12 | MI00445 | Yes (Lee et al., 2018) | NG | N/A | - | 1 | 1 | N |
| *NOS3* | 7876 | FBgn0011676 | *Nos* | 11 | MI15126 | Yes - This study | Yes - This study | NM_000603.4 | Y | 1 | 0 | N |
| *NPFFR2* | 4525 | FBgn0038880 | *SIFaR* | 8 | MI05376 | Yes - This study | Yes - This study | NM_053036.2 | N | 1 | 1 | Y |
| *NR2F1* | 7975 | FBgn0003651 | *svp* | 9 | MI01102 | Failed conversion | Yes - This study | NM_005654 | N | 1 | 1 | - |
| *NTN1* | 8029 | FBgn0015773 | *NetA* | 12 | MI11608 | Yes - This study | Yes - This study | NM_004822 | N | 1 | 1 | N |
| *NTN5* | 25208 | FBgn0015774 | *NetB* | 4 | MI11608 | Yes - This study | Yes - This study | NM_145807 | N | 1 | 0 | N |
| *P4HA2* | 8547 | FBgn0039776 | *PH4alphaEFB* | 11 | MI06111 | Yes - This study | Yes - This study | NM_001017974.1 | Y | 1 | 1 | Y |
| *PC* | 8636 | FBgn0027580 | *PCB* | 14 | MI02451 | Yes - This study | Yes - This study | NM_022172.2 | Y | 1 | 1 | N |
| *PCDH15* | 14674 | FBgn0039709 | *Cad99C* | 11 | MI08439 | Yes (Lee et al., 2018) | NG | N/A | - | 1 | 0 | N |
| *PDGFRB* | 8804 | FBgn0032006 | *Pvr* | 6 | MI04181 | Yes - This study | Yes - This study | NM_002609.3 | Y | 1 | 1 | Y |
| *PDK2* | 8810 | FBgn0017558 | *Pdk* | 13 | MI07697 | Yes - This study | Yes - This study | NM_002611 | N | 1 | 1 | Y |
| *PDZD2* | 18486 | FBgn0000008 | *a* | 4 | MI10698 | Yes (Lee et al., 2018) | NG | N/A | - | 1 | 0 | N |
| *PEAR1* | 33631 | FBgn0027594 | *drpr* | 8 | MI07659 | Yes - This study | Yes - This study | NM_001080471.1 | N | 2 | 1 | N |
| *PELI1* | 8827 | FBgn0025574 | *Pli* | 14 | MI00302 | Yes (Lee et al., 2018) | Yes - This study | NM_020651 | Y | 1 | 1 | N |
| *PIEZO2* | 26270 | FBgn0264953 | *Piezo* | 13 | MI04189 | Yes (Lee et al., 2018) | NG | N/A | - | 1 | 0 | N |
| *PITX1* | 9004 | FBgn0020912 | *Ptx1* | 8 | MI11305 | Yes (Lee et al., 2018) | Yes - This study | NM_002653 | Y | 1 | 1 | Y |
| *PLXDC1* | 20945 | FBgn0028331 | *l(1)G0289* | 11 | MI06498 | Yes (Lee et al., 2018) | Yes - This study | NM_020405 | N | 1 | 1 | Y |
| *PNISR* | 21222 | FBgn0051211 | *CG31211* | 8 | MI09858 | Yes (Lee et al., 2018) | NG | N/A | - | 1 | 0 | N |
| *PPP1R9A* | 14946 | FBgn0010905 | *Spn* | 11 | MI06873 | Yes (Lee et al., 2018) | Yes - This study | NM_017650 | N | 1 | 0 | N |
| *PRKD1* | 9407 | FBgn0038603 | *PKD* | 13 | MI09308 | Yes - This study | Yes - This study | NM_002742 | N | 1 | 1 | N |
| *PRPS1L1* | 9463 | FBgn0036030 | *CG6767* | 11 | MI09551 | Yes (Lee et al., 2018) | Yes - This study | NM_175886.2 | N | 1 | 1 | Y |
| *PSD4* | 19096 | FBgn0051158 | *Efa6* | 8 | MI00261 | Failed conversion | NG | N/A | - | 1 | 0 | - |
| *PTK7* | 9618 | FBgn0004839 | *otk* | 14 | MI14316 | Yes (Lee et al., 2018) | Failed injection | NM_152882 | N | 3 | 1 | N |
| *PTPRF* | 9670 | FBgn0000464 | *Lar* | 11 | MI03909 | Yes (Lee et al., 2018) | Yes - This study | BC048768 | N | 1 | 1 | Y |
| *PXDN* | 14966 | FBgn0011828 | *Pxn* | 13 | MI01492 | Yes (Lee et al., 2018) | Yes - This study | NM_012293 | N | 1 | 1 | Y |
| *PXDNL* | 26359 | FBgn0011828 | *Pxn* | 11 | MI01492 | Yes (Lee et al., 2018) | NG | N/A | - | 1 | 0 | Y |
| *RALGAPA1* | 17770 | FBgn0039466 | *CG5521* | 14 | MI06530 | Yes (Lee et al., 2018) | Yes - This study | NM_001346243.2 | N | 1 | 1 | Y |

| | | | | | | | | | | | | |
|---|---|---|---|---|---|---|---|---|---|---|---|---|
| *RIMS2* | 17283 | FBgn0053547 | *Rim* | 10 | MI07142 | Yes - This study | Yes - This study | NM_014677.4 | Y | 1 | 0 | N |
| *SCARB2* | 1665 | FBgn0010435 | *emp* | 10 | MI12296 | Yes (Lee et al., 2018) | Yes - This study | NM_005506.3 | Y | 1 | 1 | Y |
| *SDK2* | 19308 | FBgn0021764 | *sdk* | 11 | MI01498 | Yes (Lee et al., 2018) | Yes - This study | NM_001144952.1 | Y | 1 | 0 | N |
| *SEC14L5* | 29032 | FBgn0031814 | *retm* | 14 | MI12531 | Yes (Lee et al., 2018) | Yes - This study | NM_014692.1 | N | 1 | 0 | N |
| *SEMA5A* | 10736 | FBgn0284221 | *Sema5c* | 13 | MI10577 | Yes (Lee et al., 2018) | NG | N/A | - | 1 | 0 | N |
| *SH2D3C* | 16884 | FBgn0031762 | *CG9098* | 14 | MI14545 | Yes - This study | Yes - This study | NM_005489 | Y | 1 | 1 | N |
| *SLC23A1* | 10974 | FBgn0037807 | *CG6293* | 13 | MI14396 | Yes (Lee et al., 2018) | Yes - This study | NM_152685 | N | 1 | 1 | N |
| *SLC8A2* | 11069 | FBgn0013995 | *Calx* | 12 | MI09964 | Failed conversion | Yes - This study | NM_015063.2 | N | 1 | 1 | - |
| *SLCO4A1* | 10953 | FBgn0051634 | *Oatp26F* | 15 | MI00414 | Failed conversion | Yes - This study | NM_016354 | N | 1 | 1 | - |
| *SOGA3* | 21494 | FBgn0031869 | *CG18304* | 6 | MI00083 | Yes (Lee et al., 2018) | Yes - This study | NM_001012279.2 | Y | 1 | 1 | Y |
| *SRCAP* | 16974 | FBgn0020306 | *dom* | 8 | MI08014 | Yes (Lee et al., 2018) | Yes - This study | NM_006662.2 | N | 3 | 2 | Y |
| *TANC2* | 30212 | FBgn0041096 | *rols* | 11 | MI02479 | Yes (Lee et al., 2018) | Yes - This study | NM_025185.3 | N | 2 | 2 | Y |
| *TMEM201* | 33719 | FBgn0034447 | *CG7744* | 7 | MI04719 | Yes (Lee et al., 2018) | NG | N/A | - | 1 | 0 | Y |
| *TNIK* | 30765 | FBgn0010909 | *msn* | 13 | MI09440 | Yes (Lee et al., 2018) | NG | N/A | - | 1 | 0 | Y |
| *TNKS* | 11941 | FBgn0027508 | *Tnks* | 12 | MI05449 | Failed conversion | NG | N/A | - | 1 | 0 | - |
| *TRIP12* | 12306 | FBgn0260794 | *ctrip* | 10 | MI14762 | Yes - This study | Yes - This study | NM_001284214.1 | N | 3 | 1 | Y |
| *TRPM1* | 7146 | FBgn0265194 | *Trpm* | 12 | MI04585 | Yes (Lee et al., 2018) | Yes - This study | NM_001252024.1 | N | 1 | 1 | Y |
| *TRPM6* | 17995 | FBgn0265194 | *Trpm* | 11 | MI04585 | Yes (Lee et al., 2018) | Yes - This study | NM_017662.4 | Y | 2 | 2 | Y |
| *TRPM7* | 17994 | FBgn0265194 | *Trpm* | 12 | MI04585 | Yes (Lee et al., 2018) | Yes - This study | NM_017672.5 | Y | 1 | 1 | Y |
| *TRRAP* | 12347 | FBgn0053554 | *Nipped-A* | 13 | MI10513 | Yes (Lee et al., 2018) | Failed injection | N/A | - | 3 | 0 | Y |
| *TSC2* | 12363 | FBgn0005198 | *gig* | 13 | MI14761 | Yes (Lee et al., 2018) | Yes - This study | NM_000548 | Y | 2 | 2 | Y |
| *TULP4* | 15530 | FBgn0039530 | *Tusp* | 13 | MI04698 | Yes (Lee et al., 2018) | Yes - This study | NM_020245.4 | N | 1 | 0 | N |
| *UNC80* | 26582 | FBgn0039536 | *unc80* | 15 | MI07137 | Yes (Lee et al., 2018) | NG | N/A | - | 1 | 0 | Y |
| *USP30* | 20065 | FBgn0029819 | *Usp30* | 14 | MI09252 | Yes (Lee et al., 2018) | Yes - This study | NM_032663 | Y | 2 | 1 | N |
| *WDFY3* | 20751 | FBgn0043362 | *bchs* | 15 | MI14478 | Yes (Lee et al., 2018) | NG | N/A | - | 3 | 0 | N |
| *WDFY4* | 29323 | FBgn0043362 | *bchs* | 7 | MI14478 | Yes (Lee et al., 2018) | NG | N/A | - | 2 | 0 | N |
| *XPNPEP2* | 12823 | FBgn0038072 | *CG6225* | 6 | MI15524 | Yes (Lee et al., 2018) | NG | N/A | - | 1 | 0 | N |
| *YIPF5* | 24877 | FBgn0032465 | *Yip1d1* | 15 | MI10524 | Yes (Lee et al., 2018) | Yes - This study | NM_030799.9 | N | 1 | 0 | Y |
| *ZMYND8* | 9397 | FBgn0039863 | *CG1815* | 13 | MI01646 | Yes - This study | Yes - This study | NM_001281771.2 | N | 1 | 0 | N |
