## Supplementary material for "*Drosophila* functional screening of *de novo* variants in autism uncovers deleterious variants and facilitates discovery of rare neurodevelopmental diseases": Table S3

**Supplemental Table 3: UAS variant transgenic generation**

| Human Gene | HGNCID | *H. sap.* Transcript | SSC var (cDNA) | SSC var protein | Mutagenesis method | Forward primer | Reverse primer | PolyPhen2 | SIFT | CADD |
|---|---|---|---|---|---|---|---|---|---|---|
| *ABCC4* | 55 | NM_001105515 | c.826A>G | p.(M276V) | SDM | GATCAGGACCGTGAATGAA | CTGGCATCCGTGAAAGTTG | 0.654 | 0.024 | 17.16 |
| *ABCC5* | 56 | NM_005688 | c.3137C>T | p.(T1046M) | SDM | GACAATATCATGCAGTCAC | CAGACGCTTCAGCTCCCG | 0.987 | 0.001 | 20.8 |
| *ABCC5* | 56 | NM_005688 | c.2089C>T | p.(R697W) | SDM | CAGCCTTGCCTGGGCCTTC | ATCCTCTGGCGCTGCCCA | 1 | 0 | 22.4 |
| *ABL2* | 77 | NM_001100108 | c.3295G>A | p.(A1099T) | SDM | ACTGTCCAGTACACTCACC | AGGTCAGCACATTCCAGC | 0.998 | 0.013 | 20.9 |
| *ACE* | 2707 | NM_000789 | c.2453A>G | p.(Y818C) | HiTM | GCTGCCCGGCTCAATGGC | CGAGTCCCCTGCATCTACA | 0.954 | 0.003 | 7.512 |
| *ACHE* | 108 | NM_015831 | c.1643C>T | p.(P548L) | HiTM | TGTCCTCGTCTGGATCTATA | CCACTGTAGAAGCCACCC | 0.847 | 0.183 | 21 |
| *ACHE* | 108 | NM_015831 | c.451G>A | p.(G151R) | SDM | CTGGATCTATAGGGGTGGC | ACGAGGACAGGGGTGGGG | 1 | 0 | 27.7 |
| *AK1* | 361 | NM_000476 | c.173C>T | p.(S58L) | HiTM | GCCAGGGGCAAGAAGCTG | CCCCTTCTCCATGATTTCC | 0.261 | 0.08 | 17.16 |
| *ALDH18A1* | 9722 | NM_002860.3 | c.2107G>C | p.(D703H) | SDM | CGTCACAGAGCACGAAAAC | ATGACATCCGTGTGGGAG | 0.007 | 0.012 | 14.45 |
| *ALDH1L1* | 3978 | NM_001270364.1 | c.2698A>C | p.(N900H) | HiTM | TCTAGGAGAGGCGGCTCTC | TTGACCCGCAGGTACTCGT | 0.133 | 0.152 | 9.814 |
| *ALDH3A1* | 405 | NM_014244 | c.1206C>G | p.(F402L) | HiTM | CCTTGCACTCTCTGCCCTT | CGCTGTTCCCCACGCCCC | 0.995 | 0.066 | 14.36 |
| *ATP2B2* | 815 | NM_001001331.3 | c.2453C>T | p.(T818M) | SDM | GTGGCCGTGATGGGGGAC | CACCTGCCGCTGCTCAGTC | 0.997 | 0 | 29.7 |
| *BAIAP2L1* | 21649 | NM_018842 | c.1442C>T | p.(A481V) | SDM | GGCATCGTTAGGAGCCGC | AACGGGACTGtAAAGCCGC | 0.003 | 0.164 | 9.777 |
| *BCHE* | 983 | NM_000055 | c.1297T>G | p.(F433V) | HiTM | CATATGCCCTGCCTTGGAC | TCTGAGAACTTCTTGGTGA | 0.727 | 0.033 | 11.96 |
| *BEST3* | 17105 | NM_032735.2 | c.388C>A | p.(R130S) | HiTM | GCTTAGAAGGACGCTGATC | GAGGTGAGATTGACGTAGC | 1 | 0 | 23.2 |
| *BMP1* | 1067 | NM_001655 | c.2779G>A | p.(G927S) | SDM | GCTCTTCGACAGCTACGAC | TCCATGTAGTCATAGCCG | 0.999 | 0.022 | 32 |
| *CAMK2A* | 1460 | NM_015981 | c.548A>T | p.(E183V) | HiTM | CCTGGATATCTCTCCCCAC | CGGGTCCTTCCGCAGCAC | 0.999 | 0 | 32 |
| *CARS* | 1493 | NM_001751.5 | c.1043A>G | p.(N348S) | HiTM | GGTTACGGCTATGTCTCCA | ATCAAAGTAGACAGACCCA | 0.483 | 0.062 | 11.42 |
| *CAT* | 1516 | NM_001003696 | c.611G>A | p.(G204E) | HiTM | TTCTTGTTCAGTGATCGGG | GCGATGTCCATCTGGAATC | 1 | 0 | 23.1 |
| *CEP135* | 29086 | NM_025009.4 | c.2839T>C | p.(S947P) | HiTM | TGCTTATGAATCTCAGATCC | ATGGCTTTTGCCATTGATGC | 0.999 | 0.003 | 25.1 |
| *CHST2* | 1970 | NM_004267.4 | c.155G>C | p.(R52P) | HiTM | CTCGGAATGAAGGTGTTCC | CAACACCAGCGCCTTCCTA | 0.989 | 0 | 12.82 |
| *CLCNKB* | 2027 | NM_000085.4 | c.528G>C | p.(M176I) | HiTM | TGCACCTGTCTGTGATGAT | CACGGCCCAGGTAGGCAG | 0.001 | 0.564 | 5.89 |
| *CLIP2* | 2586 | NM_003388.4 | c.37G>T | p.(G13W) | HiTM | CCTGAAGCCCCCCGGCCC | GGGCTGGAGTGCTTCCCC | 0.958 | 0.015 | 17.36 |
| *CSAD* | 18966 | NM_015989 | c.1232C>T | p.(A411V) | HiTM | GAGCGGCGCATCGACCAG | GTACCGGGCAAGGACAAA | 0.796 | 0.756 | 11.5 |
| *CTNNB1* | 2514 | NM_001098209 | c.1652C>T | p.(T551M) | HiTM | CAGGATACCCAGCGCCGT | CTGTGTCCCACCCATGGAC | 0.1 | 0.069 | 15 |
| *DPYSL2* | 3014 | NM_001386.5 | c.1486C>T | p.(R496C) | HiTM | TGAGCTGAGAGGGGTTCCT | GGTCCGTCATACAGGCCAC | 0.992 | 0.004 | 23.3 |
| *DPYSL3* | 3015 | NM_001197294.1 | c.415G>A | p.(V139I) | HiTM | TATCAAGGGAGGCAGAATC | AAGGACTGATCATCATTGA | 0.099 | 0.15 | 15.81 |
| *ELAVL3* | 3314 | NM_001420 | c.557T>C | p.(L186P) | HiTM | GAAGAGGCTATCAAAGGAC | CAGCGGCTTCTGCCCATTC | 0.998 | 0 | 18.85 |
| *EPHA1* | 3385 | NM_020384 | c.1699G>A | p.(V567I) | HiTM | CTTGCTGCTTGGGATTCTC | GCTCTCCTGGACCGGAAA | 0 | 0.773 | 0.038 |
| *EPHB1* | 3392 | NM_004441 | c.2746G>A | p.(V916M) | HiTM | CTTCACGGCCTTTACCACC | GCGCTGAGCCAGTCATCC | 0.999 | 0.011 | 32 |
| *EPT1* | 29361 | NM_033505 | c.244C>A | p.(H82N) | HiTM | TTATGCCTCAGCACCAGGT | CAGTCAGGCACGTGCTTGT | 0.122 | 0.532 | 13.85 |
| *EXD2* | 20217 | NM_001193363.1 | c.1539G>C | p.(E513D) | HiTM | GGGCCCTGCTCAACGCGG | TTCGATGAGTAGGCAGGCT | 0.002 | 0.744 | 7.915 |
| *GCLC* | 4311 | NM_001498.4 | c.382C>T | p.(R128W) | HiTM | GGCCAACATGCGAAAACG | ATAGAAGTAGCCTCCTTCC | 0.999 | 0 | 23.1 |
| *GLRA2* | 4327 | NM_002063 | c.407A>G | p.(N136S) | HiTM | CCAGATTTGTTCTTTGCCAC | GAAGTTGGCACCCTTCTCA | 0.999 | 0 | 25.1 |
| *GPC5* | 4453 | NM_004466.5 | c.398T>C | p.(M133T) | HiTM | TGCAGTACCTACAGGAACA | AGCAGCAGCCTCCAAGGC | 0.978 | 0.001 | 18.43 |
| *GRIA1* | 4571 | NM_000827.3 | c.653G>A | p.(R218H) | HiTM | GTGGACTGTGAATCAGAAC | GCCCAAGATAGCATTGAGC | 0.997 | 0.021 | 29.8 |
| *GRK4* | 4543 | NM_182982.3 | c.1153C>G | p.(P385A) | HiTM | AATGATTCAGGGACATTCT | TCTTTGTATTTTTTGAATGC | 0.999 | 0 | 24.9 |
| *HTR1D* | 5289 | NM_000864 | c.296C>A | p.(T99N) | HiTM | CCCATCAGCATCGCCTATA | GTTCCAGGTGTGGGTGATG | 0.99 | 0.076 | 14.42 |
| *IRF2BPL* | 14282 | NM_024496.3 | c.2102delA | N701X | SDM | CATTCCGGATTCCCCCATC | TTTGGGGGTGCACTTGGT | - | - | - |
| *IRF2BPL* | 14282 | NM_024496.3 | c.90C>G | p.(F30L) | SDM | TCTGGGACTTGTCGGAACC | TCATGGCCCAGGGCATGC | 0.989 | 0.002 | 14.94 |
| *ITGA8* | 6144 | NM_003638.1 | c.2242C>T | p.(R748C) | SDM | TGCAGTTCCATGTCTTGAG | AATCGGAGGCCCAGGGAA | 0.956 | 0.006 | 24.2 |
| *JUP* | 6207 | NM_002230.2 | c.2069A>G | p.(N690S) | SDM | ATTCCCATCAGTGAGCCCT | CATGCTCTGGGCAGCCTC | 0.039 | 0.553 | 2.456 |
| *KCND3* | 6239 | NM_172198.2 | c.257G>C | p.(R86P) | HiTM | AAGGAGTACTTCTTCGACC | GCGGAACACCTCGGGGTC | 1 | 0.001 | 22.9 |
| *KDM2A* | 13606 | NM_012308.2 | c.1346G>A | p.(R449K) | HiTM | TGTGCTCCCCGAAAGGACA | ATGGGTCAGATGCACTTGC | 0 | 1 | 5.746 |
| *KDR* | 6307 | NM_002253 | c.3511G>A | p.(D1171N) | SDM | TGCTCAGCAGAATGGCAAA | TTAGCTTGCAAGAGATTTCC | 0.697 | 0.004 | 25.9 |
| *LRCH4* | 6691 | NM_002319.4 | c.124G>A | p.(V42M) | HiTM | GCGGGCCCTAGAGGAGGC | TTCAGGGTCCCGGTGGCC | 0.891 | 0.002 | 22.5 |
| *MADD* | 6766 | NM_130471 | c.1540C>T | p.(R514C) | HiTM | GATGCACACGCGTACCCTC | ACAGGCCGAGGAAAGAGG | 1 | 0 | 29.8 |
| *MAP4K1* | 6863 | NM_007181.5 | c.2174T>C | p.(M725T) | HiTM | GATATGGTGATGGTGTTGA | CAGCTTCACAGAGCCATCC | 0.111 | 0.059 | 13.81 |
| *MBNL1* | 6923 | NM_021038 | c.134T>C | p.(V45A) | HiTM | CCTTCGAAAAGCTGCCAAC | GATTACTCGTCCATTTTCAC | 0.843 | 0.019 | 15.52 |
| *MEGF11* | 29635 | NM_032445 | c.2731C>T | p.(R911C) | HiTM | TGCTTGTGGAATGGATAGA | ATAATGTATGTGTTCTGACA | 0.365 | 0.062 | 16.38 |
| *MINK1* | 17565 | NM_153827.4 | c.805T>C | p.(C269R) | SDM | CATTGACACACGTCTCATC | AAGTCAATGAACTTCTTAG | 0.996 | 0 | 20.9 |
| *MYH9* | 7579 | NM_002473.4 | c.4712G>A | p.(R1571Q) | HiTM | ATGAAGGCCCAGTTCGAGC | GTCCCGGCCCTGCAGGTC | 0.973 | 0.106 | 36 |
| *NCOR1* | 7672 | NM_006311.3 | c.1705C>T | p.(P569S) | SDM | GCAAGCCACATCCCGGGG | TCTCTTTCCTCAGTTTCTTC | 0.998 | 0.575 | 13.24 |
| *NLGN1* | 14291 | NM_014932.2 | c.2383C>T | p.(H795Y) | HiTM | ACCAGGGATTCAGCCCTTA | GTAAATGTATTGAATGTGTA | 0.075 | 0.001 | 13.59 |
| *NLGN3* | 14289 | NM_018977.4 | c.583C>T | p.(R195W) | HiTM | GGATGAAGATGAAGACATC | GGTTTAGCACCACTGTCCC | 0.982 | 0.003 | 18.28 |

| | | | | | | | | | | |
|---|---|---|---|---|---|---|---|---|---|---|
| *NPFFR2* | 4525 | NM_053036.2 | c.489G>A | p.(M163I) | HiTM | TCTTCTTTTTGTGCATGATA | AGCAAACCACAGTATTTCC | 0.002 | 0.562 | 8.023 |
| *NR2F1* | 7975 | NM_005654 | c.1211G>A | p.(R404H) | HiTM | CCCATCGAAACTCTCATCC | CCCAGACAGTAACATATCC | 0.987 | 0.033 | 28.5 |
| *NTN1* | 8029 | NM_004822 | c.1346C>A | p.(A449D) | HiTM | CAGAGCCGCTCTCCCATC | AGGGATCTTTATGCAGGGG | 0.961 | 0.121 | 32 |
| *P4HA2* | 8547 | NM_001017974.1 | c.458G>A | p.(G153E) | SDM | ATTTCCAGAGAGGAACTTC | TGTGCCTGGGTCCAGCCT | 1 | 0 | 29.2 |
| *PC* | 8636 | NM_022172.2 | c.3125C>G | p.(P1042R) | HiTM | CGCCTCTTCCTGCAGGGA | AAACTCCTCTGCGATCTTG | 0.995 | 0.047 | 16.69 |
| *PDGFRB* | 8804 | NM_002609.3 | c.1096G>A | p.(A366T) | SDM | TGGCGAAATCACCCTGTCC | GCGCTGGAGTCGCCCAGG | 0.005 | 1 | 0.449 |
| *PDK2* | 8810 | NM_002611 | c.359G>A | p.(R120Q) | HiTM | GACGCCCTGGTCACCATC | CACGTCGTTGTGCCGGTTC | 0.381 | 0.018 | 25.3 |
| *PEAR1* | 33631 | NM_001080471.1 | c.2471C>T | p.(T824I) | HiTM | TCCAACCCCAGCTACCAC | TGGGGAGCACTGCGACAG | 0.997 | 0.003 | 32 |
| *PELI1* | 8827 | NM_020651 | c.809C>T | p.(A270V) | HiTM | ACCGTGAAGCATTTAGAAG | ATTGATTTCCTGTCTTAAAA | 0.242 | 0.183 | 18.26 |
| *PITX1* | 9004 | NM_002653 | c.724C>T | p.(L242F) | HiTM | TGGCATGCCCAACTCGGG | AGGTTGTTGATGTTGTTGA | 0.986 | 0.01 | 28.2 |
| *PLXDC1* | 20945 | NM_020405 | c.125G>A | p.(R42Q) | HiTM | GCTGCCAAAGGGACCGTG | GGCTCTCCGGTTCCAGCC | 0 | 0.778 | 6.386 |
| *PRKD1* | 9407 | NM_002742 | c.1321C>T | p.(R441W) | HiTM | CAAGGACACGCTGCGGAA | TCCAATCTCCAATAGTGCC | 0.994 | 0.001 | 18.13 |
| *PRPS1L1* | 9463 | NM_175886.2 | c.182G>A | p.(G61D) | HiTM | ATCGTTCAGAGTGGTTGTG | TAGACTGTCGTTGATTTCG | 0.993 | 0.004 | 18.17 |
| *PTK7* | 9618 | NM_152882 | c.1709G>A | p.(R570Q) | HiTM | CATTTTGCCCGGGTGACTC | GTAGTTGCCAGCGTCATCT | 0.964 | 0.09 | 32 |
| *PTPRF* | 9670 | BC048768 | c.1000A>C | p.(S334R) | SDM | AACTGCCACCCGTGTCAC | GTCTCTGTCACCACAAGAT | 0.999 | 0.028 | 19.01 |
| *PXDN* | 14966 | NM_012293 | c.1928G>A | p.(R643Q) | SDM | AACTCAACCCAAACACATT | TATAGCTCTGTCAACAGTC | 0.988 | 0.328 | 19.98 |
| *RALGAPA1* | 17770 | NM_001346243.2 | c.5305del3 | LL1815L | SDM | AGTATATTGGGAATGAATTC | AAGCAATCTGCAATAATAA | - | - | - |
| *SCARB2* | 1665 | NM_005506.3 | c.518T>C | p.(V173A) | HiTM | CTCTTTGTGACTCACACAG | GCCCCAGAGCAATTCGTC | 0.463 | 0.036 | 11.75 |
| *SH2D3C* | 16884 | NM_005489 | c.680G>A | p.(R227Q) | HiTM | TACCATGGCCGCATCCCC | CAAGGTCTCCGAGACCTC | 0.943 | 0.081 | 29.5 |
| *SLC23A1* | 10974 | NM_152685 | c.1393C>A | p.(L465M) | HiTM | CTCTCGCAACCTCTTCGTG | AAGAACATGGAAAATCCCA | 1 | 0.059 | 17.94 |
| *SLC8A2* | 11069 | NM_015063.2 | c.2374G>C | p.(G792R) | HiTM | TGTTGTCTTCGTTGCCCTG | GTGTCAGGGATGGAGGTG | 1 | 0 | 22.1 |
| *SLCO4A1* | 10953 | NM_016354 | c.2035G>A | p.(V679I) | HiTM | CCTGTACAAGGTGCTGGG | GCTATGGCAAAGAAGAGG | 0.001 | 1 | 0.012 |
| *SRCAP* | 16974 | NM_006662.2 | c.5809G>A | p.(G1937S) | SDM | CAGCCCCATCAGCCCTCG | GCAACAGGTTGGGGCAGG | 0.15 | 0.203 | 14.31 |
| *TANC2* | 30212 | NM_025185.3 | c.5066A>G | p.(H1689R) | SDM | ATCTGTCAGCGTGGAGGAT | GGCGCCTGCACTCAGGCT | 0.84 | 0.123 | 10.2 |
| *TRIP12* | 12306 | NM_001284214.1 | c.4928G>A | p.(R1595Q) | SDM | ACTGTGAACCAAGAGGAG | ACGTTTTTTTCTATCCAATC | 0.945 | 0 | 36 |
| *TRPM1* | 7146 | NM_001252024.1 | c.2382C>G | p.(F794L) | SDM | ATGATGATTTGTCGTATCA | ATGTGCGAAATTCCAAAAA | 0.001 | 0.677 | 5.937 |
| *TRPM6* | 17995 | NM_017662.4 | c.6031A>C | p.(T2011P) | SDM | AGCAAGGGAGCCGGGTAG | GGAGGCTCCTCAGCTGAT | 0.526 | 0.214 | 12.22 |
| *TRPM6* | 17995 | NM_017662.4 | c.1922C>A | p.(A641E) | SDM | GCCGTGATTGAGTGTATCC | TTTAACCGTGGCCTCCTC | 1 | 0 | 17.81 |
| *TRPM7* | 17994 | NM_017672.5 | c.1135A>G | p.(T379A) | SDM | GGAGCTTATCGCTGTTTTC | TTTCTTTTCATGCACTCCAT | 0.901 | 0.001 | 26.4 |
| *TSC2* | 12363 | NM_001114382.3 | c.1643G>T | p.(R548M) | SDM | CTGGAAGAAATGGATGTGG | CTCCGGGGGTGGGGAGAG | 0.999 | 0.023 | 19.51 |
| *TSC2* | 12363 | NM_001114382.3 | c.4669C>T | p.(R1557W) | SDM | GGGCCTGGGCTGGCTCAT | GTCAGGAACTCCGTGTAC | 0.958 | 0.022 | 15.98 |
| *USP30* | 20065 | NM_032663 | c.598C>T | p.(P200S) | HiTM | GCAGCAGTCAGAAATAACT | CGGCAGGTAATTTGTTTGG | 0.022 | 0.141 | 14.73 |
