## Supplementary material for "*Drosophila* functional screening of *de novo* variants in autism uncovers deleterious variants and facilitates discovery of rare neurodevelopmental diseases": Table S4

**Supplemental Table 4: Complementation testing of homozygous lethal TG4 lines**

| Fly gene | Deficiency (or Duplication for X-chromsome) | Result | SSC *H. sap* genes | Notes |
|---|---|---|---|---|
| *a* | *Df(2R)Exel6078* | Viable - 2nd site mutation | *PDZD2* | Viable with 2nd DF line *Df(2R)Exel7171* |
| *Abl* | *Df(3L)BSC561* | Lethal | *ABL2* | |
| *Ace* | *Df(3R)Exel6167* | Lethal | *ACHE* | |
| *Ace* | *Df(3R)Exel6167* | Lethal | *BCHE* | |
| *Aldh-III* | *Df(2R)BSC264* | Lethal | *ALDH3A1* | |
| *Ance* | *Df(2L)BSC345* | Viable - 2nd site mutation | *ACE* | Viable with 2nd DF line *Df(2L)BSC253* |
| *arm* | *Dp(1;3)DC446* | Rescued - lethal | *CTNNB1* | X-chromsome |
| *arm* | *Dp(1;3)DC446* | Rescued - lethal | *JUP* | X-chromsome |
| *b* | *Df(2L)BSC252* | Viable - 2nd site mutation | *CSAD* | Viable with 2nd DF line *Df(2L)Exel7059* |
| *CASK* | *Df(3R)Exel6187* | Viable - 2nd site mutation | *CASK* | Viable with 2nd DF line Df(3R)BSC678 |
| *Cat* | *Df(3L)BSC775* | Lethal | *CAT* | |
| *CG11594* | *Df(3L)BSC369* | Viable - 2nd site mutation | *FGGY* | |
| *CG1815* | *Df(3R)BSC505* | Viable - 2nd site mutation | *ZMYND8* | |
| *CG18304* | *Df(2L)Exel7029* | Lethal | *SOGA3* | |
| *CG31637* | *Df(2L)BSC354* | Lethal* | *CHST2* | |
| *CG4562* | *Df(3R)BSC636* | Lethal | *ABCC4* | |
| *CG4562* | *Df(3R)BSC636* | Lethal | *ABCC5* | |
| *CG5521* | *Df(3R)ED6255* | Lethal* | *RALGAPA1* | |
| *CG6767* | *Df(3L)BSC669* | Lethal | PRPS1L1 | |
| *CG7470* | *Df(3L)BSC223* | Lethal | *ALDH18A1* | |
| *CG7744* | *Df(2R)BSC782* | Lethal | *TMEM201* | |
| *CG9098* | *Df(2L)ED385* | Viable - 2nd site mutation | *SH2D3C* | Viable with 2nd DF line *Df(2L)ED354* |
| *ck* | *Df(2L)ED3* | Lethal | *MYO7B* | |
| *ClC-a* | *Df(3R)PS2* | Lethal | *CLCNKB* | |
| *CLIP-190* | *Df(2L)Exel7068* | Viable - 2nd site mutation | *CLIP2* | Viable with 2nd DF line *Df(2L)BSC294* |
| *ctrip* | *Df(3R)ED5066* | Lethal | TRIP12 | |
| *dally* | *Df(3L)ED4413* | Viable - 2nd site mutation | GPC5 | Viable with 2nd DF line *Df(3L)ED4415* |
| *dom* | *Df(2R)BSC821* | Lethal | *EP400* | |
| *dom* | *Df(2R)BSC821* | Lethal | *SRCAP* | |
| *emp* | *Df(2R)BSC608* | Lethal | *SCARB2* | |
| *fry* | *Df(3L)BSC576* | Lethal | *FRYL* | |
| *gig* | *Df(3L)BSC446* | Lethal | *TSC2* | |
| *GluRIB* | *Df(3L)ED4421* | Viable - 2nd site mutation | GRIA1 | Viable with 2nd DF line *Df(3L)BSC390* |
| *Gprk2* | *Df(3R)BSC793* | Lethal | *GRK4* | |
| *if* | *Df(1)BSC725* | Lethal | *ITGA2B* | |
| *if* | *Df(1)BSC725* | Lethal | *ITGA8* | |
| *Lar* | *Df(2L)Exel6044* | Lethal | *PTPRF* | |
| *Lpt* | *Df(2R)BSC136* | Lethal* | *KMT2C* | |
| *Lrch* | *Df(2L)BSC294* | Lethal | LRCH4 | |
| *LRP1* | *Df(2R)Exel6057* | Viable - 2nd site mutation | *LRP1* | Viable with 2nd DF line *Df(2R)Exel6056* |
| *mbc* | *Df(3R)Exel9014* | Lethal | *DOCK1* | |
| *mbl* | *Df(2R)Exel6066* | Lethal | *MBNL1* | |

| | | | | |
|---|---|---|---|---|
| *Mhc* | *Df(2L)BSC325* | Viable - 2nd site mutation | *MYH3* | Viable with 2nd DF line *Df(2L)Exel7067* |
| *Mhc* | *Df(2L)BSC325* | Viable - 2nd site mutation | MYH9 | Viable with 2nd DF line *Df(2L)Exel7067* |
| *msn* | *msn[172] P{ry[+t7.2]=neoFRT}80B* | Lethal | *MINK1* | |
| *msn* | *msn[172] P{ry[+t7.2]=neoFRT}80B* | Lethal | *TNIK* | |
| *Nipped-A* | *Df(2R)BSC630* | Lethal | *TRRAP* | |
| *Nlg3* | *Df(3R)BSC747* | Lethal* | *NLGN3* | |
| *osa* | *Df(3R)BSC790* | Lethal | *ARID1B* | |
| *Pdk* | *Df(2R)BSC408* | Lethal | PDK2 | |
| *PH4alphaEFB* | *Df(3R)Exel6216* | Lethal | *P4HA2* | |
| *Pits* | *Dp(1;3)DC256* | Lethal - rescued by Dp line | *IRF2BPL* | X-chromsome |
| *PMCA* | *Df(4)ED6369* | Lethal | *ATP2B2* | |
| *PMCA* | *Df(4)ED6369* | Lethal | *ATP2B4* | |
| *Ptx1* | *Df(3R)ED6346* | Lethal | *PITX1* | |
| *Pvr* | *Df(2L)ED578* | Lethal | *KDR* | |
| *Pvr* | *Df(2L)ED578* | Lethal | *PDGFRB* | |
| *Pxn* | *Df(3L)BSC119* | Lethal | *PXDN* | |
| *Pxn* | *Df(3L)BSC119* | Lethal | *PXDNL* | |
| *rols* | *Df(3L)ED4475* | Lethal | *TANC2* | |
| *Sema-5c* | *Df(3L)BSC395* | Viable - 2nd site mutation | *SEMA5A* | Viable with 2nd DF line *Df(3L)BSC458* |
| *shot* | *Df(2R)BSC383* | Lethal | *MACF1* | |
| *SIFaR* | *Df(3R)ED10845* | Lethal | *NPFFR2* | |
| *Smr* | *Dp(1;3)DC258* | Lethal - rescued by Dp line | *NCOR1* | X-chromsome |
| *spg* | *Df(3R)BSC789* | Viable - 2nd site mutation | *DOCK4* | Viable with 2nd DF line *Df(2R)BSC305* |
| *Spn* | *Df(3L)BSC116* | Viable - 2nd site mutation | *PPP1R9A* | Viable with 2nd DF line *Df(3L)ED4287* |
| *tok* | *Df(3R)BSC397* | Lethal | *BMP1* | |
| *Trpm* | *Df(2R)ED2426* | Lethal | *TRPM1* | |
| *Trpm* | *Df(2R)ED2426* | Lethal | *TRPM6* | |
| *Trpm* | *Df(2R)ED2426* | Lethal | *TRPM7* | |
| *wb* | *Df(2L)ED793* | Lethal | *LAMA1* | |
| *wb* | *Df(2L)ED793* | Lethal | *LAMA2* | |
| *Yip1d1* | *Df(2L)ED779* | Lethal | *YIPF5* | |
| *Dh44-R2* | *Df(2R)BSC305* | Viable - 2nd site mutation | *CRHR1* | Viable with 2nd DF line *Df(2R)BSC859* |
| *shot* | *Df(2R)BSC383* | Lethal | *DST* | |
| *l(1)G0289* | *Dp(1;3)DC223* | Lethal - rescued by Dp line | *PLXDC1* | X-chromsome |
| *otk* | *Df(2R)BSC199* | Viable - 2nd site mutation | *PTK7* | Viable with 2nd DF line *Df(2R)BSC153* |
| *unc80* | *Df(3R)BSC497* | Lethal | *UNC80* | |
