## Supplementary material for "*Drosophila* functional screening of *de novo* variants in autism uncovers deleterious variants and facilitates discovery of rare neurodevelopmental diseases": Table S5

**Supplemental Table 5: Screen data**

| H. sap gene | D. mel gene | Unique fly peptides | LoF phenotype | UAS-Ref (Rescue) | UAS-Var(Rescue) | Rescue screen - observations | OE screen | OE screen - observations | Combined Screens | pLI | LOEUF | mis O/E | OMIM AD | MIM AD NI | OMIM AR | MIM AR NI | H sap homologs | Chr | Position | Variant | rsID | DbsnpBuil | ncestralAlle | GenomeC | nomAD Fre | ExacFreq | EspAaFreq | EspEaFreq | Transcript | FunctionClass | cDnaVariant | ProteinVariant | PolyPhen(HVar) | SiftScore | CaddScore |
|---|---|---|---|---|---|---|---|---|---|---|---|---|---|---|---|---|---|---|---|---|---|---|---|---|---|---|---|---|---|---|---|---|---|---|---|
| *ABL2* | *Abl* | *9* | Lethal | Y | Y | Ref_Var_significant_difference | No_Pheno | No_Pheno | FD | 0.00 | 0.58 | 0.79 | N | N | N | N | 2 | 1 | 179077107 | C>T | 0 | 0 | C | 0 | 0 | 0 | 0 | 0 | NM_007314.3 | missense | c.3295G>A | p.(A1099T) | 0.998 | 0.013 | 20.9 |
| *CAT* | *Cat* | *1* | Lethal | Y | Y | Ref_Var_significant_difference | No_Pheno | No_Pheno | FD | 0.00 | 1.05 | 0.99 | N | N | Y | N | 1 | 11 | 34475373 | G>A | 0 | 0 | G | 0 | 7.95E-06 | 0 | 0 | 0 | NM_001752.3 | missense | c.611G>A | p.(G204E) | 1 | 0 | 23.1 |
| *CHST2* | *CG31637* | *1* | Lethal | Y | Y | Ref_Var_significant_difference | No_Pheno | No_Pheno | FD | 0.02 | 0.81 | 0.65 | N | N | N | N | 5 | 3 | 142839813 | G>C | 0 | 0 | G | 0 | 0 | 0 | 0 | 0 | NM_004267.4 | missense | c.155G>C | p.(R52P) | 0.989 | 0 | 12.82 |
| *TRPM6* | *Trpm* | *9* | Lethal | Y | Y | Ref_Var_significant_difference | No_Pheno | No_Pheno | FD | 0.00 | 0.44 | 0.84 | N | N | Y | Y | 7 | 9 | 77339567 | T>G | 0 | 0 | T | 0 | 0 | 0 | 0 | 0 | NM_017662.4 | missense | c.6031A>C | p.(T2011P) | 0.526 | 0.214 | 12.22 |
| *TRPM6* | *Trpm* | *9* | Lethal | Y | Y | Ref_Var_significant_difference | No_Pheno | No_Pheno | FD | 0.00 | 0.44 | 0.84 | N | N | Y | Y | 7 | 9 | 77416901 | G>T | 0 | 0 | G | 0 | 0 | 0 | 0 | 0 | NM_017662.4 | missense | c.1922C>A | p.(A641E) | 1 | 0 | 17.81 |
| *TRIP12* | *ctrip* | *9* | Lethal | Y | N | Ref_Var_significant_difference | CP_Tub | CP_Tub | FD | 1.00 | 0.03 | 0.59 | Y | Y | N | N | 1 | 2 | 230650558 | C>T | 0 | 0 | C | 0 | 0 | 0 | 0 | 0 | NM_004238.1 | missense | c.4784G>A | p.(R1595Q) | 0.945 | 0 | 36 |
| *ABCC4* | *CG4562* | *3* | Lethal | Y | Y | Rescue comparable | No_Pheno | No_Pheno | CF | 0.00 | 0.50 | 0.82 | N | N | N | N | 2 | 13 | 95860139 | T>C | 0 | 0 | T | 0 | 0 | 0 | 0 | 0 | NM_005845.3 | missense | c.826A>G | p.(M276V) | 0.654 | 0.024 | 17.16 |
| *ABCC5* | *CG4562* | *3* | Lethal | Y | Y | Rescue comparable | No_Pheno | No_Pheno | CF | 0.02 | 0.38 | 0.65 | N | N | N | N | 2 | 3 | 183667631 | G>A | 200996972 | 137 | G | 0 | 6.41E-05 | 8.20E-05 | 2.37E-04 | 1.18E-04 | NM_005688.2 | missense | c.3137C>T | p.(T1046M) | 0.987 | 0.001 | 20.8 |
| *ABCC5* | *CG4562* | *3* | Lethal | Y | Y | Rescue comparable | No_Pheno | No_Pheno | CF | 0.02 | 0.38 | 0.65 | N | N | N | N | 2 | 3 | 183681319 | G>A | 0 | 0 | G | 0 | 0 | 0 | 0 | 0 | NM_005688.2 | missense | c.2089C>T | p.(R697W) | 1 | 0 | 22.4 |
| *ALDH18A1* | *CG7470* | *1* | Lethal | Y | Y | Rescue comparable | - | - | CF | 0.00 | 0.50 | 0.72 | Y | Y | Y | Y | 1 | 10 | 97371016 | C>G | 0 | 0 | C | 0 | 0 | 0 | 0 | 0 | NM_002860.3 | missense | c.2107G>C | p.(D703H) | 0.007 | 0.012 | 14.45 |
| *BMP1* | *tok* | *1* | Lethal | Y | Y | Rescue comparable | CP_Tub, nub, GMR | CP_Tub, nub, GMR | CF | 0.00 | 0.46 | 0.78 | N | N | Y | N | 3 | 8 | 22067161 | G>A | 147298430 | 134 | G | 0 | 1.2E-05 | 8.00E-06 | 0 | 1.16E-04 | NM_006129.4 | missense | c.2779G>A | p.(G927S) | 0.999 | 0.022 | 32 |
| *GRK4* | *Gprk2* | *1* | Lethal | Y | Y | Rescue comparable | LoF - Tub | LoF - Tub | FD | 0.00 | 1.11 | 1.08 | N | N | N | N | 4 | 4 | 3031020 | C>G | 0 | 0 | C | 0 | 0 | 0 | 0 | 0 | NM_001004057.1 | missense | c.1153C>G | p.(P385A) | 0.999 | 0 | 24.9 |
| *P4HA2* | *PH4alphaEFB* | *1* | Lethal | Y | Y | Rescue comparable | CP_Tub, nub | CP_Tub, nub | CF | 0.00 | 0.67 | 0.86 | Y | Y | N | N | 2 | 5 | 131549620 | C>T | 0 | 0 | C | 0 | 0 | 0 | 0 | 0 | NM_001142599.1 | missense | c.458G>A | p.(G153E) | 1 | 0 | 29.2 |
| *PTPRF* | *Lar* | *7* | Lethal | Y | Y | Rescue comparable | No_Pheno | No_Pheno | CF | 1.00 | 0.20 | 0.79 | N | N | Y | N | 3 | 1 | 44056693 | A>C | 0 | 0 | A,A | 0 | 0 | 0 | 0 | 0 | NM_002840.3 | missense | c.1000A>C | p.(S334R) | 0.999 | 0.028 | 19.01 |
| *PXDN* | *Pxn* | *2* | Lethal | Y | Y | Rescue comparable | No_Pheno | No_Pheno | CF | 0.00 | 0.51 | 0.77 | N | N | Y | Y | 8 | 2 | 1658190 | C>T | 0 | 0 | C,C | 0 | 8.03E-06 | 0 | 0 | 0 | NM_012293.1 | missense | c.1928G>A | p.(R643Q) | 0.988 | 0.328 | 19.98 |
| *TRPM7* | *Trpm* | *9* | Lethal | Y | Y | Rescue comparable | No_Pheno | No_Pheno | CF | 0.00 | 0.55 | 0.79 | Y | Y | N | N | 7 | 15 | 50923683 | T>C | 0 | 0 | T | 0 | 4.04E-06 | 0 | 0 | 0 | NM_017672.4 | missense | c.1135A>G | p.(T379A) | 0.901 | 0.001 | 26.4 |
| *NPFFR2* | *SIFaR* | *1* | Lethal | Y | Y | Rescue comparable | LoF - Tub | LoF - Tub | FD | 0.00 | 1.14 | 1.19 | N | N | N | N | 5 | 4 | 72994491 | G>A | 0 | 0 | G | 0 | 0 | 0 | 0 | 0 | NM_004885.2 | missense | c.489G>A | p.(M163I) | 0.002 | 0.562 | 8.023 |
| *PDK2* | *Pdk* | *3* | Lethal | Y | Y | Rescue comparable | LoF - Tub | LoF - Tub | FD | 0.00 | 0.92 | 0.64 | Y | Y | N | N | 8 | 17 | 48183275 | G>A | 0 | 0 | G | 0 | 4.02E-06 | 0 | 0 | 0 | NM_001199900.1 | missense | c.359G>A | p.(R120Q) | 0.381 | 0.018 | 25.3 |
| *CLCNKB* | *ClC-a* | *7* | Lethal | Y | Y | Rescue comparable | CP_Tub, nub, GMR | CP_Tub, nub, GMR | CF | 0.00 | 1.23 | 1.08 | N | N | Y | N | 5 | 1 | 16374867 | G>C | 0 | 0 | g | 0 | 2.79E-05 | 8.00E-06 | 0 | 0 | NM_000085.4 | missense | c.528G>C | p.(M176I) | 0.001 | 0.564 | 5.89 |
| *ACHE* | *Ace* | *2* | Lethal | N | N | Failed_Rescue | No_Pheno | No_Pheno | NA | 1.00 | 0.21 | 0.60 | N | N | N | N | 2 | 7 | 100488870 | G>A | 0 | 0 | G | 0 | 0 | 0 | 0 | 0 | NM_000665.3 | missense | c.1643C>T | p.(P548L) | 0.847 | 0.183 | 21 |
| *ACHE* | *Ace* | *2* | Lethal | N | N | Failed_Rescue | No_Pheno | No_Pheno | NA | 1.00 | 0.21 | 0.60 | N | N | N | N | 2 | 7 | 100491403 | C>T | 0 | 0 | C | 0 | 0 | 0 | 0 | 0 | NM_015831.2 | missense | c.451G>A | p.(G151R) | 1 | 0 | 27.7 |
| *ALDH3A1* | *Aldh-III* | *9* | Lethal | N | N | Failed_Rescue | No_Pheno | No_Pheno | NA | 0.00 | 1.54 | 0.95 | N | N | N | N | 4 | 17 | 19642319 | G>C | 0 | 0 | G | 0 | 0 | 0 | 0 | 0 | NM_000691.4 | missense | c.1206C>G | p.(F402L) | 0.995 | 0.066 | 14.36 |
| *BCHE* | *Ace* | *2* | Lethal | N | N | Failed_Rescue | No_Pheno | No_Pheno | NA | 0.00 | 1.29 | 1.15 | N | N | Y | N | 2 | 3 | 165547525 | A>C | 0 | 0 | A | 0 | 0 | 0 | 0 | 0 | NM_000055.2 | missense | c.1297T>G | p.(F433V) | 0.727 | 0.033 | 11.96 |
| *CTNNB1* | *arm* | *2* | Lethal | N | N | Failed_Rescue | CP_Tub, nub | CP_Tub, nub | CF | 1.00 | 0.13 | 0.46 | Y | Y | N | N | 2 | 3 | 41275757 | C>T | 0 | 0 | C | 0 | 0 | 0 | 0 | 0 | NM_001904.3 | missense | c.1652C>T | p.(T551M) | 0.1 | 0.069 | 15 |
| *IRF2BPL* | *Pits* | *3* | Lethal | N | N | Failed_Rescue | LoF - Tub, nub | LoF - Tub, nub | FD | 0.96 | 0.33 | 0.87 | Y | Y | N | N | 3 | 14 | 77492034 | GT>G | 0 | 0 | T | 0 | 0 | 0 | 0 | 0 | NM_024496.3 | frameshift | NA | N701X | | | |
| *IRF2BPL* | *Pits* | *3* | Lethal | N | N | Failed_Rescue | LoF - nub, CP tub | LoF - nub, CP tub | FD | 0.96 | 0.33 | 0.87 | Y | Y | N | N | 3 | 14 | 77494046 | G>C | 0 | 0 | G | 0 | 0 | 0 | 0 | 0 | NM_024496.3 | missense | c.90C>G | p.(F30L) | 0.989 | 0.002 | 14.94 |
| *ITGA8* | *if* | *2* | Lethal | N | N | Failed_Rescue | LoF - Tub | LoF - Tub | FD | 0.00 | 0.66 | 1.03 | N | N | Y | N | 4 | 10 | 15634273 | G>A | 0 | 0 | G | 0 | 0.000104 | 1.33E-04 | 0 | 0 | NM_003638.1 | missense | c.2242C>T | p.(R748C) | 0.956 | 0.006 | 24.2 |
| *JUP* | *arm* | *2* | Lethal | N | N | Failed_Rescue | CP_Tub, nub | CP_Tub, nub | CF | 0.00 | 0.61 | 0.83 | Y | N | Y | N | 2 | 17 | 39912444 | T>C | 147628503 | 134 | T | 0 | 9.96E-05 | 8.40E-05 | 0 | 2.33E-04 | NM_021991.2 | missense | c.2069A>G | p.(N690S) | 0.039 | 0.553 | 2.456 |
| *KDR* | *Pvr* | *11* | Lethal | N | N | Failed_Rescue | No_Pheno | No_Pheno | NA | 1.00 | 0.25 | 0.88 | N | N | N | N | 8 | 4 | 55953925 | C>T | 0 | 0 | C | 0 | 0 | 0 | 0 | 0 | NM_002253.2 | issense-near-splic | c.3511G>A | p.(D1171N) | 0.697 | 0.004 | 25.9 |
| *MBNL1* | *mbl* | *18* | Lethal | N | N | Failed_Rescue | CP_Tub, nub | CP_Tub, nub | CF | 0.75 | 0.42 | 0.49 | N | N | N | N | 3 | 3 | 152018116 | T>C | 0 | 0 | T | 0 | 0 | 0 | 0 | 0 | NM_207293.1 | missense | c.134T>C | p.(V45A) | 0.843 | 0.019 | 15.52 |
| *MINK1* | *msn* | *6* | Lethal | N | N | Failed_Rescue | LoF - Tub | LoF - Tub | FD | 1.00 | 0.13 | 0.57 | N | N | N | N | 3 | 17 | 4789777 | T>C | 0 | 0 | T | 0 | 0 | 0 | 0 | 0 | NM_015716.4 | missense | c.805T>C | p.(C269R) | 0.996 | 0 | 20.9 |
| *NCOR1* | *Smr* | *2* | Lethal | N | N | Failed_Rescue | CP_Tub, nub, GMR | CP_Tub, nub, GMR | CF | 1.00 | 0.16 | 0.68 | N | N | N | N | 2 | 17 | 16024513 | G>A | 0 | 0 | G | 0 | 1.19E-05 | 1.60E-05 | 0 | 0 | NM_006311.3 | missense | c.1705C>T | p.(P569S) | 0.998 | 0.575 | 13.24 |
| *PDGFRB* | *Pvr* | *11* | Lethal | N | N | Failed_Rescue | CP_Tub | CP_Tub | CF | 0.75 | 0.34 | 0.81 | Y | Y | N | N | 8 | 5 | 149512344 | C>T | 0 | 0 | C | 0 | 1.21E-05 | 2.90E-05 | 0 | 1.16E-04 | NM_002609.3 | missense | c.1096G>A | p.(A366T) | 0.005 | 1 | 0.449 |
| *PITX1* | *Ptx1* | *3* | Lethal | N | N | Failed_Rescue | CP_Tub, nub, GMR | CP_Tub, nub, GMR | CF | 0.89 | 0.41 | 0.73 | Y | N | N | N | 3 | 5 | 134364690 | G>A | 0 | 0 | G | 0 | 0 | 0 | 0 | 0 | NM_002653.4 | missense | c.724C>T | p.(L242F) | 0.986 | 0.01 | 28.2 |
| *RALGAPA1* | *CG5521* | *2* | Lethal | N | N | Failed_Rescue | No_Pheno | No_Pheno | NA | 1.00 | 0.24 | 0.68 | N | N | Y | Y | 2 | 14 | 36096328 | TAAG>T | 0 | 0 | A | 0 | 0 | 0 | 0 | 0 | NM_001283044.1 | codingComplex | NA | LL1815L | | | |
| *SCARB2* | *emp* | *3* | Lethal | N | N | Failed_Rescue | CP_Tub | CP_Tub | CF | 0.00 | 0.71 | 0.76 | N | N | Y | Y | 3 | 4 | 77100764 | A>G | 0 | 0 | A | 0 | 0 | 0 | 0 | 0 | NM_005506.3 | missense | c.518T>C | p.(V173A) | 0.463 | 0.036 | 11.75 |
| *SRCAP* | *dom* | *4* | Lethal | N | N | Failed_Rescue | No_Pheno | No_Pheno | NA | 1.00 | 0.09 | 0.86 | Y | Y | N | N | 2 | 16 | 30740437 | G>A | 0 | 0 | G | 0 | 1.59E-05 | 1.60E-05 | 0 | 0 | NM_006662.2 | missense | c.5809G>A | p.(G1937S) | 0.15 | 0.203 | 14.31 |
| *SRCAP* | *dom* | *4* | Lethal | N | N | Failed_Rescue | No_Pheno | No_Pheno | NA | 1.00 | 0.09 | 0.86 | Y | Y | N | N | 2 | 16 | 30740738 | ATGCC>A | 0 | 0 | N | 0 | 0 | 0 | 0 | 0 | NM_006662.2 | frameshift | c.5973_5976del4 | p.(C1993Tfs*42) | | | |
| *TANC2* | *rols* | *7* | Lethal | N | N | Failed_Rescue | No_Pheno | No_Pheno | NA | 1.00 | 0.16 | 0.80 | N | N | N | N | 2 | 17 | 61482569 | C>T | 0 | 0 | C | 0 | 0 | 0 | 0 | 0 | NM_025185.3 | stop-gained | c.3196C>T | p.(R1066*) | | | |
| *TANC2* | *rols* | *7* | Lethal | N | N | Failed_Rescue | No_Pheno | No_Pheno | NA | 1.00 | 0.16 | 0.80 | N | N | N | N | 2 | 17 | 61498409 | A>G | 0 | 0 | A | 0 | 0 | 0 | 0 | 0 | NM_025185.3 | missense | c.5066A>G | p.(H1689R) | 0.84 | 0.123 | 10.2 |
| *TRPM1* | *Trpm* | *9* | Lethal | N | N | Failed_Rescue | No_Pheno | No_Pheno | NA | 0.00 | 1.03 | 0.99 | N | N | Y | Y | 7 | 15 | 31330304 | G>C | 0 | 0 | G | 0 | 0 | 0 | 0 | 0 | NM_001252024.1 | missense | c.2382C>G | p.(F794L) | 0.001 | 0.677 | 5.937 |
| *TSC2* | *gig* | *2* | Lethal | N | N | Failed_Rescue | CP_Tub | CP_Tub | NA | 1.00 | 0.07 | 1.01 | Y | Y | N | N | 1 | 16 | 2115563 | G>T | 0 | 0 | g | 0 | 0 | 0 | 0 | 0 | NM_001077183.1 | missense | c.1643G>T | p.(R548M) | 0.999 | 0.023 | 19.51 |
| *TSC2* | *gig* | *2* | Lethal | N | N | Failed_Rescue | LoF - Tub | LoF - Tub | FD | 1.00 | 0.07 | 1.01 | Y | Y | N | N | 1 | 16 | 2136269 | C>T | 0 | 0 | C | 0 | 8E-06 | 1.70E-05 | 0 | 0 | NM_001114382.1 | missense | c.4669C>T | p.(R1557W) | 0.958 | 0.022 | 15.98 |
| *PRPS1L1* | *CG6767* | *6* | Lethal | N | N | Failed_Rescue | No_Pheno | No_Pheno | NA | 0.23 | 0.87 | 1.09 | Y | N | N | N | 3 | 7 | 18067224 | C>T | 0 | 0 | C | 0 | 0 | 0 | 0 | 0 | NM_175886.2 | missense | c.182G>A | p.(G61D) | 0.993 | 0.004 | 18.17 |
| *ATP2B2* | *PMCA* | *7* | Lethal | - | - | - | LoF - nub | LoF - nub | FD | 1.00 | 0.54 | 0.15 | N | N | Y | Y | 4 | 3 | 10387773 | G>A | 0 | 0 | G | 0 | 0 | 0 | 0 | 0 | NM_001001331.2 | missense | c.2453C>T | p.(T818M) | 0.997 | 0 | 29.7 |
| *ACE* | *Ance* | *1* | Viable | - | - | ND | GoF - Tub, CP_nub | GoF - Tub, CP_nub | FD | 0.00 | 1.08 | 1.06 | N | N | Y | Y | 2 | 17 | 61566305 | A>G | 0 | 0 | A | 0 | 2.39E-05 | 1.60E-05 | 0 | 0 | NM_000789.3 | missense | c.2453A>G | p.(Y818C) | 0.954 | 0.003 | 7.512 |
| *GLRA2* | *GluClalpha* | *11* | Copulation defects | N | Copulation defects | Ref_Var_significant_difference | LoF - Tub | LoF - Tub | FD | 0.98 | 0.30 | 0.43 | N | N | N | N | 2 | X | 14599441 | A>G | 0 | 0 | A | 0 | 0 | 0 | 0 | 0 | NM_001118885.1 | missense | c.407A>G | p.(N136S) | 0.999 | 0 | 25.1 |
| *GPC5* | *dally* | *1* | ND | - | - | ND | GoF - Tub | GoF - Tub | FD | 0.00 | 1.08 | 1.09 | N | N | N | N | 2 | 13 | 92345513 | T>C | 0 | 0 | T | 0 | 0 | 0 | 0 | 0 | NM_004466.4 | missense | c.398T>C | p.(M133T) | 0.978 | 0.001 | 18.43 |
| *KCND3* | *Shal* | *3* | Copulation defects | N | ent and decreased g | Ref_Var_significant_difference | LoF - Tub, nub | LoF - Tub, nub | FD | 0.99 | 0.27 | 0.48 | Y | Y | N | N | 2 | 1 | 112525092 | C>G | 0 | 0 | C | 0 | 0 | 0 | 0 | 0 | NM_004980.4 | missense | c.257G>C | p.(R86P) | 1 | 0.001 | 22.9 |
| *MYH9* | *Mhc* | *20* | ND | - | - | ND | GoF - Tub, nub | GoF - Tub, nub | FD | 1.00 | 0.09 | 0.70 | Y | Y | N | N | 3 | 22 | 36684831 | C>T | 0 | 0 | C | 0 | 0 | 0 | 0 | 0 | NM_002473.4 | missense | c.4712G>A | p.(R1571Q) | 0.973 | 0.106 | 36 |
| *PC* | *PCB* | *2* | Courtship defects | - | - | ND | GoF - Tub | GoF - Tub | FD | 0.01 | 0.45 | 0.69 | N | N | Y | Y | 2 | 11 | 66617104 | G>C | 0 | 0 | G | 0 | 0 | 0 | 0 | 0 | NM_000920.3 | missense | c.3125C>G | p.(P1042R) | 0.995 | 0.047 | 16.69 |
| *SLC23A1* | *CG6293* | *1* | Copulation defects | Y | Y | Ref and Var Rescue | GoF - Tub | GoF - Tub | FD | 0.07 | 0.52 | 0.69 | N | N | N | N | 3 | 5 | 138713736 | G>T | 0 | 0 | G | 0 | 0 | 0 | 0 | 0 | NM_152685.3 | missense | c.1393C>A | p.(L465M) | 1 | 0.059 | 17.94 |
| *EPHA1* | *Eph* | *5* | ND | - | - | ND | LoF - nub | LoF - nub | FD | 0.00 | 1.05 | 0.96 | N | N | N | N | 2 | 7 | 143094667 | C>T | 0 | 0 | C | 0 | 6.76E-05 | 9.80E-05 | 0 | 0 | NM_005232.4 | missense | c.1699G>A | p.(V567I) | 0 | 0.773 | 0.038 |
| *MAP4K1* | *hppy* | *5* | Copulation, Movement & Grooming defects | - | - | ND | Complex_LoF_GMR, GoF_nub | Complex_LoF_GMR, GoF_nub | FD | 0.99 | 0.30 | 0.57 | N | N | N | N | 8 | 19 | 39086584 | A>G | 0 | 0 | A | 0 | 0 | 0 | 0 | 0 | NM_007181.4 | missense | c.2174T>C | p.(M725T) | 0.111 | 0.059 | 13.81 |
| *BAIAP2L1* | *IRSp53* | *4* | ND | - | - | ND | GoF - nub, CP_Tub | GoF - nub, CP_Tub | FD | 0.00 | 0.68 | 0.90 | N | N | N | N | 4 | 7 | 97923404 | G>A | 0 | 0 | G | 0 | 0 | 0 | 0 | 0 | NM_018842.4 | missense | c.1442C>T | p.(A481V) | 0.003 | 0.164 | 9.777 |
| *HTR1D* | *5-HT1B* | *1* | Copulation defects | N | N | Comparable_Effects | GoF -nub, GMR, CP_tub | GoF -nub, GMR, CP_tub | FD | 0.00 | 1.30 | 0.98 | N | N | N | N | 4 | 1 | 23520417 | G>T | 0 | 0 | G | 0 | 0 | 0 | 0 | 0 | NM_000864.4 | missense | c.296C>A | p.(T99N) | 0.99 | 0.076 | 14.42 |
| *PLXDC1* | *l(1)G0289* | *1* | ND | - | - | ND | CP nub , CP_Tub | CP nub , CP_Tub | NA | 0.00 | 0.88 | 0.84 | N | N | N | N | 2 | 17 | 37296037 | C>T | 0 | 0 | C | 0 | 1.2E-05 | 1.60E-05 | 0 | 0 | NM_020405.4 | missense | c.125G>A | p.(R42Q) | 0 | 0.778 | 6.386 |
| *EPHB1* | *Eph* | *5* | ND | - | - | ND | Complex_LoF - GMR, GoF_nub | Complex_LoF - GMR, GoF_nub | FD | 1.00 | 0.23 | 0.72 | N | N | N | N | 2 | 3 | 134968233 | G>A | 0 | 0 | G | 0 | 3.61E-05 | 2.50E-05 | 0 | 0 | NM_004441.4 | missense | c.2746G>A | p.(V916M) | 0.999 | 0.011 | 32 |
| *AK1* | *Adk1* | *1* | Copulation & Grooming defects | - | - | ND | No_Pheno | No_Pheno | CF | 0.03 | 0.92 | 0.82 | N | N | Y | Y | 2 | 9 | 130635003 | G>A | 138798179 | 134 | G | 0 | 0.001667 | 0.001948 | 0 | 0 | NM_000476.2 | missense | c.173C>T | p.(S58L) | 0.261 | 0.08 | 17.16 |
| *ALDH1L1* | *CG8665* | *1* | Increased courtship | N | courship and increas | Ref_Var_significant_difference | No_Pheno | No_Pheno | FD | 0.00 | 0.77 | 0.93 | N | N | N | N | 1 | 3 | 125822675 | T>G | 0 | 0 | T | 0 | 0 | 0 | 0 | 0 | NM_001270364.1 | missense | c.2698A>C | p.(N900H) | 0.133 | 0.152 | 9.814 |
| *BEST3* | *Best1* | *4* | ND | - | - | ND | No_Pheno | No_Pheno | CF | 0.00 | 1.37 | 0.91 | N | N | N | N | 2 | 12 | 70087547 | G>T | 0 | 0 | G | 0 | 0 | 0 | 0 | 0 | NM_032735.2 | missense | c.388C>A | p.(R130S) | 1 | 0 | 23.2 |
| *DPYSL2* | *CRMP* | *3* | ND | - | - | ND | No_Pheno | No_Pheno | CF | 0.99 | 0.27 | 0.44 | N | N | N | N | 2 | 8 | 26510772 | C>T | 0 | 0 | C | 0 | 3.98E-06 | 8.00E-06 | 0 | 0 | NM_001386.5 | missense | c.1486C>T | p.(R496C) | 0.992 | 0.004 | 23.3 |
| *DPYSL3* | *CRMP* | *3* | ND | - | - | ND | No_Pheno | No_Pheno | CF | 1.00 | 0.22 | 0.65 | N | N | N | N | 2 | 5 | 146804493 | C>T | 0 | 0 | C | 0 | 2.41E-05 | 8.00E-06 | 0 | 1.22E-04 | NM_001197294.1 | missense | c.415G>A | p.(V139I) | 0.099 | 0.15 | 15.81 |
| *ELAVL3* | *fne* | *2* | ND | - | - | ND | CP_Tub | CP_Tub | NA | 0.80 | 0.45 | 0.47 | N | N | N | N | 2 | 19 | 11569032 | A>G | 0 | 0 | A | 0 | 0 | 0 | 0 | 0 | NM_001420.3 | missense | c.557T>C | p.(L186P) | 0.998 | 0 | 18.85 |
| *GCLC* | *Gclc* | *1* | ND | - | - | ND | No_Pheno | No_Pheno | CF | 0.57 | 0.39 | 0.60 | N | N | Y | N | 2 | 6 | 53385640 | G>A | 0 | 0 | G | 0 | 0 | 0 | 0 | 0 | NM_001498.3 | missense | c.382C>T | p.(R128W) | 0.999 | 0 | 23.1 |
| *KDM2A* | *Kdm2* | *1* | Increased courtship | Y | also Increased copul | Ref rescue, Ref_Var_significant_difference | No_Pheno | No_Pheno | FD | 1.00 | 0.05 | 0.43 | N | N | N | N | 2 | 11 | 66999298 | G>A | 0 | 0 | G | 0 | 0 | 0 | 0 | 0 | NM_012308.2 | missense | c.1346G>A | p.(R449K) | 0 | 1 | 5.746 |
| *PRKD1* | *PKD* | *3* | ND | - | - | ND | No_Pheno | No_Pheno | CF | 0.00 | 0.64 | 0.80 | Y | N | N | N | 3 | 14 | 30102146 | G>A | 0 | 0 | G | 0 | 1.2E-05 | 1.60E-05 | 0 | 0 | NM_002742.2 | missense | c.1321C>T | p.(R441W) | 0.994 | 0.001 | 18.13 |
| *USP30* | *Usp30* | *1* | Increased courtship | Y | out decreased groomi | Ref_Var_significant_difference | No_Pheno | No_Pheno | FD | 0.00 | 0.66 | 0.74 | N | N | N | N | 1 | 12 | 109510128 | C>T | 0 | 0 | C | 0 | 0 | 0 | 0 | 0 | NM_032663.3 | missense | c.598C>T | p.(P200S) | 0.022 | 0.141 | 14.73 |
| *CLIP2* | *CLIP-190* | *15* | ND | - | - | ND | CP_nub | CP_nub | NA | 1.00 | 0.20 | 0.72 | N | N | N | N | 5 | 7 | 73731913 | G>T | 0 | 0 | G | 0 | 5.42E-06 | 9.00E-06 | 0 | 0 | NM_003388.4 | missense | c.37G>T | p.(G13W) | 0.958 | 0.015 | 17.36 |
| *CSAD* | *b* | *2* | ND | - | - | ND | No_Pheno | No_Pheno | CF | 0.00 | 1.01 | 0.94 | N | N | N | N | 5 | 12 | 53553919 | G>A | 0 | 0 | G | 0 | 0 | 0 | 0 | 0 | NM_015989.4 | missense | c.1232C>T | p.(A411V) | 0.796 | 0.756 | 11.5 |
| *GRIA1* | *GluRIB* | *3* | ND | - | - | ND | CP_Tub | CP_Tub | NA | 1.00 | 0.28 | 0.56 | N | N | N | N | 2 | 5 | 153030052 | G>A | 148008926 | 134 | G | 0 | 7.58E-05 | 9.00E-05 | 0 | 0 | NM_001258021.1 | missense | c.653G>A | p.(R218H) | 0.997 | 0.021 | 29.8 |
| *LRCH4* | *Lrch* | *3* | ND | - | - | ND | No_Pheno | No_Pheno | CF | 0.00 | 0.70 | 1.02 | N | N | N | N | 8 | 7 | 100183600 | C>T | 0 | 0 | C | 0 | 0 | 0 | 0 | 0 | NM_002319.3 | missense | c.124G>A | p.(V42M) | 0.891 | 0.002 | 22.5 |
| *MEGF11* | *drpr* | *6* | No Significant Behavior Defects | - | - | ND | CP_Tub | CP_Tub | NA | 0.00 | 1.12 | 0.88 | N | N | N | N | 3 | 15 | 66198459 | G>A | 201758507 | 137 | G | 2 | 7.33E-05 | 1.26E-04 | 0 | 0 | NM_032445.2 | missense | c.2731C>T | p.(R911C) | 0.365 | 0.062 | 16.38 |
| *PELI1* | *Pli* | *1* | ND | - | - | ND | No_Pheno | No_Pheno | CF | 0.42 | 0.50 | 0.63 | N | N | N | N | 3 | 2 | 64322284 | G>A | 0 | 0 | G | 0 | 0 | 0 | 0 | 0 | NM_020651.3 | missense | c.809C>T | p.(A270V) | 0.242 | 0.183 | 18.26 |
| *MADD* | *Rab3-GEF* | 10 | Increased courtship | Y | Y | Ref and Var Rescue | ND | ND | CF | 0.00 | 0.58 | 0.89 | N | N | Y | Y | 1 | 11 | 47304002 | C>T | 0 | 0 | C | 0 | 0 | 0 | 0 | 0 | NM_130471.2 | missense | c.1540C>T | p.(R514C) | 1 | 0 | 29.8 |
