## Supplementary material for "*Drosophila* functional screening of *de novo* variants in autism uncovers deleterious variants and facilitates discovery of rare neurodevelopmental diseases": Table S6

| Supplemental Table 6: List of essential and viable TG4 mutants and corresponding human gen | | | | | |
|---|---|---|---|---|---|
| ***H. sap* gene** | ***D. mel* gene** | **Essential** | **pLI** | **LOEUF** | **Mis O/E** |
| *ABL2* | *Abl* | Y | 0.00 | 0.58 | 0.79 |
| *ACHE* | *Ace* | Y | 1.00 | 0.21 | 0.60 |
| *BCHE* | *Ace* | Y | 0.00 | 1.29 | 1.15 |
| *ALDH3A1* | *Aldh-III* | Y | 0.00 | 1.54 | 0.95 |
| *CTNNB1* | *arm* | Y | 1.00 | 0.13 | 0.46 |
| *JUP* | *arm* | Y | 0.00 | 0.61 | 0.83 |
| *CAT* | *Cat* | Y | 0.00 | 1.05 | 0.99 |
| *CHST2* | *CG31637* | Y | 0.02 | 0.81 | 0.65 |
| *ABCC4* | *CG4562* | Y | 0.00 | 0.50 | 0.82 |
| *ABCC5* | *CG4562* | Y | 0.02 | 0.38 | 0.65 |
| *RALGAPA1* | *CG5521* | Y | 1.00 | 0.24 | 0.68 |
| PRPS1L1 | *CG6767* | Y | 0.23 | 0.87 | 1.09 |
| *ALDH18A1* | *CG7470* | Y | 0.00 | 0.50 | 0.72 |
| *CLCNKB* | *ClC-a* | Y | 0.00 | 1.23 | 1.08 |
| TRIP12 | *ctrip* | Y | 1.00 | 0.03 | 0.59 |
| *SRCAP* | *dom* | Y | 1.00 | 0.09 | 0.86 |
| *SCARB2* | *emp* | Y | 0.00 | 0.71 | 0.76 |
| *TSC2* | *gig* | Y | 1.00 | 0.07 | 1.01 |
| *GRK4* | *Gprk2* | Y | 0.00 | 1.11 | 1.08 |
| *ITGA8* | *if* | Y | 0.00 | 0.66 | 1.03 |
| *PTPRF* | *Lar* | Y | 1.00 | 0.20 | 0.79 |
| LRCH4 | *Lrch* | Y | 0.00 | 0.70 | 1.02 |
| *MBNL1* | *mbl* | Y | 0.75 | 0.42 | 0.49 |
| *MINK1* | *msn* | Y | 1.00 | 0.13 | 0.57 |
| PDK2 | *Pdk* | Y | 0.00 | 0.92 | 0.64 |
| *P4HA2* | *PH4alphaEFB* | Y | 0.00 | 0.67 | 0.86 |
| *IRF2BPL* | *Pits* | Y | 0.96 | 0.33 | 0.87 |
| *PITX1* | *Ptx1* | Y | 0.89 | 0.41 | 0.73 |
| *KDR* | *Pvr* | Y | 1.00 | 0.25 | 0.88 |
| *PDGFRB* | *Pvr* | Y | 0.75 | 0.34 | 0.81 |
| *PXDN* | *Pxn* | Y | 0.00 | 0.51 | 0.77 |
| *TANC2* | *rols* | Y | 1.00 | 0.16 | 0.80 |
| *NPFFR2* | *SIFaR* | Y | 0.00 | 1.14 | 1.19 |
| *NCOR1* | *Smr* | Y | 1.00 | 0.16 | 0.68 |
| *BMP1* | *tok* | Y | 0.00 | 0.46 | 0.78 |
| *TRPM1* | *Trpm* | Y | 0.00 | 1.03 | 0.99 |
| *TRPM6* | *Trpm* | Y | 0.00 | 0.44 | 0.84 |
| *TRPM7* | *Trpm* | Y | 0.00 | 0.55 | 0.79 |
| *PLXDC1* | *l(1)G0289* | Y | 0.00 | 0.88 | 0.84 |
| *UNC80* | *unc80* | Y | 0.18 | 0.30 | 0.62 |
| *ACE* | *Ance* | N | 0.00 | 1.08 | 1.06 |
| *AK1* | *Adk1* | N | 0.03 | 0.92 | 0.82 |
| *ALDH1L1* | *CG8665* | N | 0.00 | 0.77 | 0.93 |

| | | | | | |
|---|---|---|---|---|---|
| *ATP10A* | *CG33298* | N | 0.00 | 0.46 | 0.92 |
| *BAIAP2L1* | *IRSp53* | N | 0.00 | 0.68 | 0.90 |
| *BEST3* | *Best1* | N | 0.00 | 1.37 | 0.91 |
| *CASK* | *CASK* | N | 1.00 | 0.07 | 0.36 |
| *CEP135* | *Cep135* | N | 0.00 | 0.82 | 0.98 |
| *CLIP2* | *CLIP-190* | N | 1.00 | 0.20 | 0.72 |
| *CSAD* | *b* | N | 0.00 | 1.01 | 0.94 |
| *DDR2* | *Ddr* | N | 0.88 | 0.34 | 0.68 |
| *DLC1* | *cv-c* | N | 1.00 | 0.24 | 1.27 |
| *DPP6* | *CG17684* | N | 0.35 | 0.38 | 0.68 |
| *DPYSL2* | *CRMP* | N | 0.99 | 0.27 | 0.44 |
| *DPYSL3* | *CRMP* | N | 1.00 | 0.22 | 0.65 |
| *ELAVL3* | *fne* | N | 0.80 | 0.45 | 0.47 |
| *EPHA1* | *Eph* | N | 0.00 | 1.05 | 0.96 |
| *EPHB1* | *Eph* | N | 1.00 | 0.23 | 0.72 |
| *FGGY* | *CG11594* | N | 0.00 | 1.22 | 1.01 |
| *GCLC* | *Gclc* | N | 0.57 | 0.39 | 0.60 |
| *GLRA2* | *GluClalpha* | N | 0.98 | 0.30 | 0.43 |
| *GNAO1* | *Galphao* | N | 0.99 | 0.26 | 0.41 |
| *GPC5* | *dally* | N | 0.00 | 1.08 | 1.09 |
| *GRIA1* | *GluRIB* | N | 1.00 | 0.28 | 0.56 |
| *HTR1D* | *5-HT1B* | N | 0.00 | 1.30 | 0.98 |
| *IGF2R* | *Lerp* | N | 1.00 | 0.22 | 0.84 |
| *KCND3* | *Shal* | N | 0.99 | 0.27 | 0.48 |
| *KDM2A* | *Kdm2* | N | 1.00 | 0.05 | 0.43 |
| *MADD* | *Rab3-GEF* | N | 0.00 | 0.58 | 0.89 |
| *MANBA* | *beta-Man* | N | 0.00 | 0.94 | 0.97 |
| *MAP4K1* | *hppy* | N | 0.99 | 0.30 | 0.57 |
| *MEGF11* | *drpr* | N | 0.00 | 1.12 | 0.88 |
| *MYH3* | *Mhc* | N | 0.00 | 0.62 | 0.83 |
| *MYH9* | *Mhc* | N | 1.00 | 0.09 | 0.70 |
| *NID2* | *Ndg* | N | 0.00 | 0.53 | 1.02 |
| *NLGN1* | *Nlg3* | N | 0.88 | 0.36 | 0.70 |
| *NOS3* | *Nos* | N | 0.00 | 0.50 | 0.78 |
| *NTN1* | *NetA* | N | 1.00 | 0.14 | 0.64 |
| *NTN5* | *NetB* | N | 0.00 | 1.73 | 0.79 |
| *PC* | *PCB* | N | 0.01 | 0.45 | 0.69 |
| *PEAR1* | *drpr* | N | 0.00 | 0.90 | 0.92 |
| *PELI1* | *Pli* | N | 0.42 | 0.50 | 0.63 |
| *PPP1R9A* | *Spn* | N | 0.00 | 0.48 | 0.86 |
| *PRKD1* | *PKD* | N | 0.00 | 0.64 | 0.80 |
| *RIMS2* | *Rim* | N | 1.00 | 0.21 | 0.92 |
| *SDK2* | *sdk* | N | 0.02 | 0.34 | 0.78 |
| *SEC14L5* | *retm* | N | 0.00 | 1.22 | 1.20 |
| *SH2D3C* | *CG9098* | N | 0.00 | 0.58 | 0.76 |

| | | | | | |
|---|---|---|---|---|---|
| *SLC23A1* | *CG6293* | N | 0.07 | 0.52 | 0.69 |
| *TULP4* | *Tusp* | N | 1.00 | 0.13 | 0.87 |
| *USP30* | *Usp30* | N | 0.00 | 0.66 | 0.74 |
| *ZMYND8* | *CG1815* | N | 1.00 | 0.08 | 0.57 |
| *ATP2B2* | *PMCA* | Y | 1.00 | 0.54 | 0.15 |
