## Supplementary material for "*Drosophila* functional screening of *de novo* variants in autism uncovers deleterious variants and facilitates discovery of rare neurodevelopmental diseases": Figure S

### Supplemental Information

**Figure S1: Lethality of *TG4* mutants and its implication on essentiality of *Drosophila* genes**

(A) Assessment of lethal *TG4* lines by complementation tests with corresponding deficiency (Df) lines. Lethality of 17 TG4 lines were due to a second site mutations, potentially present in the original MiMIC line or introduced during RMCE. (B) Table summarizing the tools generated in this study in respect to essentiality of fly genes. (C-E) Gene level statistics for human homologs of essential and non-essential fly genes. No significant differences were found with t-test analysis.

### Figure S1:

A

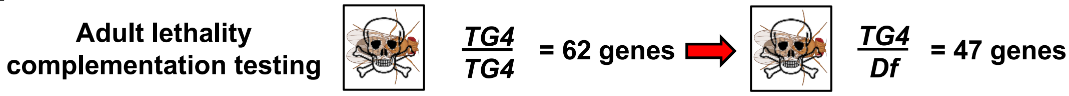


B

| | *D. mel* LoF | *H. sap* transgenics | |
|---|---|---|---|
| | ***TG4*** **lines (Corresponding # *H. sap* genes)** | **Ref *UAS-cDNA*** | **Var *UAS-cDNA*** |
| **essential genes** | 47 (60) | 45 | 49 |
| **non-essential genes** | 62 (68) | 52 | 34 |
| **failed MiMIC conversions** | 13 (15) | 9 | 7 |
| **Total fly lines made** | 109* (128) | 106 | 90 |

*36 lines are novel since publication of Lee *et al.*, 2018

C D E

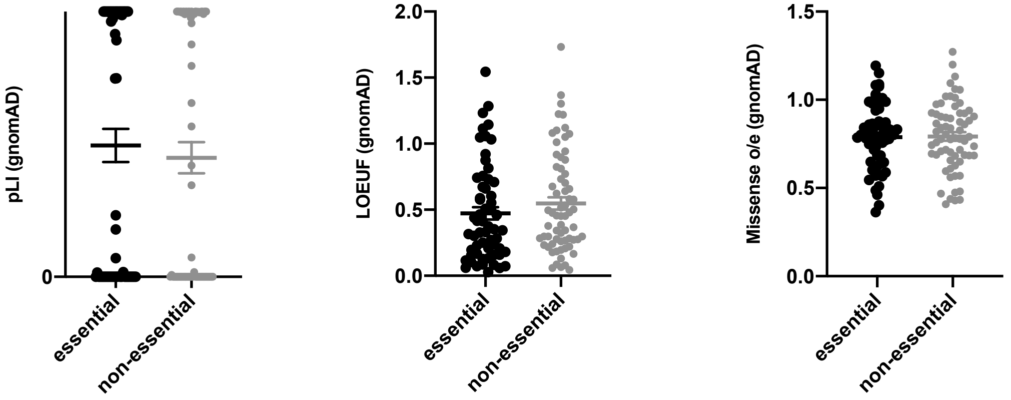

**Figure S2: Expression of TG4 lines corresponding to hits from the humanization screen corresponding to essential fly genes in larval brains**

Third instar larval brain imaging of *TG4* in *Abl*, *Cat*, *CG31637*, *ctrip* and *Trpm* driving UAS-nlsGFP (green in top row) co-stained with neuronal (Elav) and glial (Repo) nuclear markers displayed using corresponding colocalization channels to indicate neuronal and/or glial expression (white in the bottom two rows). Scale bar = 25 μm.

### Figure S2:

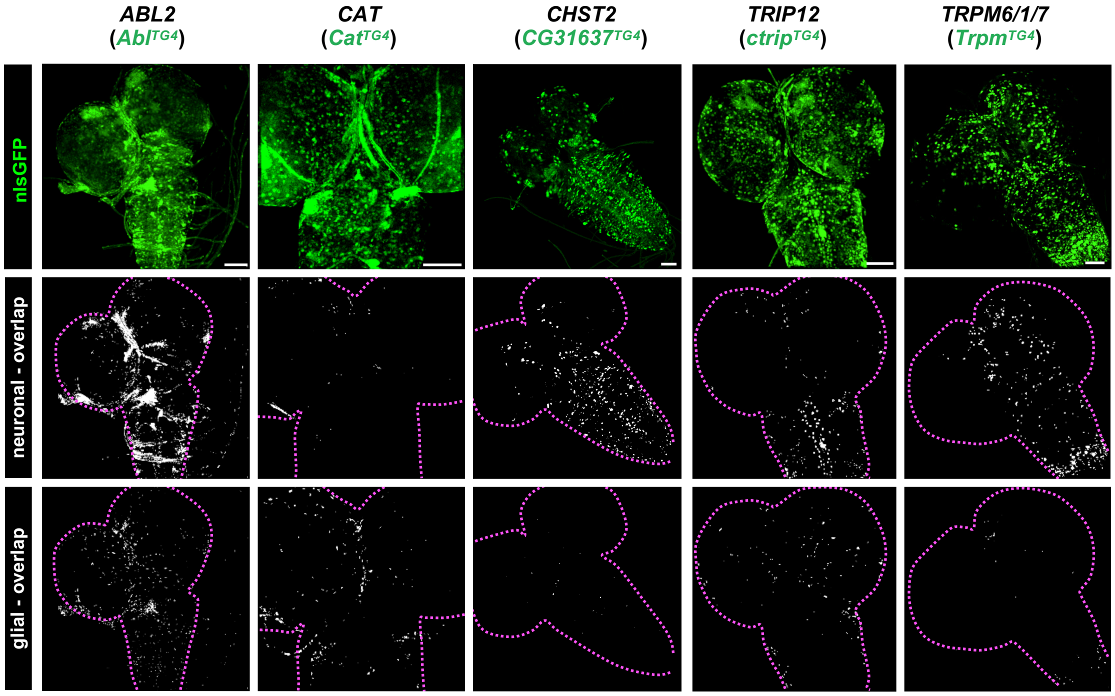
ABL2
(AblTG4)
CAT
(CatTG4)
CHST2
(CG31637TG4)
TRIP12
(ctripTG4)
TRPM6/1/7
(TrpmTG4)
nlsGFP
neuronal - overlap
glial - overlap

**Figure S3: Additional behavior data for viable *TG4* fly mutants**

(A-D) For non-essential *TG4* mutants that we were unable to humanized due to technical reasons, we also assessed the number of frames male flies spent performing single-wing extensions (courtship), copulating, moving within the chamber, or grooming during a test period. The red line represents the average number of frames a *Canton-S* male spends doing the same activity. *p<0.05, **p<0.01, ***p<0.001, ****p<0.0001.

**Figure S3:**

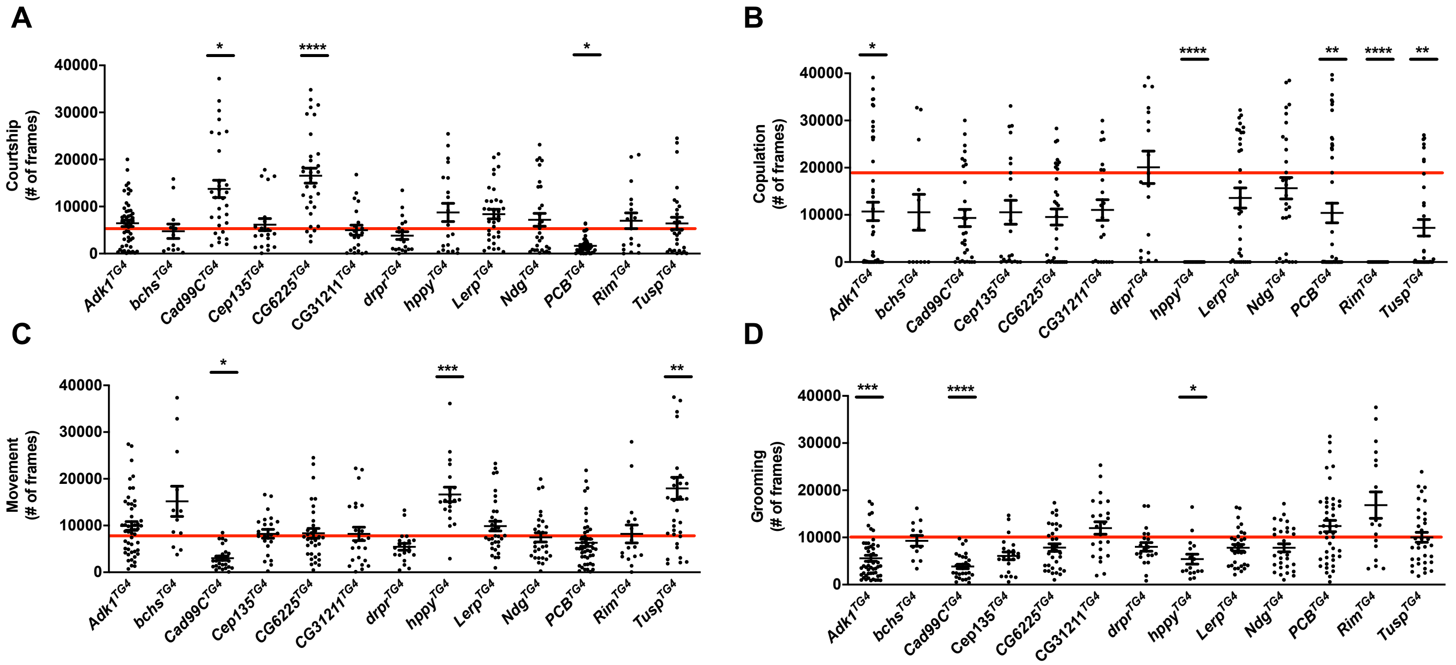
A
B
C
D
Courtship (# of frames)
Copulation (# of frames)
Movement (# of frames)
Grooming (# of frames)
40000
30000
20000
10000
0
Adk1TG4
bchsTG4
Cad99CTG4
Cep135TG4
CG6225TG4
CG31211TG4
drprTG4
hppyTG4
LerpTG4
NdgTG4
PCBTG4
RimTG4
TuspTG4
*
****
***
**

**Figure S4: Expression of TG4 lines in larval brain corresponding to hits from the overexpression screen**

Third instar larval brain imaging of *TG4* in *PMCA*, *IRSp53*, *Eph*, *Grpk2*, *if*, hppy, *msn*, *SIFaR*, *PCB*, *Pdk*, *gig* TG4 mutants driving UAS-nlsGFP (green in top row) co-stained with neuronal (Elav) and glial (Repo) nuclear markers using corresponding colocalization channels to indicate neuronal and/or glial expression (white in the bottom two rows). Scale bar = 25 μm.

**Figure S4:**

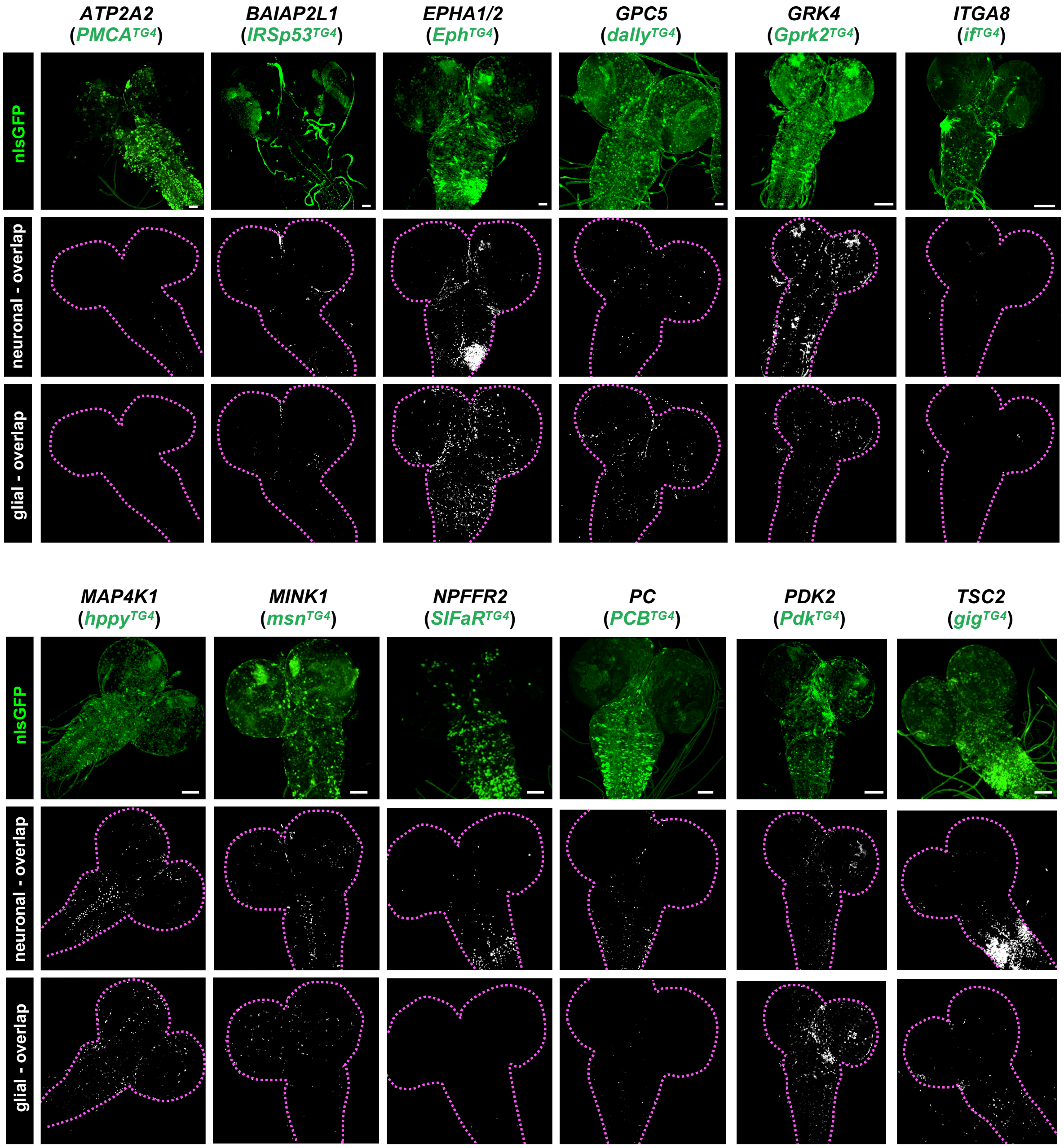

ATP2A2
(PMCA^TG4)
BAIAP2L1
(IRSp53^TG4)
EPHA1/2
(Eph^TG4)
GPC5
(dally^TG4)
GRK4
(Gprk2^TG4)
ITGA8
(if^TG4)
nlsGFP
neuronal - overlap
glial - overlap
MAP4K1
(hppy^TG4)
MINK1
(msn^TG4)
NPFFR2
(SIFaR^TG4)
PC
(PCB^TG4)
PDK2
(Pdk^TG4)
TSC2
(gig^TG4)

**Figure S5: Summary of gene and variant level statistics for hits identified from overall screen and each sub-screen**

(A) Contingency graph for variant consequences (red=loss of function, blue=gain of function, yellow=complex) grouped by variants corresponding to lethal (essential genes) or viable (non-essential genes) *TG4* mutants. Chi square ($p<0.0001$). (B-Y) Gene level constraints from gnomAD or variant level pathogenicity predictions (PolyPhen2, SIFT, CADD) for entire screen (B-G), rescue-based screen of lethality (H-M) or behavior (N-S) and overexpression of SSC related genes (T-Y) fly genes. No statistically significant differences were found using ANOVA followed by Dunn's multiple comparison test (B-J, T-Y) or t-test (K-S). (Z) Percent of SSC-DNMs present in the gnomAD database for entire screen. No statistical difference found using Chi square.

**Figure S5:**

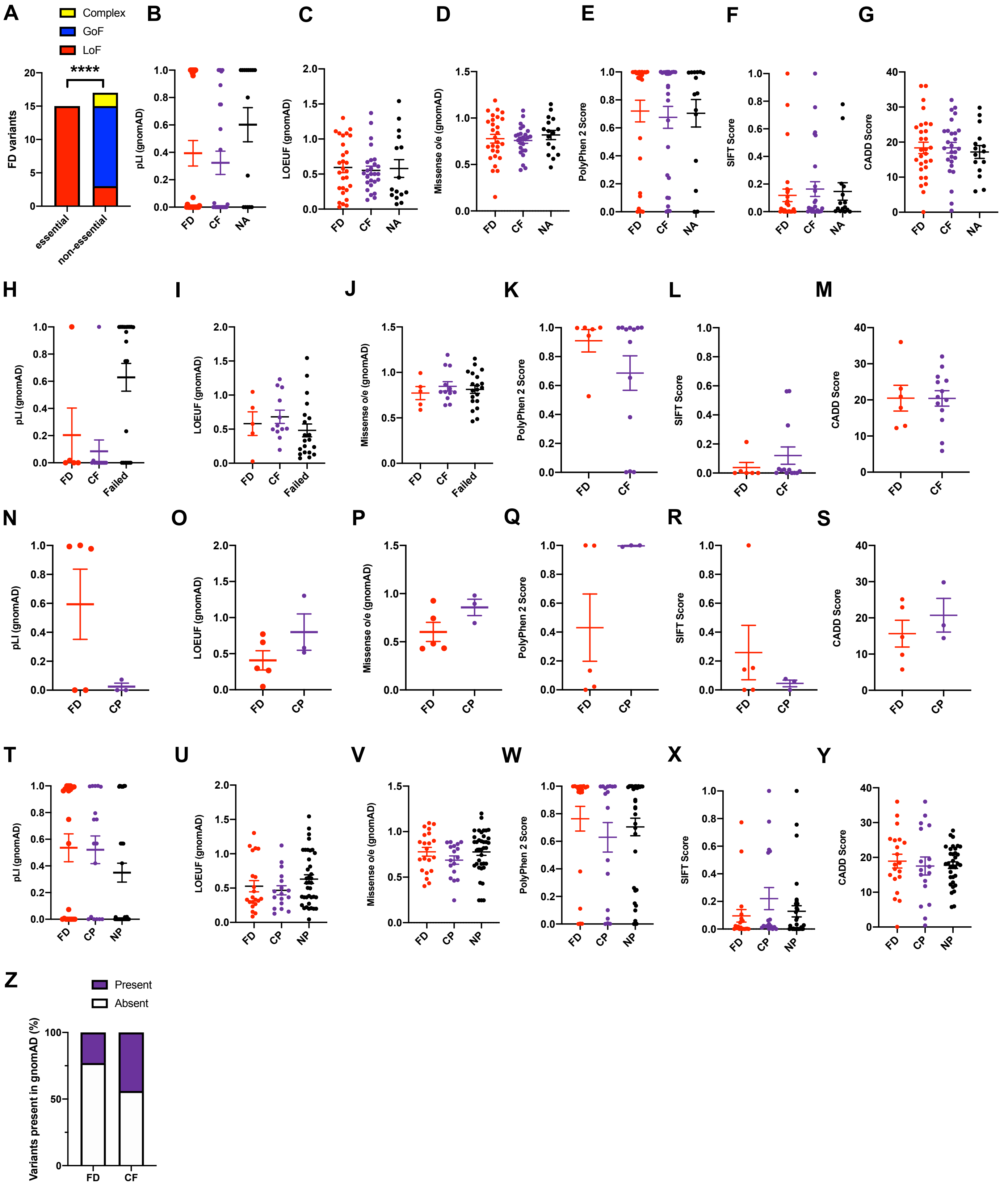
A
Complex
GoF
LoF
****
FD variants
essential
non-essential
B
pLI (gnomAD)
FD
CF
NA
C
LOEUF (gnomAD)
D
Missense o/e (gnomAD)
E
PolyPhen 2 Score
F
SIFT Score
G
CADD Score
H
I
J
K
L
M
Failed
N
O
P
Q
R
S
CP
T
U
V
W
X
Y
NP
Z
Present
Absent
Variants present in gnomAD (%)

**Figure S6: Gene Ontology (GO) of ASD candidate genes corresponding to variants with functional alterations compared to reference allele**

(A-B) Output of GO enrichment analysis for (A) ‘cellular compartment’ and (B) ‘molecular process’ based on PANTHER. A minimum of 4 genes was used as a threshold. No statistically significant GO terms corresponding to ‘biological process’ was found.

### Figure S6:

**A**

| GO cellular component complete | # | # | expected | Fold Enrichment | +/- | raw P value | FDR |
|---|---|---|---|---|---|---|---|
| peroxisome | 149 | 4 | 0.24 | 16.96 | + | 9.61E-05 | 3.20E-02 |
| -microbody | 149 | 4 | 0.24 | 16.96 | + | 9.61E-05 | 3.85E-02 |
| plasma membrane region | 1230 | 10 | 1.95 | 5.14 | + | 1.38E-05 | 9.22E-03 |
| integral compenent of plasma membrane | 1655 | 13 | 2.62 | 4.96 | + | 6.44E-07 | 6.44E-04 |
| -intrinsic compenent of plasma membrane | 1733 | 14 | 2.74 | 5.1 | + | 1.39E-07 | 2.79E-04 |
| synapse | 1304 | 9 | 2.06 | 4.36 | + | 1.45E-04 | 3.64E-02 |
| -cell junction | 2066 | 12 | 3.27 | 3.67 | + | 4.40E-05 | 2.20E-02 |
| plasma membrane bounded cell projection | 2269 | 12 | 3.59 | 3.34 | + | 1.10E-04 | 3.14E-02 |
| -cell projection | 2367 | 12 | 3.75 | 3.2 | + | 1.65E-04 | 3.67E-02 |

**B**

| GO cellular component complete | # | # | expected | Fold Enrichment | +/- | raw P value | FDR |
|---|---|---|---|---|---|---|---|
| protein kinase activity | 585 | 8 | 0.93 | 8.64 | + | 2.99E-06 | 1.78E-03 |
| -catalytic activity, acting on a protein | 2274 | 12 | 3.60 | 3.61 | + | 2.17E-05 | 5.74E-03 |
| -kinase activity | 772 | 8 | 1.22 | 6.55 | + | 2.21E-05 | 5.54E-03 |
| -transferase activity, transferring phosphorus-containing groups | 937 | 8 | 1.48 | 5.39 | + | 8.63E-05 | 1.71E-02 |
| -transferase activity | 2353 | 15 | 3.72 | 4.03 | + | 8.86E-07 | 1.05E-03 |
| -phosphotransferase activity | 691 | 8 | 1.09 | 7.32 | + | 1.00E-05 | 3.66E-03 |
| ATP binding | 1511 | 13 | 2.39 | 5.44 | + | 2.27E-07 | 1.08E-03 |
| -adenyl ribonucleotide binding | 1571 | 13 | 2.49 | 5.23 | + | 3.55E-07 | 8.45E-04 |
| -adenyl nucleotide binding | 1583 | 13 | 2.51 | 5.19 | + | 3.88E-07 | 6.15E-04 |
| -purine nucleotide binding | 1940 | 13 | 3.07 | 4.23 | + | 3.83E-06 | 1.66E-03 |
| -nucleotide binding | 2180 | 14 | 3.45 | 4.06 | + | 2.26E-06 | 2.15E-03 |
| -small molecule binding | 2591 | 15 | 4.10 | 3.66 | + | 3.02E-06 | 1.60E-03 |
| -nucleoside phosphate binding | 2181 | 14 | 3.45 | 4.06 | + | 2.27E-06 | 1.80E-03 |
| -purine ribonucleotide binding | 1926 | 13 | 3.05 | 4.26 | + | 3.54E-06 | 1.68E-03 |
| -ribonucleotide binding | 1943 | 13 | 3.08 | 4.23 | + | 3.90E-06 | 1.55E-03 |
| -carbohydrate derivative binding | 2287 | 13 | 3.62 | 3.59 | + | 2.31E-05 | 5.49E-03 |
| -purine ribonucleotide triphosphate binding | 1859 | 13 | 2.94 | 4.42 | + | 2.39E-06 | 1.62E-03 |
| -anion binding | 2892 | 15 | 4.58 | 3.28 | + | 1.19E-05 | 4.04E-03 |
| -ion binding | 6393 | 21 | 10.12 | 2.08 | + | 1.01E-04 | 1.93E-02 |

**Figure S7: Expression analysis of additional TG4 lines related to SSC-DNM genes in the adult brain**

Adult brain imaging of TG4 driving UAS-nlsGFP (green in top row) co-stained with neuronal (Elav) and glial (Repo) nuclear markers using corresponding colocalization channels to indicate neuronal and/or glial expression (white in bottom two rows). Scale bar = 25 μm.

**Figure S7:**

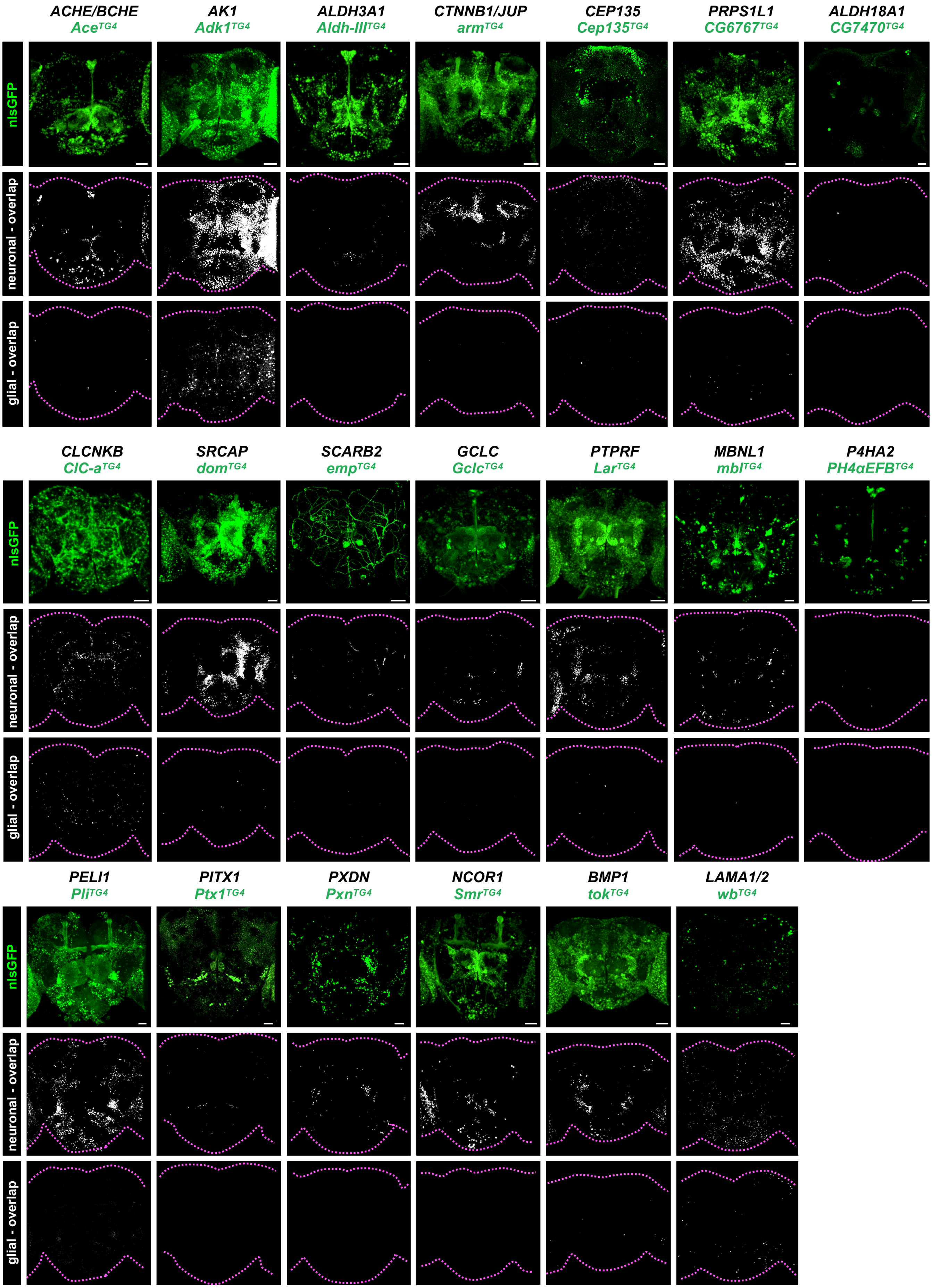
ACHE/BCHE
Ace[TG4]
AK1
Adk1[TG4]
ALDH3A1
Aldh-III[TG4]
CTNNB1/JUP
arm[TG4]
CEP135
Cep135[TG4]
PRPS1L1
CG6767[TG4]
ALDH18A1
CG7470[TG4]
nlsGFP
neuronal - overlap
glial - overlap
CLCNKB
ClC-a[TG4]
SRCAP
dom[TG4]
SCARB2
emp[TG4]
GCLC
Gclc[TG4]
PTPRF
Lar[TG4]
MBNL1
mbl[TG4]
P4HA2
PH4αEFB[TG4]
PELI1
Pli[TG4]
PITX1
Ptx1[TG4]
PXDN
Pxn[TG4]
NCOR1
Smr[TG4]
BMP1
tok[TG4]
LAMA1/2
wb[TG4]

**Figure S8: Expression analysis of additional TG4 lines related to SSC-DNM genes in the larval brain**

Third instar larval brain imaging of TG4 driving UAS-nlsGFP (green in top row) co-stained with neuronal (Elav) and glial (Repo) nuclear markers using corresponding colocalization channels to indicate neuronal and/or glial expression (white in bottom two rows). Scale bar = 25 μm.

### Figure S8:

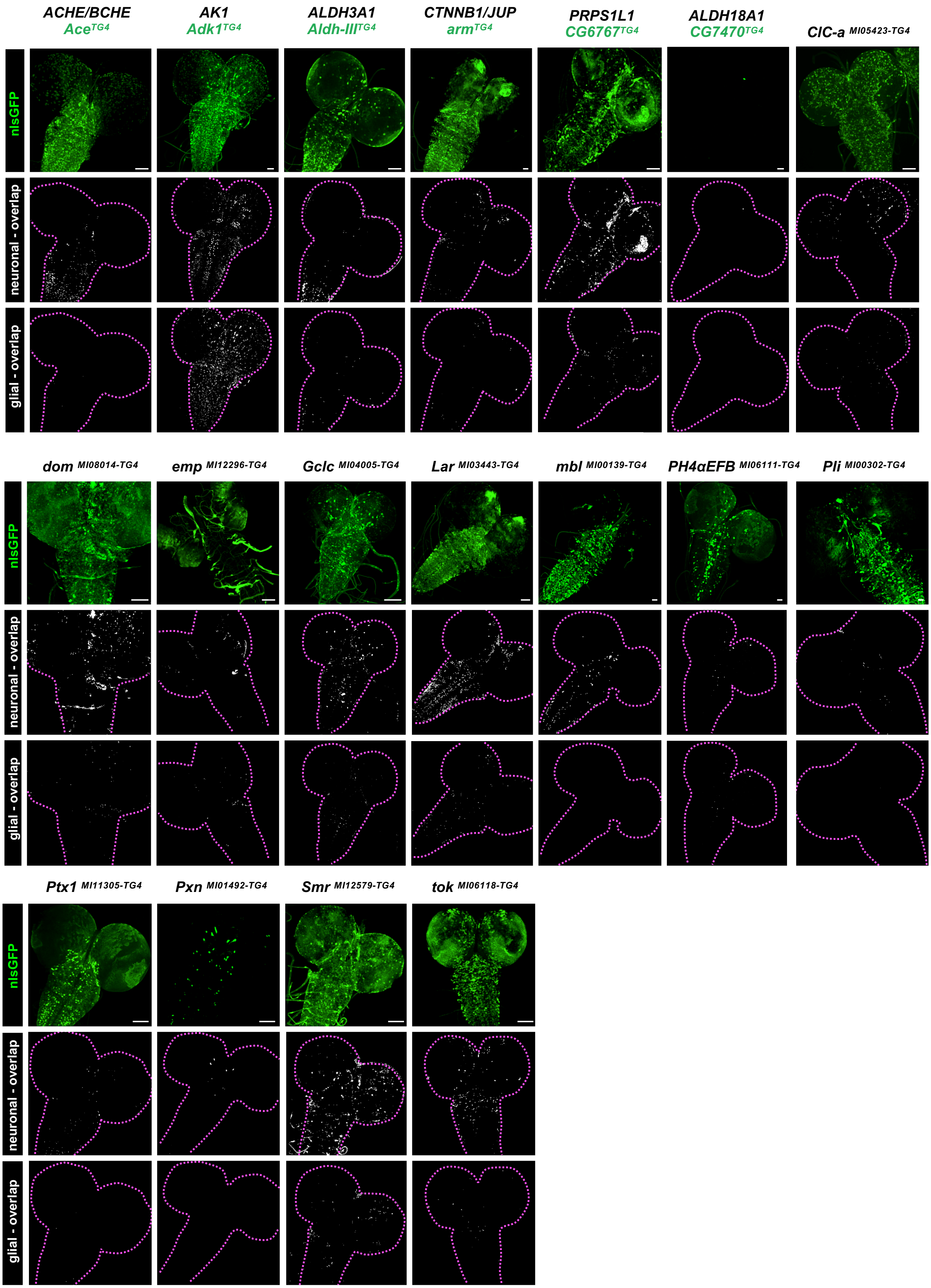
ACHE/BCHE
Ace TG4
AK1
Adk1 TG4
ALDH3A1
Aldh-III TG4
CTNNB1/JUP
arm TG4
PRPS1L1
CG6767 TG4
ALDH18A1
CG7470 TG4
ClC-a MI05423-TG4
nlsGFP
neuronal - overlap
glial - overlap
dom MI08014-TG4
emp MI12296-TG4
Gclc MI04005-TG4
Lar MI03443-TG4
mbl MI00139-TG4
PH4αEFB MI06111-TG4
Pli MI00302-TG4
Ptx1 MI11305-TG4
Pxn MI01492-TG4
Smr MI12579-TG4
tok MI06118-TG4

**Figure S9: Additional data related to *GLRA2* variant functional assessments**

(A) Western blot of 5-day old fly heads expressing *GLRA2* cDNA constructs tagged with a C terminal 3xHA tag expressed using a pan-neuronal driver (*nSyb-GAL4*). (B) Viability flies expressing *GLRA2* reference or p.T296M at 29°C. (C-F) Quantification of depolarization and "*ON*"-transient amplitudes from electroretinogram (ERG) recordings of flies expressing GLRA2 reference or variants pan-neuronally (*nSyb-GAL4*) or only in photoreceptors (*Rh1-GAL4*). (G) GLRA1 crystal structure indicating location of critical homologous GLRA2 amino acid residues relevant to amino acids affected by three patient variants (p.N136, p.T296, p.R252) experimentally tested here. ***p<0.001, ns (not significant).

### Figure S9:

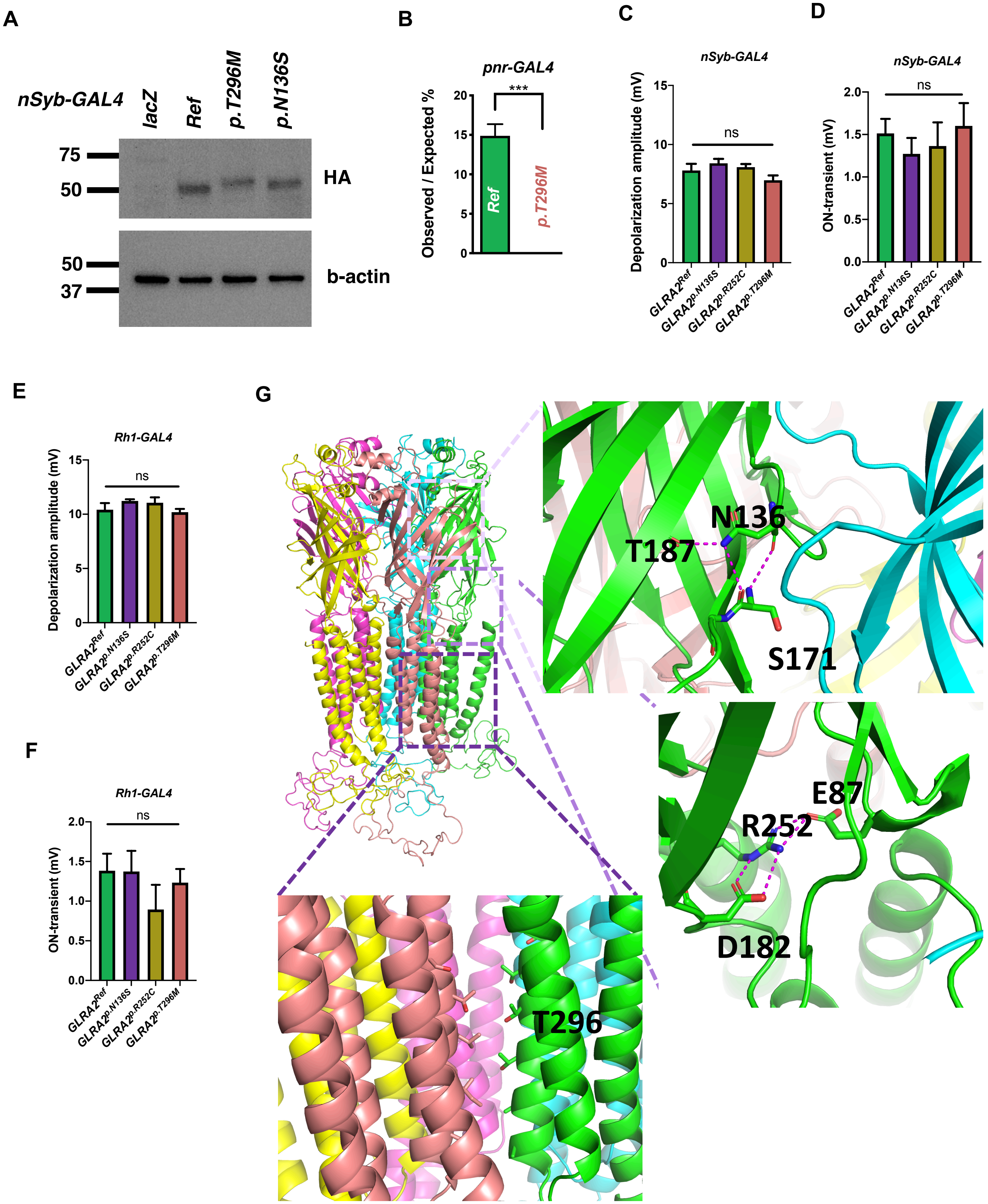
A
nSyb-GAL4
lacZ
Ref
p.T296M
p.N136S
75
50
HA
50
37
b-actin
B
pnr-GAL4
***
Observed / Expected %
20
15
10
5
0
Ref
p.T296M
C
nSyb-GAL4
Depolarization amplitude (mV)
ns
GLRA2Ref
GLRA2p.N136S
GLRA2p.R252C
GLRA2p.T296M
D
nSyb-GAL4
ON-transient (mV)
ns
E
Rh1-GAL4
Depolarization amplitude (mV)
ns
F
Rh1-GAL4
ON-transient (mV)
ns
G
N136
T187
S171
E87
R252
D182
T296
