## Extended Case Histories for "*Drosophila* functional screening of *de novo* variants in autism uncovers deleterious variants and facilitates discovery of rare neurodevelopmental diseases"

### SI Case Histories

**GLRA2 subject case histories**

Subject 1 is an 8-year-old female with global developmental and cognitive delay. Pregnancy was naturally conceived and uncomplicated, other than decreased fetal movements noted by the mother. She was delivered at term (39 weeks gestational age) via c/section due to breech presentation. Birth weight was 3,600 grams. Neonatal period was uneventful. There were no feeding difficulties and her growth remained within the normal limits. She was delayed with all her milestones but most significantly for speech (walked at 18 months, first words at 24 months and combined words to sentences at 4-5 years of age, scribbled with a crayon at 3.5 years). She was diagnosed with mixed expressive-receptive speech delay and received speech therapy, occupational therapy and physical therapy interventions. In school she exhibits learning problems, inattention and is below her grade level. She has a modified curriculum and is receiving resources in reading and math. There is no history of developmental regression or seizures. The medical history is otherwise significant for nystagmus that was first noted in infancy and improved with age, as well as myopia and astigmatism requiring corrective glasses. Family ethnicity is Hispanic and the family history was non-contributory. The patient had a normal brain MRI (magnetic resonance imaging) at 6 months of age. EEG (electroencephalography) at 4 years of age showed a slow and poorly formed background, indicative of mild encephalopathy, but did not detect epileptiform activity. Genetic testing included: mitochondrial DNA sequencing which detected a pathogenic variant m.13042 G>A though at heteroplasmy level of 1.9%, however this was felt unlikely to explain the phenotype. CMA (chromosomal microarray) was negative. Trio whole exome sequencing (WES) detected a *de novo*, heterozygous variant of unknown clinical significance in *GLRA2*, c.887C>T, p.Thr296Met (NC_000023.10: g.14627284C>T). This variant is absent in gnomAD. More recent clinical reanalysis of exome data did not detect any other candidates that may explain the phenotype.

Subject 2 is a 6-year-old female with epilepsy, developmental delay (DD), mild intellectual disability (ID) and autism spectrum disorder (ASD). Pregnancy was uncomplicated and she was delivered at term (41 weeks gestational age) via vaginal delivery with vacuum extraction. The neonatal period was uneventful. At the age of 6 months, she developed a severe epileptic encephalopathy with myoclonic seizures. Seizure control was achieved with medications, and she has been seizure-free without medications since the age of about two years old. Delayed psychomotor development was noted, most significantly for her speech with a mixed expressive-receptive speech delay (non-verbal). Her ability to concentrate is poor and she displays mood swings. The medical history is otherwise significant for nystagmus that was first noted in infancy (6 weeks old) and improved with age, and sleep disturbance. She has mild microcephaly [< 1st centile: -2.84 standard deviation (SD)] and mild bilateral cutaneous 3rd-4th syndactyly, with no other congenital anomalies. Family ethnicity is European (German/Italian) and the family history is significant for a maternal aunt that had epilepsy in adulthood but her cognitive development was normal. Brain MRI showed delayed myelination at 7 months old and a small arachnoid cyst. EEG was abnormal for bilateral synchronized, sometimes high amplitude spike/polyspike-waves-complexes, and bitemporo-occipital hints for severe functional defects with epileptic potentials. Chromosomal analysis, Angelman syndrome methylation study, epilepsy next generation sequencing (NGS) gene panel and *MECP2* sequencing were negative. Trio WES detected a *de novo*, heterozygous variant of unknown clinical significance in *GLRA2*, c.887C>T, p.Thr296Met (NC_000023.10: g.14627284C>T). This variant is absent in gnomAD.

Subject 3 is a 5 year 6 months old female with DD, microcephaly, abnormal eye movements and ataxic gait. Pregnancy was uncomplicated and she was born at term via c/section. Abnormal eye movements were noticed two weeks after birth, during hospitalization due to a lower respiratory tract infection. At the age of 6 months, clinical examination revealed mildly delayed developmental milestones and erratic conjugate eye movements akin to opsoclonus. At age 4 years OFC (occipitofrontal circumference) was 43 cm (< 1st centile: -4.28 SD) and ophthalmological evaluation revealed alternating exotropia, for which patching therapy was initiated. Language was limited to a few words and neuropsychological evaluation documented moderate developmental delay (Bayley-III). The patient could walk unsupported with ataxic gait. At age 5 years 6 months, erratic eye movements were considerably reduced and she could walk independently but her expressive language was still limited to a few words, with delayed receptive speech and nonverbal communicative skills. Family ethnicity is European and the family history is unremarkable. Brain MRI at 6 months of age showed mild cortical atrophy with thinning of the corpus callosum. EEG, while awake and asleep, laboratory and metabolic investigations were unremarkable. Array-CGH (comparative genomic hybridization) highlighted a maternally inherited 3q25.32 duplication (chr3:157746089-158324659, hg19) that was interpreted as likely benign. Trio WES detected a *de novo* heterozygous variant in *GLRA2*, c.887C>T, p.Thr296Met (NC_000023.10: g.14627284C>T). This variant is absent in gnomAD. In addition, it detected a *de novo* variant in *CACNA1B*, c.5381C>T, p.Thr1794Met (NC_000009.11:g.141000212C>T), which is a variant of unknown significance in a gene that is linked to an autosomal recessive condition (Neurodevelopmental disorder with seizures and nonepileptic hyperkinetic movements, MIM #618497). Failure to identify a second allele in this gene reduces the likelihood that this variant is responsible for this patient's phenotype.

Subject 4 was a female infant with seizures and severe developmental delay who passed away at 7 months of age secondary to complications of COVID-19 infection. Pregnancy was uneventful and she was born at term (40 weeks gestational age). She was noted to have focal seizures at 2-3 weeks of age, and was diagnosed with infantile spasms when she was 5 months old. At 6 months of age she was not reaching for objects, not sitting up and only making high-pitched sounds. She had borderline microcephaly with dysmorphic features including midface retrusion, apparent hypotelorism, deep set eyes, thick eyebrows, downturned corners of the mouth, and wide-spaced nipples. Family ethnicity is Hispanic and the family history was unremarkable. She had normal plasma and CSF (cerebrospinal fluid) lactate, pipecolic acid and piperideine-6-carboxylate, ammonia, urine organic acids, plasma amino acids, acylcarnitine profile, and CSF amino acids. An Epilepsy gene panel was non-diagnostic. Trio WES detected a *de novo* heterozygous variant in *GLRA2*, c.887C>T, p.Thr296Met (NC_000023.10: g.14627284C>T). This variant is absent in gnomAD.

Subject 5 is a 6 years and 7 months old female with a history of infantile spasms, epilepsy and intellectual disability. She was born at term and first presented with infantile spasms at 3 months of age. This evolved to atonic and tonic-clonic seizures as she grew up. She was delayed with all milestones (walked at 4.5 years old and remains non-verbal). She had nystagmus that improved with age and strabismus. The medical history is otherwise significant for hyperactivity, inattention and sleep disturbance. Her ethnicity is African (Senegal). Brain MRI at 3 years of age showed cortical and white matter atrophy, including vermian atrophy. EEG showed hypsarrythmia at onset and she had normal interictal EEG afterwards. She had normal SNP (single nucleotide polymorphism) array, negative targeted epilepsy panel and negative metabolic lab results. Trio WES identified a *de novo* heterozygous variant in *GLRA2*, c.140T>C, p.Phe47Ser (NC_000023.10: g.14550432T>C). This variant is absent in gnomAD.

Subject 6 is an 11-month-old male with hypotonia, DD and dysmorphic craniofacial features. Pregnancy was uncomplicated, he was delivered at term (38 and 3/7 weeks gestational age) and the neonatal period was uneventful. Soon after birth dysmorphic features were noted, including an elongated face, high anterior hairline, epicanthal folds, downslanting palpebral fissures and a bulbous nose. Growth remains within the normal limits. His medical history is otherwise significant for obstructive sleep apnea and strabismus. Family ethnicity is European (Dutch) and the family history is significant for the maternal grandfather who has not further specified unexplained neurological complaints, and which could not be further investigated. Investigations for metabolic disorders, Fragile X syndrome and a SNP-array were normal. Trio WES identified a rare variant in *GLRA2*, c.754C>T, p.Arg252Cys (NC_000023.10: g.14627151C>T), which was inherited from mother. No other possible disease explaining variant was identified. The mother displayed skewed X chromosome inactivation (82% on two measurements). The variant was absent in the maternal uncle and the maternal grandmother, but was inherited from the maternal grandfather, who was not available for clinical investigations. His level of functioning remains unknown. This variant is present in one heterozygous female in gnomAD.

Subject 7 is a 7-year-old male with epilepsy, DD with regression, and ASD. Pregnancy was uncomplicated. He was born full term via uncomplicated delivery, and his early development was as expected. He was speaking in sentences at 2.5 years old when he started having generalized tonic-clonic seizures. He developed staring spells, ataxia, and an increased frequency of myoclonic jerks, which around the age of 6 years old were occurring 20 times per day on average, with 5-6 atonic seizures per day each lasting less than 30 seconds. Following seizure onset he experienced developmental regression. At 3 years of age he was diagnosed with ASD. At 6 years of age his vocabulary was about 20 words, with gains in development lost following significant seizures. His ethnicity is European, and the family history is significant for a younger brother with ASD, although he has not presented with seizures. Neither mother nor father have a history of seizures or delays. At age three, EEG depicted generalized slowing and generalized epileptiform discharges associated with myoclonic jerks. MRI showed minimal increased T2 signal intensity on the occipital lobes that was thought to be within normal limits. Genetics testing for Fragile X syndrome, Prader-Willi/Angelman syndromes, and congenital disorders of glycosylation were normal. Additional tests, including plasma amino acids, lysosomal enzymes, and cerebral creatine deficiency were also normal. Microarray reported a maternally inherited 1p33 deletion of unknown significance (48,688,391-49,922,153). The patient was enrolled to The Manton Center for Orphan Disease Gene Discovery Core protocol. Trio WES discovered a maternally inherited variant in *GLRA2*, c.862G>A, p. Ala288Thr (NC_000023.10: g.14627259G>A). This variant is absent in gnomAD.

Subject 8 is a 35-year-old male with a history of DD, learning disabilities and ASD. Pregnancy was uncomplicated and he was born at term (40 weeks gestational age). Since early childhood he showed slow movement and difficulties in motor coordination. He walked and said his first words at 24 months, and first sentences at age 3 years of age. In school learning disabilities were noted, including difficulties in writing, reading, praxias, temporal orientation, calculation, drawing, and visuo-spatial organization. He graduated high school and continued to higher education, though he did not complete a degree. Neuropsychiatric assessment in adulthood was consistent with ASD and social and cognitive deficits. There is no history of seizures. The medical history is otherwise significant for environmental allergies, myopia and astigmatism. The ethnicity is European, and the family history is unremarkable, except for a maternal grandmother with Alzheimer's dementia. The patient had a normal brain MRI at 29 years old. Trio WES identified a maternally inherited variant in *GLRA2*, c.1186C>A, p. Pro396Thr (NC_000023.10: g.14748434C>A). This variant is present in 3 heterozygous females and 1 hemizygous male in gnomAD.
