## Supplemental Figures for "*Drosophila* functional screening of *de novo* variants in autism uncovers deleterious variants and facilitates discovery of rare neurodevelopmental diseases"

#### Supplementary Figures

**Supplementary Fig. 1: Lethality of *TG4* mutants and its implication on essentiality of *Drosophila* genes**

**a**, Assessment of lethal *TG4* lines by complementation tests with corresponding deficiency (Df) lines. Lethality of 17 TG4 lines were due to a second site mutations, potentially present in the original MiMIC line or introduced during RMCE. **b**, Table summarizing the tools generated in this study in respect to essentiality of fly genes. **c-e**, Gene level statistics for human homologs of essential and non-essential fly genes. No significant differences were found with t-test analysis.

#### Supplementary Fig. 1:

**a**

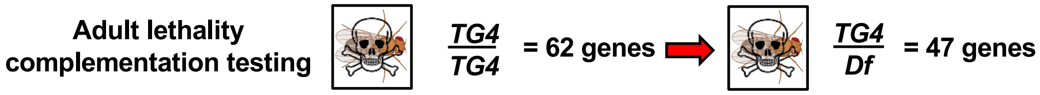


**b**

| | *D. mel* LoF *TG4* lines (Corresponding # *H. sap* genes) | *H. sap* transgenics Ref *UAS-cDNA* | Var *UAS-cDNA* |
|---|---|---|---|
| **essential genes** | 47 (60) | 45 | 49 |
| **non-essential genes** | 62 (68) | 52 | 34 |
| **failed MiMIC conversions** | 13 (15) | 9 | 7 |
| **Total fly lines made** | 109* (128) | 106 | 90 |

*36 lines are novel since publication of Lee *et al.*, 2018

**c** **d** **e**

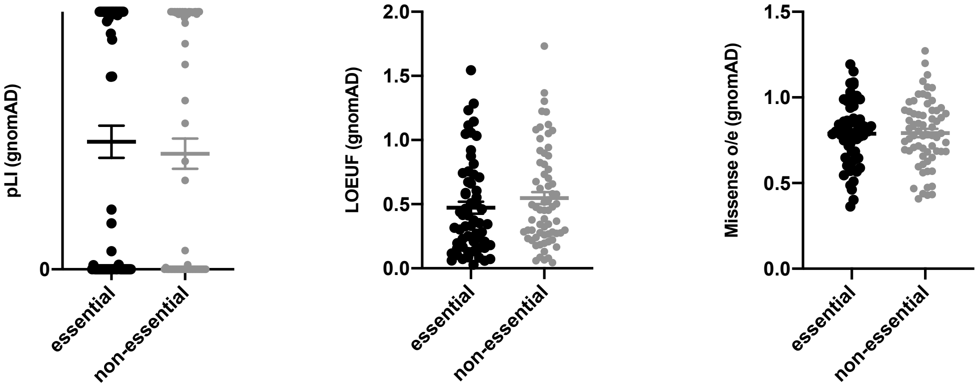

**Supplementary Fig. 2: Expression of TG4 lines corresponding to hits from the humanization screen corresponding to essential fly genes in larval brains**

Third instar larval brain imaging of *TG4* in *Abl*, *Cat*, *CG31637*, *ctrip* and *Trpm* driving UAS-nlsGFP (green in top row) co-stained with neuronal (Elav) and glial (Repo) nuclear markers displayed using corresponding colocalization channels to indicate neuronal and/or glial expression (white in the bottom two rows). Scale bar = 25 µm.

### Supplementary Fig. 2:

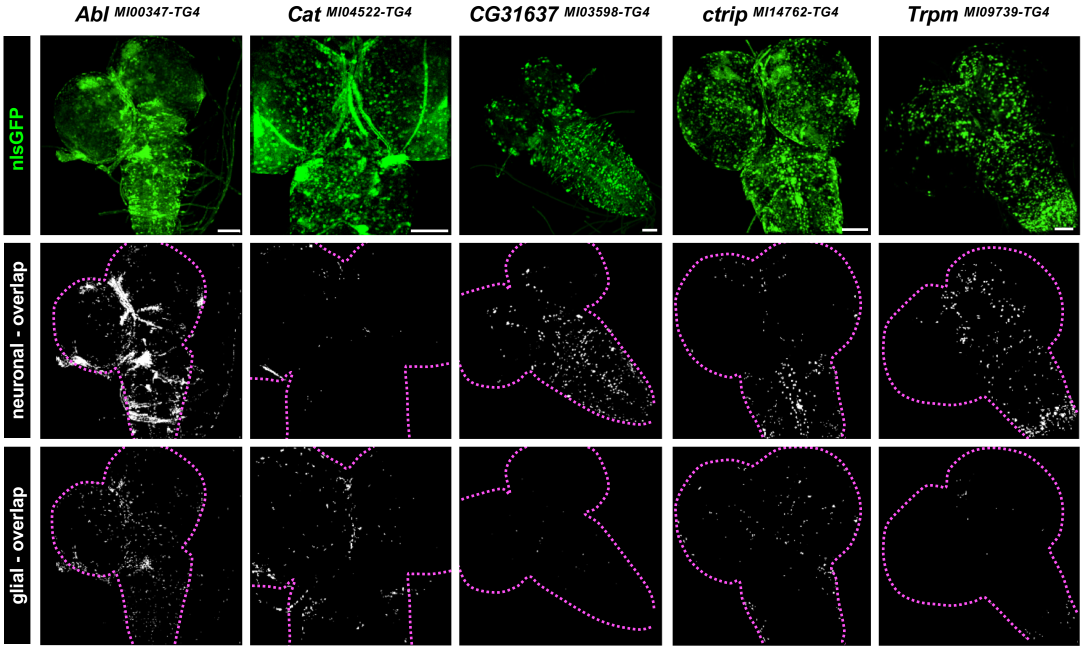

**Supplementary Fig. 3: Expression of TG4 lines corresponding to hits from the overexpression screen corresponding to essential fly genes in larval brains**

Third instar larval brain imaging of *TG4* in *gig*, *Grpk2*, *if*, *msn*, *Pdk*, *PMCA* TG4 mutants driving UAS-nlsGFP (green in top row) co-stained with neuronal (Elav) and glial (Repo) nuclear markers using corresponding colocalization channels to indicate neuronal and/or glial expression (white in the bottom two rows). Scale bar = 25 μm.

### Supplementary Fig. 3:

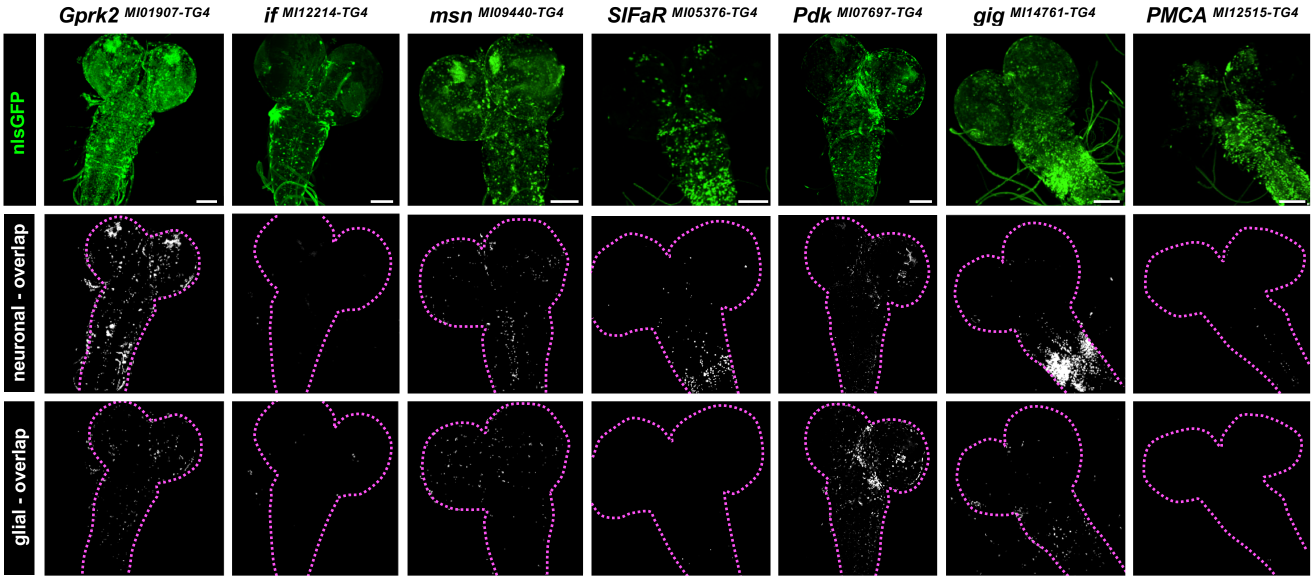

**Supplementary Fig. 4: Additional behavior data for viable *TG4* fly mutants**

**a-d**, For non-essential *TG4* mutants that we were unable to humanized due to technical reasons, we also assessed the number of frames male flies spent performing single-wing extensions (courtship), copulating, moving within the chamber, or grooming during a test period. The red line represents the average number of frames a *Canton-S* male spends doing the same activity. *p<0.05, **p<0.01, ***p<0.001, ****p<0.0001. **e**, Third instar larval brain imaging of *TG4* in *5-HT1B*, *GluCI*$\alpha$, *Rab3-GEF*, and *Usp30* TG4 mutants driving UAS-nlsGFP (green in top row) co-stained with neuronal (Elav) and glial (Repo) nuclear markers using corresponding colocalization channels to indicate neuronal and/or glial expression (white in the bottom two rows). Scale bar = 25 µm.

### Supplementary Fig. 4:

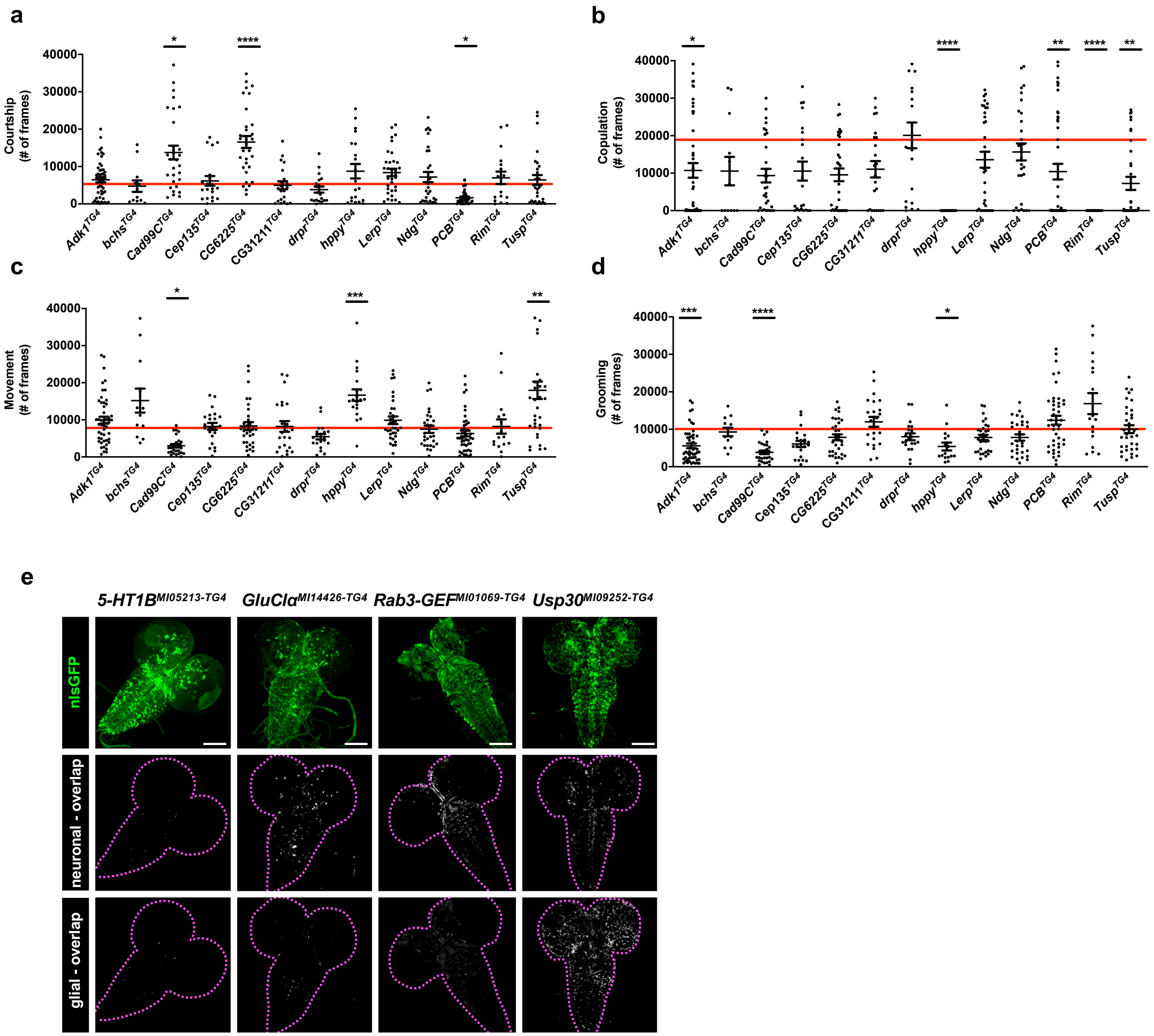

**Supplementary Fig. 5: Expression of TG4 lines corresponding to hits from the humanization screen corresponding to non-essential fly genes in larval brains**

Third instar larval brain imaging of *TG4* in *dally*, *Eph*, *hppy*, *IRSp53* driving UAS-nlsGFP (green in top row) co-stained with neuronal (Elav) and glial (Repo) nuclear markers using corresponding colocalization channels to indicate neuronal and/or glial expression (white in the bottom two rows). Scale bar = 25 μm.

#### Supplementary Fig. 5:

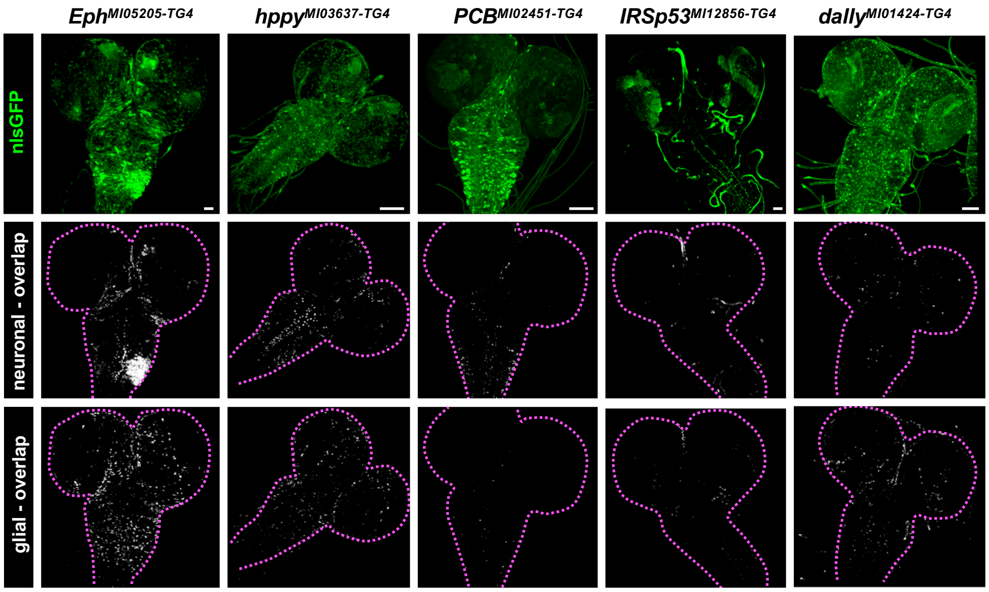

**Supplementary Fig. 6: Summary of gene and variant level statistics for hits identified from overall screen and each sub-screen**

**a**, Contingency graph for variant consequences (red=loss of function, blue=gain of function, yellow=complex) grouped by variants corresponding to lethal (essential genes) or viable (non-essential genes) *TG4* mutants. Chi square ($p<0.0001$). **b-ae**, Gene level constraints from gnomAD or variant level pathogenicity predictions (PolyPhen2, SIFT, CADD) for entire screen (b-g), rescue-based screen of lethality (h-m) or behavior (t-y) and overexpression of SSC related genes for essential (m-s) and viable (z-ae) fly genes. No statistically significant differences were found using ANOVA followed by Dunn's multiple comparison test (b-s, z-ae) or t-test (t-y). **af**, Percent of SSC-DNMs present in the gnomAD database for entire screen. No statistical difference found using Chi square.

#### Supplementary Fig. 6:

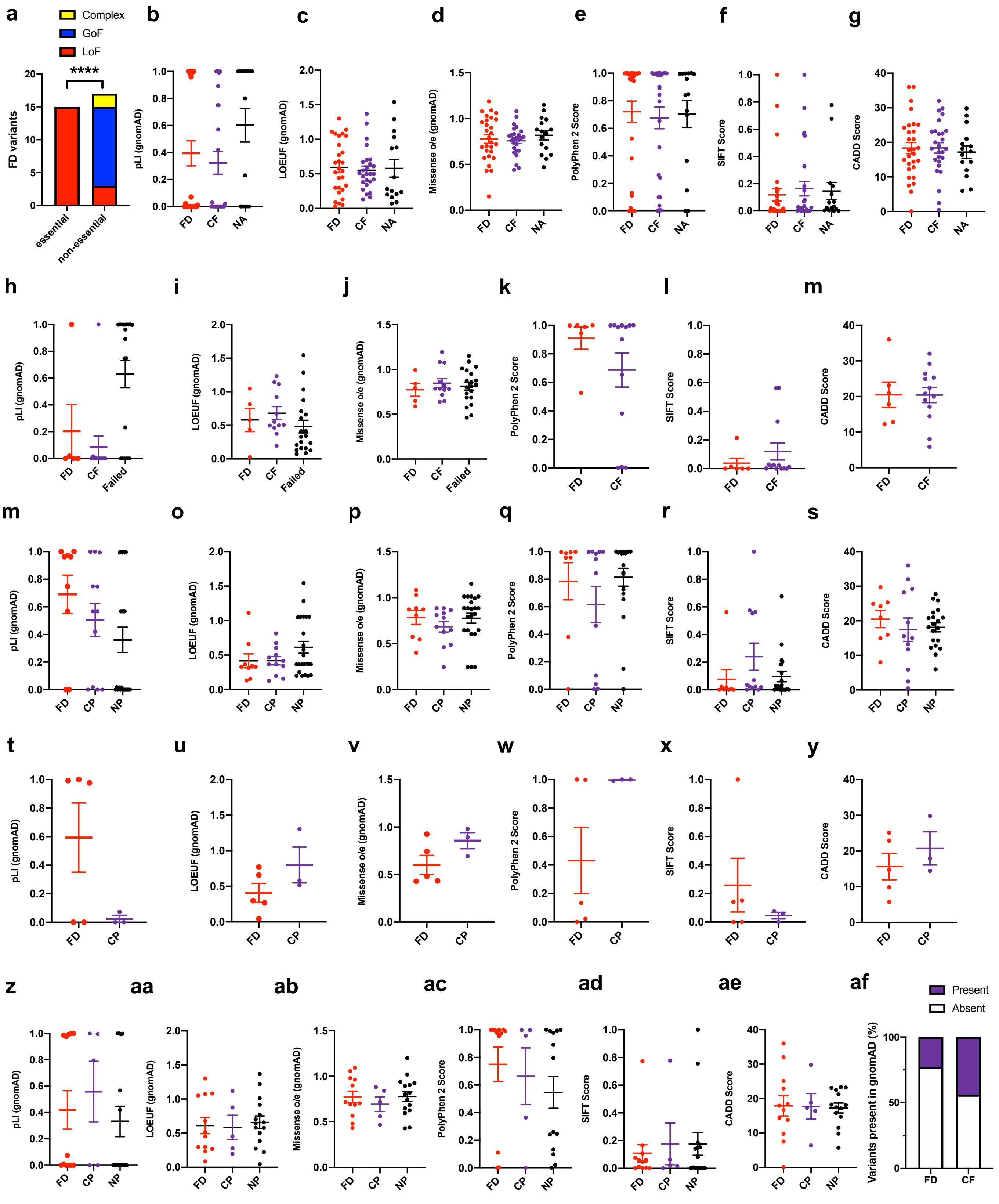

**Supplementary Fig. 7: Gene Ontology (GO) of ASD candidate genes corresponding to variants with functional alterations compared to reference allele**

**a-b**, Output of GO enrichment analysis for (**a**) ‘cellular compartment’ and (**b**) ‘molecular process’ based on PANTHER. A minimum of 4 genes was used as a threshold. No statistically significant GO terms corresponding to ‘biological process’ was found.

#### Supplementary Fig. 7:

**a**

| GO cellular component complete | # | # | expected | Fold Enrichment | +/- | raw P value | FDR |
|---|---|---|---|---|---|---|---|
| peroxisome | 149 | 4 | .24 | 16.96 | + | 9.61E-05 | 3.20E-02 |
| ↳microbody | 149 | 4 | .24 | 16.96 | + | 9.61E-05 | 3.85E-02 |
| plasma membrane region | 1230 | 10 | 1.95 | 5.14 | + | 1.38E-05 | 9.22E-03 |
| integral component of plasma membrane | 1655 | 13 | 2.62 | 4.96 | + | 6.44E-07 | 6.44E-04 |
| ↳intrinsic component of plasma membrane | 1733 | 14 | 2.74 | 5.10 | + | 1.39E-07 | 2.79E-04 |
| synapse | 1304 | 9 | 2.06 | 4.36 | + | 1.45E-04 | 3.64E-02 |
| ↳cell junction | 2066 | 12 | 3.27 | 3.67 | + | 4.40E-05 | 2.20E-02 |
| plasma membrane bounded cell projection | 2269 | 12 | 3.59 | 3.34 | + | 1.10E-04 | 3.14E-02 |
| ↳cell projection | 2367 | 12 | 3.75 | 3.20 | + | 1.65E-04 | 3.67E-02 |

**b**

| GO molecular function complete | # | # | expected | Fold Enrichment | +/- | raw P value | FDR |
|---|---|---|---|---|---|---|---|
| protein kinase activity | 585 | 8 | .93 | 8.64 | + | 2.99E-06 | 1.78E-03 |
| ↳catalytic activity, acting on a protein | 2274 | 13 | 3.60 | 3.61 | + | 2.17E-05 | 5.74E-03 |
| ↳kinase activity | 772 | 8 | 1.22 | 6.55 | + | 2.21E-05 | 5.54E-03 |
| ↳transferase activity, transferring phosphorus-containing groups | 937 | 8 | 1.48 | 5.39 | + | 8.63E-05 | 1.71E-02 |
| ↳transferase activity | 2353 | 15 | 3.72 | 4.03 | + | 8.86E-07 | 1.05E-03 |
| ↳phosphotransferase activity, alcohol group as acceptor | 691 | 8 | 1.09 | 7.32 | + | 1.00E-05 | 3.66E-03 |
| ATP binding | 1511 | 13 | 2.39 | 5.44 | + | 2.27E-07 | 1.08E-03 |
| ↳adenyl ribonucleotide binding | 1571 | 13 | 2.49 | 5.23 | + | 3.55E-07 | 8.45E-04 |
| ↳adenyl nucleotide binding | 1583 | 13 | 2.51 | 5.19 | + | 3.88E-07 | 6.15E-04 |
| ↳purine nucleotide binding | 1940 | 13 | 3.07 | 4.23 | + | 3.83E-06 | 1.66E-03 |
| ↳nucleotide binding | 2180 | 14 | 3.45 | 4.06 | + | 2.26E-06 | 2.15E-03 |
| ↳small molecule binding | 2591 | 15 | 4.10 | 3.66 | + | 3.02E-06 | 1.60E-03 |
| ↳nucleoside phosphate binding | 2181 | 14 | 3.45 | 4.06 | + | 2.27E-06 | 1.80E-03 |
| ↳purine ribonucleotide binding | 1926 | 13 | 3.05 | 4.26 | + | 3.54E-06 | 1.68E-03 |
| ↳ribonucleotide binding | 1943 | 13 | 3.08 | 4.23 | + | 3.90E-06 | 1.55E-03 |
| ↳carbohydrate derivative binding | 2287 | 13 | 3.62 | 3.59 | + | 2.31E-05 | 5.49E-03 |
| ↳purine ribonucleoside triphosphate binding | 1859 | 13 | 2.94 | 4.42 | + | 2.39E-06 | 1.62E-03 |
| ↳anion binding | 2892 | 15 | 4.58 | 3.28 | + | 1.19E-05 | 4.04E-03 |
| ↳ion binding | 6393 | 21 | 10.12 | 2.08 | + | 1.01E-04 | 1.93E-02 |

**Supplementary Fig. 8: Expression analysis of additional TG4 lines related to SSC-DNM genes in the adult brain**

Adult brain imaging of TG4 driving UAS-nlsGFP (green in top row) co-stained with neuronal (Elav) and glial (Repo) nuclear markers using corresponding colocalization channels to indicate neuronal and/or glial expression (white in bottom two rows). Scale bar = 25 μm.

#### Supplementary Fig. 8:

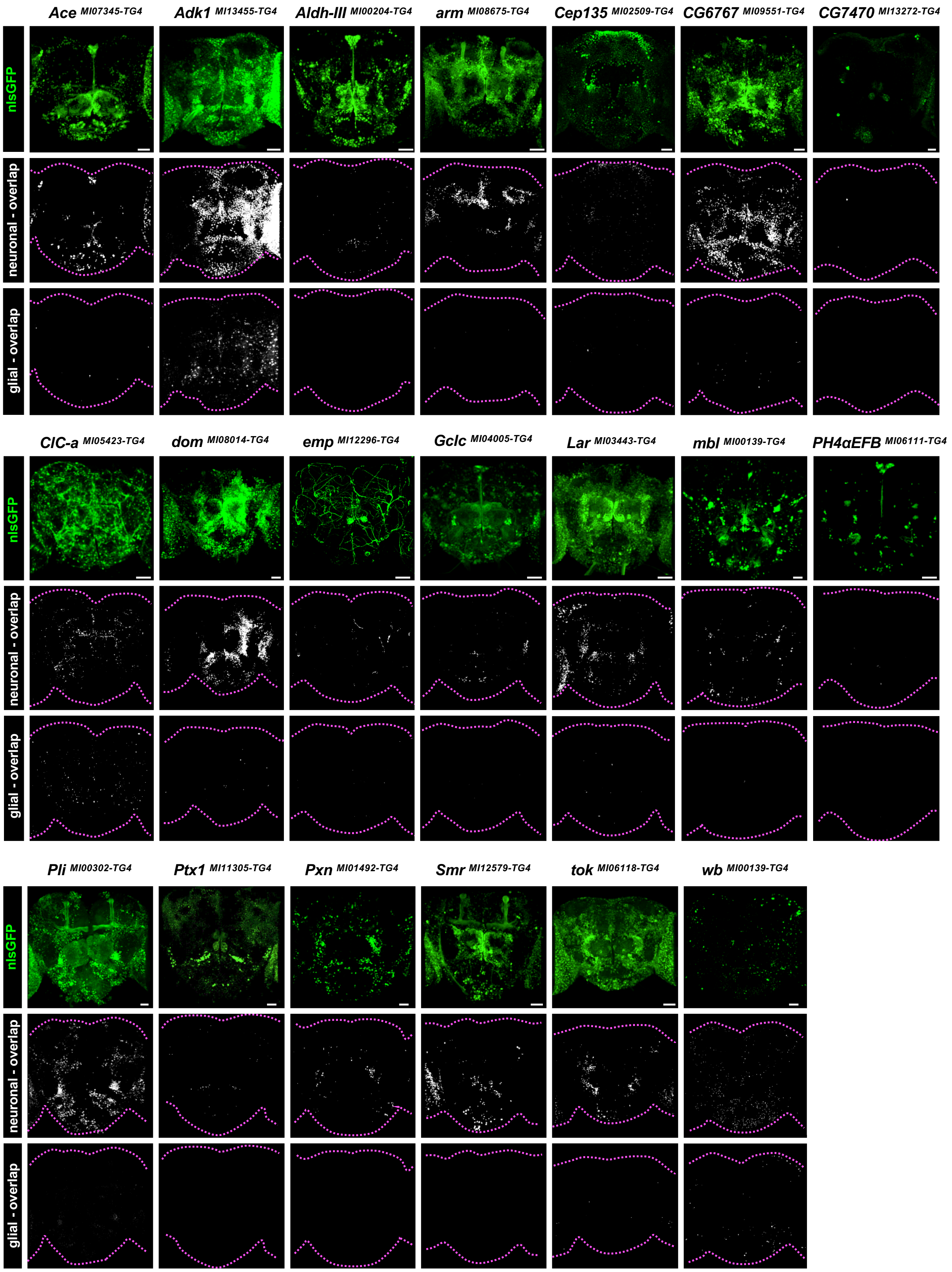

**Supplementary Fig. 9: Expression analysis of additional TG4 lines related to SSC-DNM genes in the larval brain**

Third instar larval brain imaging of TG4 driving UAS-nlsGFP (green in top row) co-stained with neuronal (Elav) and glial (Repo) nuclear markers using corresponding colocalization channels to indicate neuronal and/or glial expression (white in bottom two rows). Scale bar = 25 μm.

#### Supplementary Fig. 9:

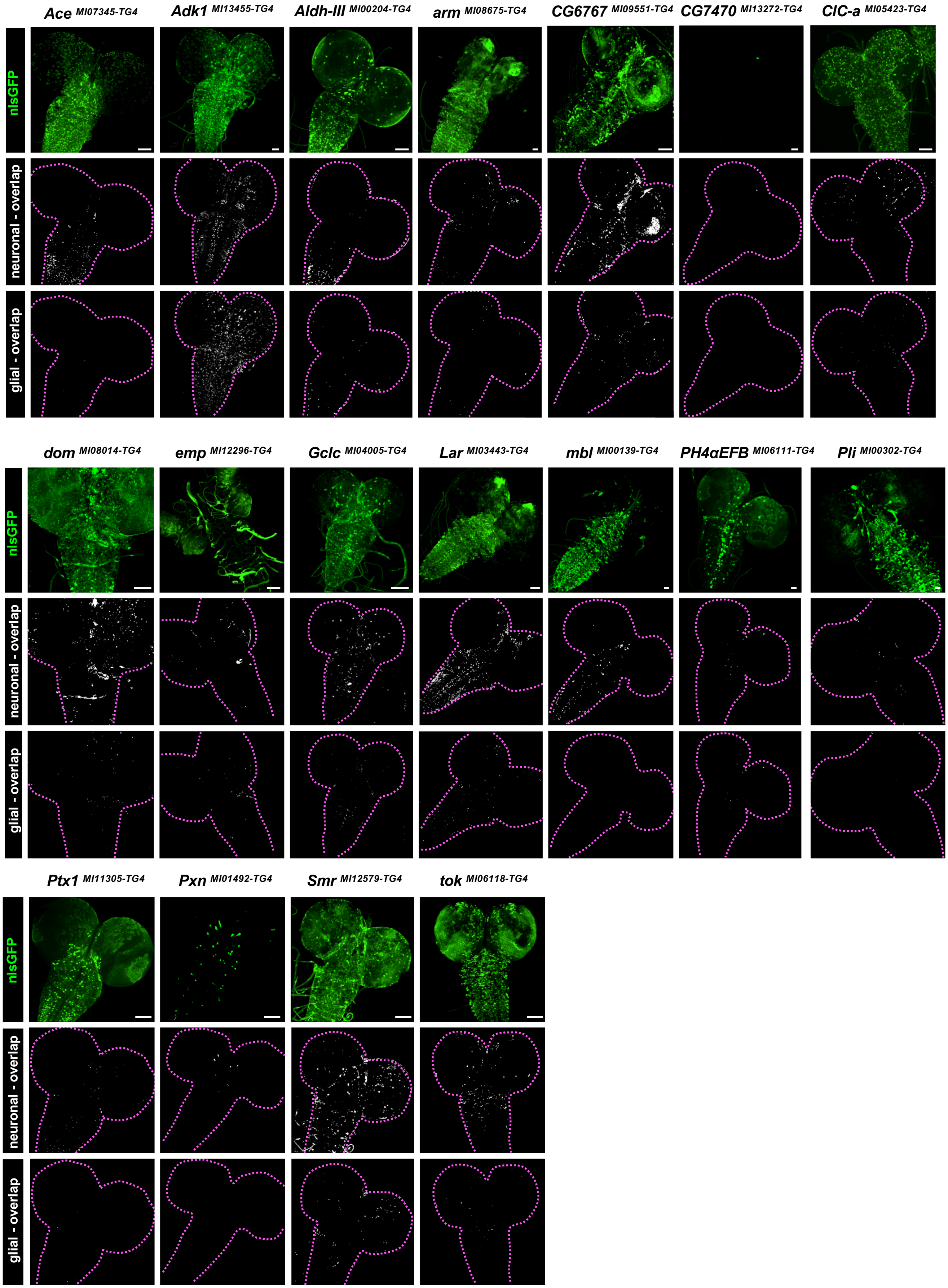

**Supplementary Fig. 10: Additional data related to *GLRA2* variant functional assessments**

**a**, Western blot of 5-day old fly heads expressing *GLRA2* cDNA constructs tagged with a C terminal 3xHA tag expressed using a pan-neuronal driver (*nSyb-GAL4*). **b**, Viability flies expressing *GLRA2* reference or p.T296M at 29°C. **c-f**, Quantification of depolarization and "*ON*"-transient amplitudes from electroretinogram (ERG) recordings of flies expressing GLRA2 reference or variants pan-neuronally (*nSyb-GAL4*) or only in photoreceptors (*Rh1-GAL4*). **g**, GLRA1 crystal structure indicating location of critical homologous GLRA2 amino acid residues relevant to amino acids affected by three patient variants (p.N136, p.T296, p.R252) experimentally tested here. ***p<0.001, ns (not significant).

#### Supplementary Fig. 10:

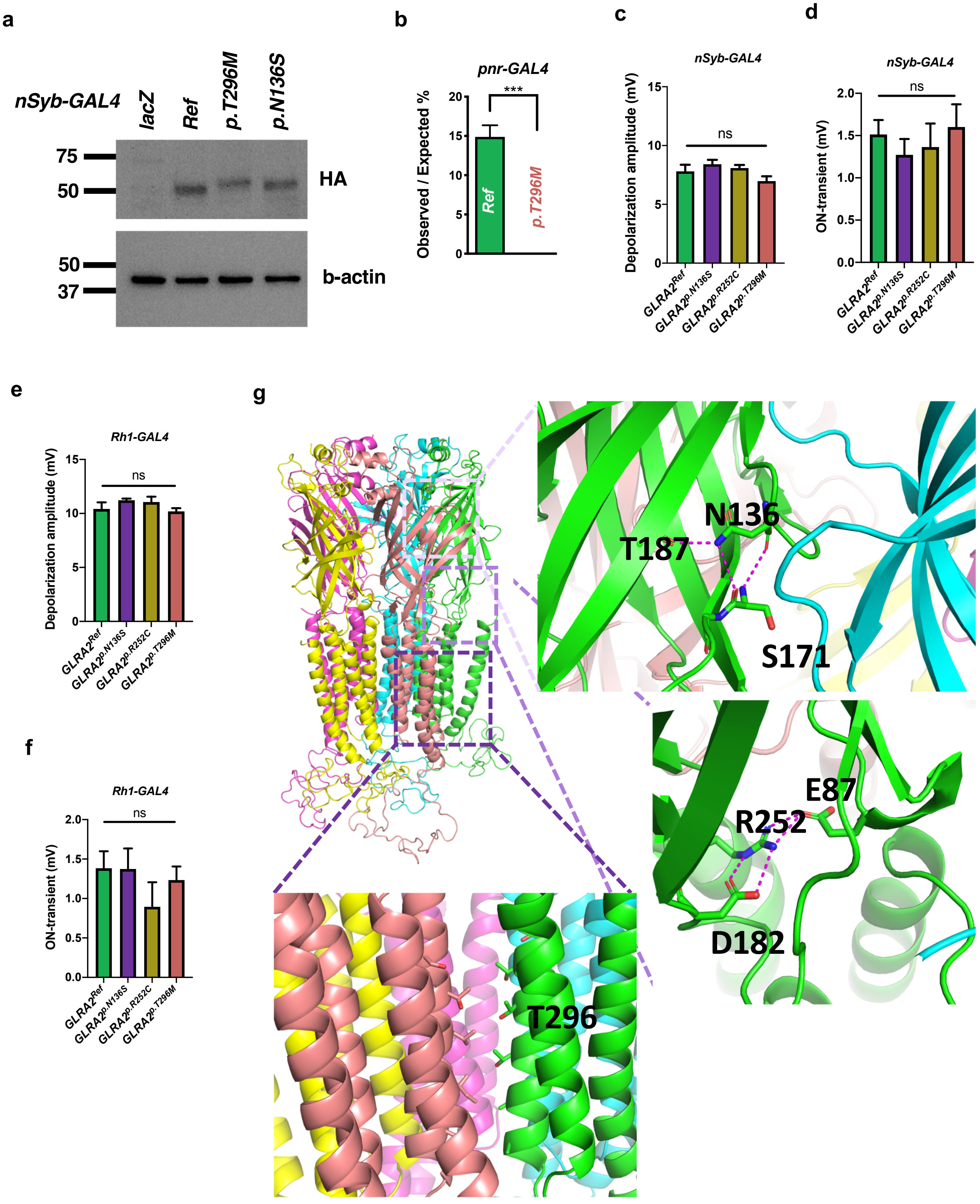
